# One-carbon metabolism sustains effector T cell function through cell-surface glycosylation

**DOI:** 10.64898/2026.10.06.756889

**Authors:** Peng Wang, Amy Yu, Mohammed Mutaher, Almut S. Eisele, Sarah Mouche, Kaitlyn Hung, Reece C. Larson, Madeline Gish, Margaret Zimmer, Jeanette R. Brook, Miriam McNerney, Lea Hammer, Arjun Garg, Efrat Ozeri, Alan Y. L. Wong, Boryana Petrova, Michal Weitman, Ramon C. Sun, Jerome Tamburini, Mathias Wenes, Marc Schwartz, Nir Hacohen, Naama Kanarek

## Abstract

Folate is an essential driver of one-carbon metabolism. The cellular consequences of its deprivation in CD8^+^ T cell and impact on anti-tumor function are unknown. Single cell RNA-seq analysis of primary central nervous system lymphoma (PCNSL) patients’ tumor samples revealed exhaustion in T cells with high folate metabolism gene expression signature. Functional genomics, metabolite profiling, and glycomics identified disrupted cell surface glycosylation as a major consequence of folate deprivation in CD8^+^ T cells. Mechanistically, folate deprivation reduced nucleotide sugar levels, perturbed glycosylation, and triggered the apoptotic unfolded protein response. Overexpressing folate transporters in adoptively transferred CD8^+^ T cells increased intratumoral T cell numbers and reduced exhaustion. Therefore, our data draw a mechanistic line between one-carbon metabolism, cell-surface glycosylation, and anti-tumor T cell function, and support a hypothesis that increased folate uptake can enhance CD8^+^ T cell function including ex vivo and in vivo CAR-T cells.

**Highlights:**

- Although folate metabolism genes are highly expressed in both proliferating tumor and CD8^+^ T cells from primary central nervous system lymphoma patients, the T cells are exhausted while the tumor cells thrive
- Folate is depleted in the TME of MC38-OVA and B16-OVA tumors
- Folate deprivation impairs function and proliferation of CD8^+^ T cells
- Folate is essential for nucleotide sugar metabolism, and its depletion results in perturbation of cell surface glycosylation
- Genetic engineering to increase folate uptake by T cells increases intratumoral T cell numbers and reduces exhaustion

## INTRODUCTION

Throughout their lifespan, T cells encounter a wide variety of metabolic environments. Initial development in the thymus, migration to peripheral lymph nodes and tissues, and systemic circulation throughout the body demands metabolic flexibility for T cells to adapt and carry out their effector functions in diverse environments. In addition to extracellular cues, T cell-intrinsic metabolism is dictated by its cell state and fate (e.g., quiescence, activation, subset type, exhaustion). Thus, T cells encounter many catabolic and anabolic demands during the different stages of their life cycle.

T cell metabolism is highly dynamic and shifts drastically upon T cell activation^1^. Availability of required nutrients and biosynthetic precursors is critical for metabolic pathways that support T cell function and fate^1^. For example, activated T cells require glucose for the rapid induction of aerobic glycolysis to support their rapid proliferation and effector functions^2,3,4^. Amino acids such as serine^5^, arginine^6^, and glutamine^7^ are also critical for T cell proliferation and differentiation, and depletion of these amino acids in the solid tumor microenvironment (TME) contributes to the immunosuppressive milieu^8,9,10^.

One-carbon metabolism (1CM) is a metabolic pathway essential for the biosynthesis of nucleotides^11^, amino acids^12^, and other key metabolites and cofactors whose levels dictate cellular metabolism and function; this includes the methyl donor S-adenosylmethionine (SAM), the antioxidant glutathione (GSH), and the cofactors dihydronicotinamide-adenine dinucleotide phosphate (NADPH) and dihydronicotinamide-adenine dinucleotide (NADH). 1CM is a central metabolic pathway in all cells, but fast proliferating cells heavily rely on 1CM due to their immense demand for *de novo* nucleotide synthesis^13^. 1CM is crucial for many facets of T cell biology: activation^14,15^, proliferation^5,16,17,18,19^, differentiation^20^, survival^15,21^, stemness^22^, and effector functions^16,20,22^. Additionally, boosting 1CM in anti-tumor CD8^+^ T cells by supplementing the one-carbon (1C) unit formate was shown to augment αPD-1 and αPD-L1 checkpoint therapies in mouse cancer models^23,24,25^.

However, 1C units alone are not sufficient to support 1CM because they require a carrier — folate. Folate, also known as the essential vitamin B9, mediates 1CM by functioning as the 1C acceptor and donor, and insufficient folate levels inhibit 1CM^13^. Importantly, there is reason to suspect that folate levels are insufficient to support effector T cell function in cancer patients: plasma folate depletion has been reported in both hematologic^26^ and solid tumor^27,28,29,30,31^ cancer patients. However, it is unknown whether folate is also depleted at the tumor site because the status of folate in the TME has not been previously reported. Many tumors rely on 1CM for their growth, making 1CM a target for cancer therapy (e.g., folate mimetic chemotherapies like methotrexate (MTX) and pemetrexed). For example, primary central nervous system lymphoma (PCNSL), is MTX-responsive^32,33,34,35^, and has poor effector T cell infiltration^36,37,38,39^. It is unknown whether cancer cells’ folate consumption leads to local depletion, and, if so, what potential consequences this may have for T cell function and anti-tumor immunity. To better understand effector T cell’s demand for and utilization of folate, we asked what the functional and metabolic consequences of folate deprivation are in activated effector T cells, and what cellular pathways are the most impacted by this nutritional challenge.

Here we found that although all proliferating cells in PCNSL highly express folate metabolism genes, including both malignant tumor and T cells, the CD8 T cells are exhausted. Based on these findings, we posited that T cell exclusion in these tumors may also be nutritional, including the essential nutrient folate. We demonstrate that folate is depleted in the tumor interstitial fluid of MC38-OVA and B16-OVA tumor mouse models, and show that folate deprivation impaired CD8^+^ T cells function. LC-MS metabolite profiling and an *ex vivo* genome-wide CRISPR knockout screen revealed cell-surface glycosylation as a critically impaired pathway in folate-deprived CD8^+^ T cells, and we confirmed this with N-glycan glycomics. Mechanistically, we show that *de novo* purine synthesis is a bottleneck in nucleotide sugar metabolism, which leads to a global dysregulation of surface proteins glycosylation and activation of the pro-apoptotic unfolded protein response (UPR). We demonstrate that genetic enhancement of folate uptake through overexpression of the folate transporters, SLC19A1 and SLC46A1, in adoptively transferred CD8^+^ T cells increased intratumoral CD8^+^ T cell numbers and activation markers, reduced exhaustion markers, and improved glycosylation profiles. Collectively, these findings demonstrate that folate is critical for proper cell-surface glycosylation in effector T cells, underpinning folate’s role for adequate T cell activation and function, with actionable genetic approaches for T cell functional enhancement in cancer immunity.

## RESULTS

### One-carbon metabolism gene expression is high in proliferating cancer- and T-cells from human primary central nervous system lymphoma (PCNSL), but these T cells are exhausted

PCNSL, which is often diagnosed as diffuse large B-cell lymphoma (DLBCL)^40^, is a deadly cancer with a median survival rate of 3**–**4 years^41^. These tumors are responsive to the antifolate drug, methotrexate^42,43,44,45^, indicating one-carbon metabolism (1CM) reliance. Also, the PCNSL tumor microenvironment (TME) is hostile towards immune cells^38^, with a relatively lower number of activated CD8^+^ T cells compared to other lymphomas^46,47^. The immune status of PCNSL is suggested to be a confounding factor in PCNSL prognosis^48^, underlining the importance of addressing this clinically. The notion that T cell nutritional limitation due to local competition plays a role in reduced tumor immunity is often discussed^49,50^, and we hypothesized that the one-carbon carrier folate can be such a limiting factor, given PCNSL dependency on folate.

To delineate which cells in PCNSL tumors express folate metabolism genes, we analyzed single-cell RNA sequencing (scRNA-seq) data from the publicly available EGAD50000000685 dataset, which profiled tumor samples from 7 PCNSL patients and 2 reactive lymph node samples from patients without cancer as healthy controls (healthy rLN)^51^. From this dataset, we were able to detect malignant B cells (m-B cells) and various other healthy immune cells, including CD8^+^ T cells (**Fig. 1a and Extended Data Fig. 1a–f**). Actively proliferating cells, which were defined by cells in G2/M or S phase (**Extended Data Fig. 1c**) and labeled as “cyc” for cell cycling, were found in both healthy and malignant B cells (h-B cells cyc; m-B cells cyc) (**Fig. 1a**). We observed high variability in the representation of various cell types (**Fig. 1b and Extended Fig. 1b**) between PCNSL and rLN samples. Folate metabolism signature gene expression was high in two cell populations: proliferative B cells, both malignant and healthy (m-B cells cyc and h-B cells cyc) and a subset of CD8^+^ T cells (**Fig. 1c,d and Extended Data Fig. 1g**).

**Fig. 1.**
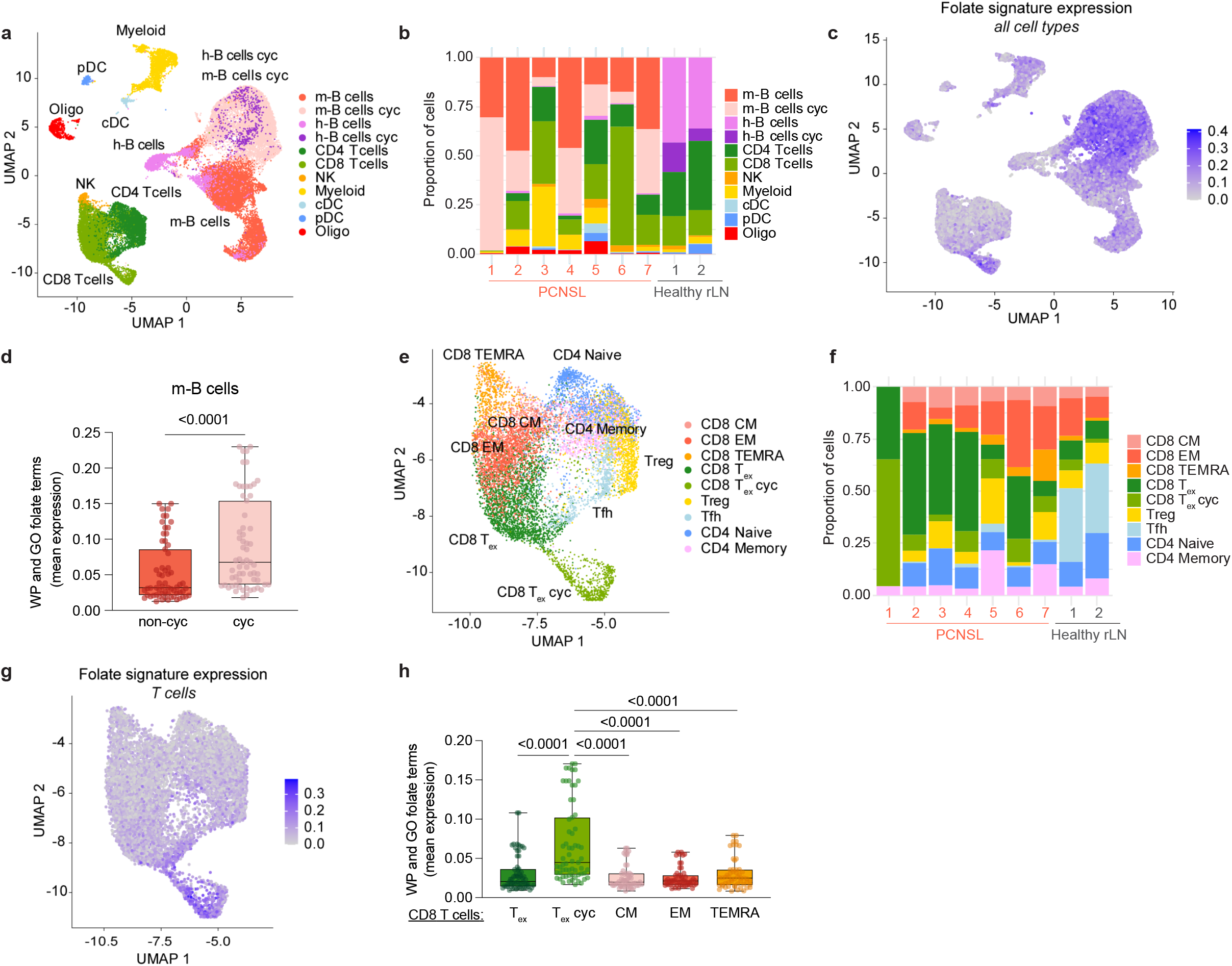
Different fates of cancer and immune cells with high folate metabolism profile in primary central nervous system lymphoma (PCNSL). **a**, Uniform manifold approximation and projection (UMAP) representation of integrated single-cell RNA sequencing (scRNA-seq) data from EGAD50000000685 PCNSL patient brain biopsies (PCNSL; *n* = 7) and reactive lymph node biopsies from patients without cancer (rLN; *n* = 2) annotated by cell type: healthy B cells (h-B cells), malignant B cells (m-B cells), proliferating/cell cycling (cyc), natural killer cells (NK), conventional dendritic cells (cDC), plasmacytoid dendritic cells (pDC), and oligodendrocytes (Oligo). **b**, Relative proportion of cell types in PCNSL and rLN samples. **c**, Folate signature gene expression overlaid on UMAP of all cell types from **(a). d**, Mean expression of each folate-related Biological Processes Gene Ontology (GO) term and WikiPathways (WP) signature across nonproliferating (non-cyc) and proliferating (cyc) m-B cells in PCNSL patients. Full signature names: WP – Disorders of folate metabolism and transport, WP – Folate metabolism, WP – Pathways into methionine and folate-cycles, GO – tetrahydrofolate metabolic process, GO – tetrahydrofolate biosynthetic process, GO – tetrahydrofolate interconversion, GO – 10-formyltetrahydrofolate metabolic process. **e**, UMAP representation of T cells from **(a)**, annotated by T cell subset. CM, central memory; EM, effector memory; TEMRA, terminally differentiated effector memory; CD8 T_ex_, exhausted CD8^+^ T cells; CD8 T_ex_ cyc, exhausted and proliferating/cycling CD8^+^ T cells; Treg, regulatory T cells; Tfh, T follicular helper. **f**, Relative proportion of T cell subsets across PCNSL and rLN samples. **g**, Folate signature gene expression overlaid on UMAP of T cells from **(e)**. **h**, Mean expression of each folate-related GO term and WP signature in CD8^+^ T cell subpopulations in PCNSL samples. Box and whisker plots represent the median and interquartile range (IQR), whiskers represent the data min and max. Dots represent individual signatures in each patient (**d, h**). Statistical significance was assessed using a Mann– Whitney U test (Wilcoxon rank-sum test).

To assess which CD8^+^ T cell subset had elevated folate metabolism gene expression, we further subset all T cells into sub-clusters using the performed of classifiers^52^ (**Fig. 1e,f and Extended Data Fig. 1h**) and analyzed their folate metabolism gene expression levels (**Fig. 1g and Extended Data Fig. 1i**). We could confirm that most T cells in PCNSL are indeed exhausted, with a fraction of those cells actively proliferating (**Fig. 1e,f**). Interestingly, folate metabolism genes were highly expressed in proliferating CD8^+^ T cells expressing exhaustion markers (CD8 T_ex_ cyc); this was significantly elevated compared to non-proliferating, exhausted CD8^+^ T cells (CD8 T_ex_) (**Fig. 1g,h**). We annotated CD8^+^ and CD4^+^ T cell clusters separately (**Extended Data Fig. 1j–m**) and validated that the CD8 T_ex_ cyc population is dominated by the exhausted T cells functional cluster (CD8 T_ex_). We also found that proliferating B cells, regardless of healthy/malignant status or location (i.e., PCNSL tumor vs. healthy rLN), have higher expression of folate metabolism genes (**Extended Data Fig. 1n**). Interestingly, only PCNSL T_ex_ had reduced folate metabolism gene expression compared to healthy rLN, with T_ex_ cyc following a similar but statistically non-significant trend (**Extended Data Fig. 1o**). Together, these results indicate that both actively proliferating malignant B cells and exhausted CD8^+^ T cells have the highest folate metabolism gene expression (**Fig. 1c,g)**. Additionally, exhausted CD8^+^ T cells at the PCNSL tumor site have reduced expression of folate metabolism genes compared to healthy rLN. These results led us to ask whether a competition over folate exist between malignant and immune cells within the TME.

### Folate is a limiting nutrient for T cell function and in the tumor microenvironment

To investigate whether folate (FA) is an essential nutrient for CD8^+^ T cell function, we tested folate’s role in CD8^+^ T cell proliferation and viability using primary activated (day −7, **Extended Data Fig. 2a**) or restimulated (day zero, **Extended Data Fig. 2a**) murine CD8^+^ T cells *ex vivo*. For this, we used two protocols, one that included folate deprivation immediately following primary activation (in blue labels), and one that included folate deprivation two days prior to restimulation (pink labels). In the “blue protocol”, after isolation, murine CD8^+^ T cells were activated and split into FA-replete (2,000 nM; as in RPMI media) and FA-depleted (200, 100, 50, 20, and 0 nM) media. FA-deprived, primary activated CD8^+^ T cells had the most reduced viability 7 days post activation (**Extended Data Fig. 2b**) and showed reduced proliferation at the second day post activation in a dose-dependent manner (**Extended Data Fig. 2c,d**). However, the CD44^+^/CD69^+^ effector population remained high in FA-deprived CD8^+^ T cells instead of dropping as expected prior to restimulation (**Extended Data Fig. 2e,f**), implying a dysregulated activation program and/or selection for more highly CD44^+^/CD69^+^ expressing CD8^+^ T cells in low FA. In the “pink protocol”, CD8^+^ T cells that were FA-deprived 2 days before restimulation (**Extended Data Fig. 2a,** pink labels),and had a later decrease in viability (**Extended Data Fig. 2g**) and better proliferation (**Extended Data Fig. 2h,i**) compared to folate-deprived CD8^+^ T in the “blue protocol”. This is likely due to the lower proliferation rate in this stage of the *ex vivo* restimulation protocol compared to the extremely high rate of proliferation that occurs immediately after primary activation. This led us to focus on the “pink protocol”, where folate deprivation is applied later (pink labels in **Extended Data Fig. 2a**). However, similar to the “blue protocol”, the CD44^+^/CD69^+^ population was increased in the 0 nM FA condition by day 3 post-restimulation also in the “pink protocol” (**Extended Data Fig. 2j**). These results validate folate’s role in CD8^+^ T cell viability, proliferation and activation, and reveal FA depletion-induced dysregulation of T cell activation. This work informed our protocol moving forward: first, to perform the “pink protocol”, where FA depletion is done prior to restimulation and not around the primary activation, as in the “blue protocol” in order to allow the cells to pass the early high proliferation stage with sufficient folate; second, to drop FA in the media to zero in order to avoid intermittent folate depletion due to the recyclable nature of intracellular folate, which is diluted by cell division and retained in cells even when extracellular levels are low^53^, and which would require multiple days of culture in FA-deprived media for intracellular folate stores to be depleted^54,55,56^.

Next, we tested whether folate deprivation impairs CD8^+^ T cell antigen-specific cytotoxic capacity. We performed an *ex vivo* killing assay using FA-deprived OT-I cells and B16-OVA GFP expressing cells. Surprisingly, we did not observe a dose-dependent decrease in killing capacity; all doses of FA deprivation we tested (200, 100, 50, 20, and 0 nM) led to significantly reduced killing capacity compared to the FA-replete control (2,000 nM) (**Fig. 2a**). This can be the result of the drastic drop from RPMI-levels of folate (2,000 nM) and reflect a limitation of our *ex vivo* system.

**Fig. 2.**
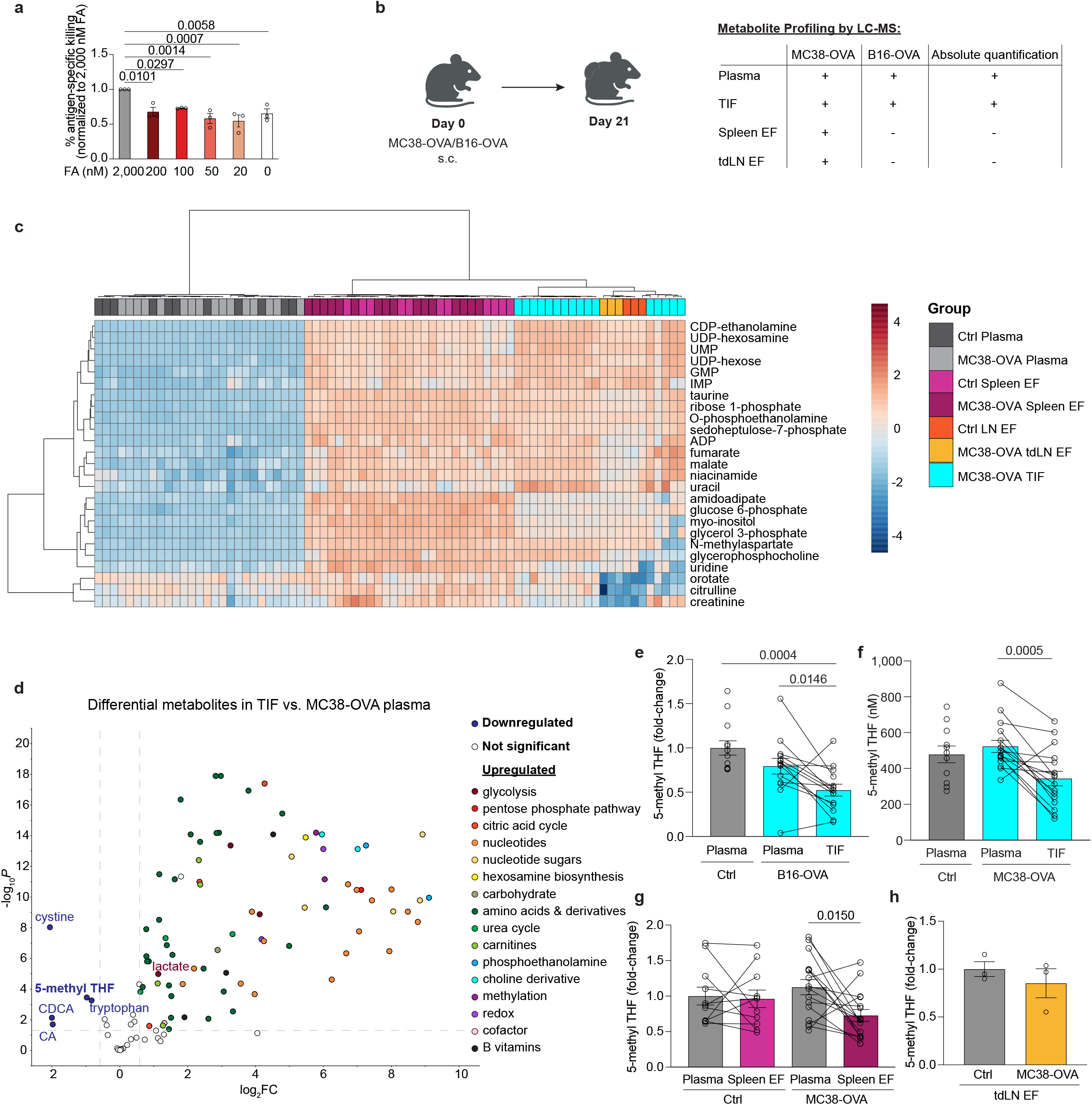
Folate is a limiting nutrient in the tumor microenvironment. **a**, Fold change in percentage of antigen-specific B16-OVA killing *ex vivo* by activated, folate-deprived OT-I cells. Values are relative to 2,000 nM folic acid (FA; folate) (*n* = 3 biological replicates). **b,** Experimental schematic for MC38-OVA and B16-OVA tumor cells administration and downstream LC-MS analysis in C57BL6/J mice. The table indicates the measurements performed with each sample: plasma, tumor interstitial fluid (TIF), spleen extracellular fluid (EF) and tumor-draining lymph node extracellular fluid (tdLN EF). **c,** Heatmap of top 25 metabolites in plasma, TIF, spleen EF, and tdLN EF in control (*n* = 11) vs. MC38-OVA mice (*n* = 16). LN EF samples represent 3 biological replicates of pooled mice (4-5 mice/sample). **d,** Volcano plot depicting metabolite changes between TIF and plasma in MC38-OVA mice. Downregulated metabolites are colored dark blue and insignificant metabolites are colored light grey. Upregulated metabolites are colored by metabolic pathway. **e,** 5-methyl tetrahydrofolate (THF) (circulating form of folate; nM) fold change values (relative to control plasma) in plasma and TIF of control (*n* = 13) and B16-OVA mice (*n* = 14). *n* = 3, biological replicates. **f,** 5-methyl THF levels in plasma and TIF of control (*n* = 11) and MC38-OVA mice (*n* = 16). 5-methyl THF concentrations were calculated using ^13^C-labelled 5-methyl THF standard curves in plasma and TIF. **g,** 5-methyl THF fold change values (relative to control plasma) in plasma and spleen EF of control (*n* = 11) and MC38-OVA mice (*n* = 16). **h,** 5-methyl THF fold change values (relative to control LN EF) in tdLN EF of control and MC38-OVA mice (*n* = 3 pooled samples, 4-5 mice/sample). All data in **a, e–h** represent mean with s.e.m. Statistical significance was determined by one-way analysis of variance (ANOVA) with Dunnett’s multiple comparisons test (**a**), one-way ANOVA with Tukey’s post-hoc test for multiple comparisons (**e**, **f**), paired, two-tailed parametric *t*-test (**e–g**), or a two-tailed Student’s *t*-test (**h**).

We next asked how consistent folate levels are *in vivo*, and whether T cells experience a drop in folate levels when they move from the blood stream to the TME, or during other migration events involving lymphoid organs such as the spleen and the lymph nodes. To address this, we focused on the circulating form of folate, 5-methyl tetrahydrofolate (5-methyl THF). We utilized immunogenic murine melanoma (B16-OVA) and colon cancer (MC38-OVA) models expressing the model antigen ovalbumin (OVA)^57,58^. 21 days post-MC38-OVA inoculation, plasma, tumor interstitial fluid (TIF), spleen extracellular fluid (EF)^59,60,61^, and tumor-draining lymph nodes (tdLNs) EF^62,63^ were collected for metabolite analysis by LC-MS^1^ (**Fig. 2b,c)**. Hierarchical clustering of the top 25 differentially abundant metabolites in MC38-OVA tumor-bearing mouse plasma, spleen EF, tdLN EF, and in TIF compared to control mice indicated that the natural grouping of these metabolites was dominated by different biological matrices, and not tumor presence (**Fig. 2c)**.

Comparing TIF to plasma of tumor-bearing mice in both the B16-OVA and MC38-OVA models (**Extended Data Fig. 3a,e,i**) revealed that most metabolites are elevated in the TIF, with a few exceptions, and one of them was indeed 5-methyl THF (**Fig. 2d**).

Upregulated metabolites included various amino acids and metabolites related to glycolysis and the citric acid cycle; these are intracellular metabolites, not expected in the extracellular space, and may originate from spillage of dying cells at the tumor site. Strikingly, only a few metabolites were significantly lower in TIF: tryptophan, cystine, chenodeoxycholic acid (CDCA), cholic acid (CA), and 5-methyl THF. Aligned with previous reports, lactate was increased in the TIF (**Extended Data Fig. 3b,f,j**) and tryptophan was decreased in the TIF compared to plasma from both control and tumor-bearing mice (**Extended Data Fig. 3c,g,k**). Surprisingly, levels of inosine, which has been described to augment chimeric antigen T cell (CAR-T) function and stemness *in vitro*^64^ and enhance both ICB and CAR-T therapy in B16-F10 tumors^65^ was significantly increased in the TIF relative to plasma (**Extended Data Fig. 3d,h,l**). This raises the possibility that tumor-infiltrating T cells may inefficiently take up inosine or may have limited inosine-dependent metabolism, despite high inosine levels in the local environment.

To strengthen our finding of depleted 5-methyl THF in the TIF, we performed 5-methyl THF relative quantification in B16-OVA, and absolute quantification in MC38-OVA in paired plasma and TIF. Indeed, this revealed that 5-methyl THF was significantly depleted in the TIF compared to paired tumor-bearing mouse plasma (**Fig. 2e,f and Extended Data Fig. 3m)**.

Unlike folate, other B vitamins were not depleted in the TIF. Niacinamide (vitamin B3) and pantothenate (vitamin B5) were significantly increased in the TIF compared to plasma (**Extended Data Fig. 3n,o**), highlighting 5-methyl THF (vitamin B9) as a uniquely depleted B vitamin in the TME.

Our absolute quantification of 5-methyl THF indicated that the median 5-methyl THF level in control C57BL/6J mouse plasma was 485.5 nM (range: 273.9 – 742.2 nM) (**Fig. 2f**), which is higher than previously reported^66,67^. Further, our measurements show no depletion of folate in the plasma of tumor-bearing mice, that is not consistent with human gastric cancer patient data, that reported systemic folate depletion in gastric cancer patients^28^. We hypothesize that this is due to the excess amounts of FA (2 mg/kg) supplemented in standard mouse chow, leading to supraphysiological 5-methyl THF levels in mouse plasma^68^. Yet, even in this high-folate condition, 5-methyl THF levels dropped in TIF compared to paired plasma samples, with a median concentration of 333.3 nM (range: 120.3 – 660.6 nM) (**Fig. 2f and Extended Data Fig. 3m)**.

5-methyl THF was mildly reduced in the spleen EF of tumor-bearing mice compared to paired plasma samples (**Fig. 2g**), and not different between control LN EF and MC38-OVA tdLN EF (**Fig. 2h**). Due to volume limitations, ^13^C-labelled 5-methyl THF standard curves could not be generated in the spleen and tdLN EF, and thus only relative quantification could be calculated.

In summary, we conclude that folate is one of a handful of metabolites that are depleted in the relatively rich TIF. We are hesitant to conclude that this is not the case for plasma and tumor-draining lymph nodes as a limitation of our experiment is the supraphysiological levels of folate in standard mouse chow; this is reflected in the 5-methyl THF concentrations measured in the plasma of the mice in our lab.

### Genome-wide *ex vivo* CRISPR screen reveals an essential role for folate in nucleotide sugar and glycolipid biosynthesis in CD8^+^ T cells

Folate is responsible for several major metabolic pathways and has been found to play a role in various aspects of T cell biology^5,14,15,16,17,18,19,20,21,22,23,24,25^. We therefore asked, what are the specific cellular limitations of FA-deprived CD8^+^ T cells which compromise T cell fitness? To address this, we performed an *ex vivo* genome-wide CRISPR screen in normal (2,000 nM, RPMI level) and low FA conditions (50 nM) (**Fig. 3a**). The CRISPR screen was performed in 50 nM folate because the screening protocol requires 7 days in culture, and our CFSE data (**Extended Data Fig. 2h,i**) indicated reduced proliferation as early as the day of restimulation (day 0, **Extended Data Fig. 2i**) in CD8^+^ T cells cultured in 0 nM FA. We considered this to be a highly premature bottleneck and chose to perform the screen in 50 nM FA, which was very similar to 0 nM but did not exhibit this early proliferation inhibition.

**Fig. 3.**
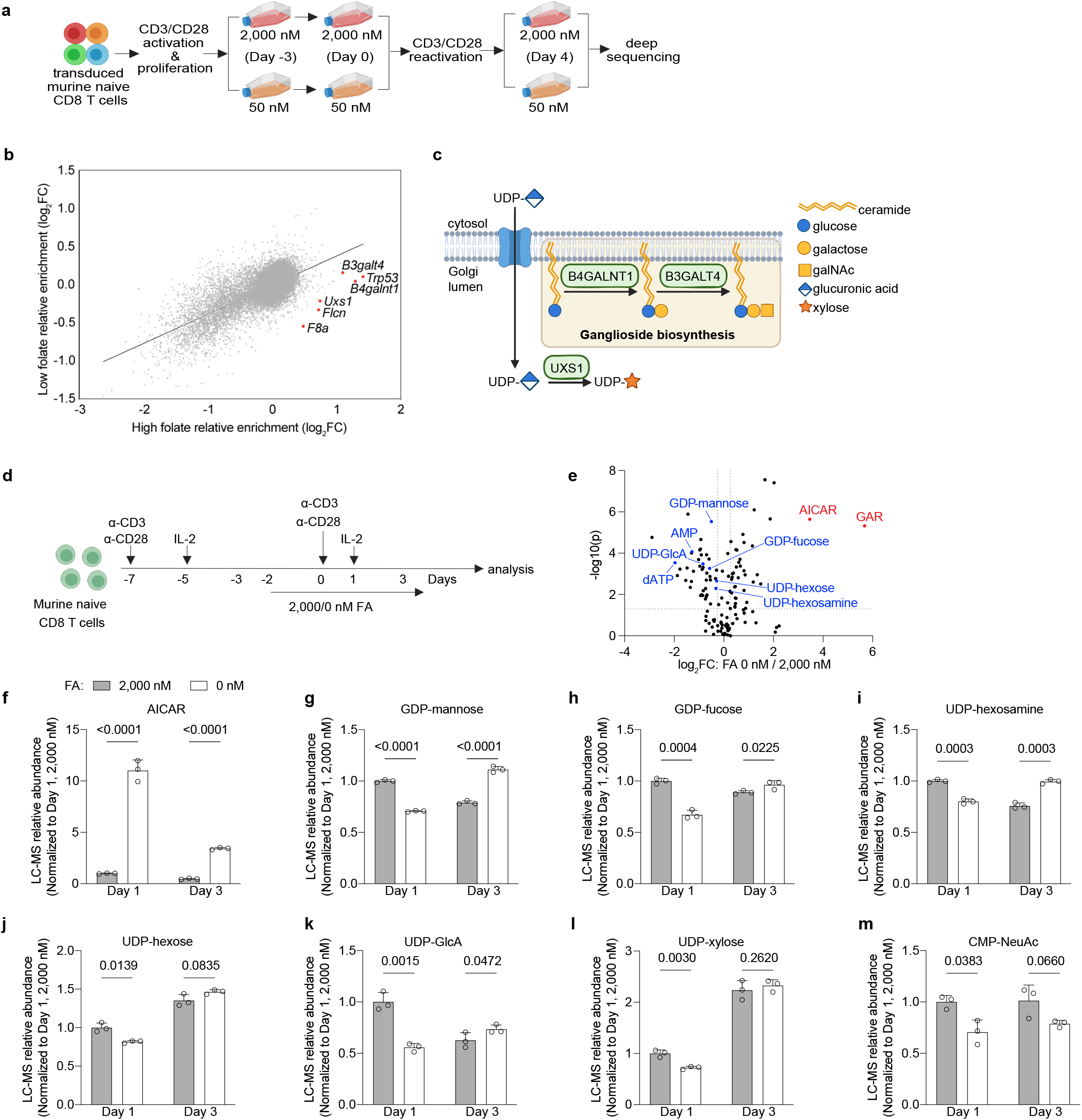
Folate-deprived CD8^+^ T cells suffer from perturbed nucleotide sugar metabolism. **a,** Experimental schematic of e*x vivo* genome-wide CRISPR screen using mouse naïve CD8^+^ T cells. Isolated murine naïve CD8^+^ T cells were transduced with the Brie sgRNA library^117^ and restimulated as before. The successfully transduced cells were selected and then cultured in folate-replete (2,000 nM) and folate-depleted (50 nM) conditions for three days prior to restimulation, then allowed to proliferate for four more days before harvesting for deep sequencing of sgRNA abundance. **b**, Scatter plot depicting the gRNA enrichment in 2,000 nM (x-axis) and 50 nM (y-axis) folic acid (FA; folate) FA. The top 6 differentially depleted guides in 50 nM FA (ranked by log_2_FC 50 nM – log_2_FC 2,000 nM) are represented by red dots. *B3galt4*: beta-1,3-galactosyltransferase 4; *Trp53*: tumor protein P53; *B4galnt1*: beta-1,4-N-acetyl-galactosaminyltransferase 1; *Uxs1*: UDP-glucuronate decarboxylase 1; *Flcn*: folliculin; *F8a*: coagulation factor VIII associated. **c**, Schematic of the top 3 glycosylation related hits’ (*B4galnt1, B3galt4, Uxs1*) function in ganglioside biosynthesis and nucleotide sugar metabolism in the Golgi lumen. **d,** Schematic of *ex vivo* restimulation assay with FA deprivation. **e,** Volcano plot visualizing differentially regulated metabolites (ranked by *t*-test) between control (2,000 nM) and FA-deprived (0 nM) CD8^+^ T cells 1 day after restimulation. Significantly downregulated nucleotide sugars in the FA-deprived group are highlighted in blue, while upregulated purine intermediates (GAR, AICAR) are highlighted in red. **f–m,** Relative abundance of purine and pyrimidine nucleotide sugar levels in control (2,000 nM) vs. FA-deprived T cells (0 nM) at days 1 and 3 after restimulation (*n* = 3 biological replicates). **f,** 5-aminoimidazole-4-carboxamide ribonucleoside (AICAR), a purine intermediate; **g, h**, Guanosine-based purine nucleotide sugars. **g,** GDP-mannose. **h,** GDP-fucose. **i–l,** Uridine-based pyrimidine nucleotide sugars. **i,** UDP-hexosamine (UDP-N-acetylglucosamine/UDP-N-acetylgalactosamine, also known as UDP-GlcNAc/UDP-GalNAc). **j,** UDP-hexose (UDP-glucose/UDP-galactose). **k,** UDP-glucuronic acid (UDP-GlcA). **l,** UDP-xylose. **m,** Cytidine-based pyrimidine nucleotide sugar, CMP-N-acetylneuraminic acid (CMP-NeuAc); precursor to CMP-sialic acid. All data in **f–m** represent mean and s.e.m. Statistical significance was determined by repeated-measures two-way ANOVA with Šídák’s multiple comparisons test.

Among the top differentially essential genes between the FA-replete and FA-depleted conditions were three genes encoding glycosylation enzymes: the nucleotide sugar metabolism enzyme, UDP-glucuronate decarboxylase 1 (UXS1), and the ganglioside biosynthesis enzymes beta-1,4-N-acetyl-galactosaminyltransferase 1 (B4GALNT1) and beta-1,3-galactosyltransferase 4 (B3GALT4) (**Fig. 3b**). UXS1 is responsible for the conversion of UDP-glucuronic acid (UDP-GlcA) to UDP-xylose, which is used to initiate glycosaminoglycan synthesis on proteoglycans, while B4GALNT1 and B3GALT4 catalyze linear steps in ganglioside biosynthesis (**Fig. 3c**). Gangliosides are sialylated glycosphingolipids predominantly found on the outer leaflet of cell membranes, serving as functional components in lipid rafts and playing important roles in cell-cell interaction and signal transduction. Consequently, gangliosides are critical for immune cell activation and modulating the tumor-immune microenvironment^69,70,71,72,73^. B4GALNT1 catalyzes the transfer of GalNAc onto GM3 or GD3 to create GM2 or GD2, respectively. Then, B3GALT4 catalyzes the transfer of a galactose to GM2 or GD2, generating GM1 or GD1b, respectively (**Fig. 3c**). While *Uxs1* single guide RNA (sgRNA) abundance was enriched in high FA, it was mildly depleted in low FA conditions (**Fig. 3b**). This suggests that *Uxs1* loss was detrimental to survival and proliferation in low FA but beneficial in FA-replete conditions. Similarly, *B4galnt1* and *B3galt4* sgRNA abundance was enriched in both high and low FA conditions, but to a lesser degree in low FA. This suggests that reduced ganglioside biosynthesis conferred a survival advantage for CD8^+^ T cells, but less so in low FA conditions (**Fig. 3b**).

To validate our CRISPR screen results, we performed a competitive growth assay in FA-replete and FA-depleted conditions using control GFP^+^ CD8^+^ T cells and single gene KO (Δ) GFP^-^ CD8^+^ T cells, which were generated by sgRNA nucleofection (**Extended Data Fig. 4a and Supplementary Table 1**). Consistent with our screen results, *B4galnt1*Δ, *B3galt4*Δ, and *Uxs1*Δ CD8^+^ T cells were enriched compared to intergenic control in FA-replete media (**Extended Data Fig. 4b**). Additionally, these growth benefits were reduced (for *B4galnt1*Δ and *B3galt4*Δ) or eliminated (for *Uxs1*Δ) in the FA-deprived conditions (50 nM), confirming our screen results. We also profiled levels of the glycolipid, asialo-GM1 (ganglio-N-tetraosylceramide; GgOse4Cer), using flow cytometry on our Δ CD8^+^ T cells to verify whether knockout of these genes impaired ganglioside cell surface levels. Indeed, asialo-GM1 levels were decreased in *B4galnt1*Δ and *B3galt4*Δ CD8^+^ T cells, confirming successful KO of these genes (**Extended Data Fig. 4c**). Notably, *Uxs1*Δ CD8^+^ T cells also had reduced cell surface asialo-GM1 levels. Since UXS1 is the key glycosyltransferase for UDP-xylose synthesis, which is the initiating sugar for glycosaminoglycan synthesis on peptidoglycans, it is possible that peptidoglycan defects may potentially impact ganglioside homeostasis on the cell surface. We also observed reduced asialo-GM1 levels in WT FA-deprived cells. This demonstrates that folate deprivation alone is sufficient to drive a reduction in asialo-GM1 cell-surface levels, independently of the genetic perturbations we introduced; this may be due to perturbation of a folate-dependent metabolic pathway.

To explore the metabolic consequences of folate deprivation in primary, restimulated CD8^+^ T cells, we performed metabolite profiling on FA-replete (2,000 nM) and FA-deprived (0 nM) murine splenic CD8^+^ T cells (**Fig. 3d**). After initial activation with αCD3/αCD28 antibodies, activated CD8^+^ T cells were FA-deprived 48 hours prior to restimulation and were collected at days 1 and 3 post-restimulation for metabolite profiling. As expected^74,75^, we observed 1CM-related perturbations in FA-deprived CD8^+^ T cells. Purine metabolism intermediates 5-aminoimidazole-4-carboxamide ribonucleoside (AICAR) and glycinamide ribonucleotide (GAR) were increased in FA-deprived CD8^+^ T cells, while downstream purine nucleotides like IMP, AMP and dATP were decreased (**Fig. 3e,f and Extended Data Fig. 4d,e**). Other nucleotides were not perturbed (**Extended Data Fig. 4f–i**).

Beyond stand-alone nucleotides, we also observed a drop in the levels of some nucleotide sugars in the FA-deprived CD8^+^ T cells on day 1, sometimes followed by an increase by day 3 (**Fig. 3e**, highlighted in blue, and plots of the nine most abundant mammalian nucleotide sugars in **Fig. 3g–m**). Nucleotide sugars are the essential glycosyl donors for glycosylation reactions in the cell, and require a nucleotide and monosaccharide for their biosynthesis. These can be subdivided into two categories: purine nucleotide sugars (GDP-mannose, GDP-fucose) and pyrimidine nucleotide sugars (UDP-hexosamine [UDP-GlcNAc and UDP-GalNAc], UDP-hexose [UDP-glucose and UDP-galactose], UDP-GlcA, UDP-xylose, and CMP-NeuAc). With our LC-MS method, we were unable to distinguish between the epimer forms for both UDP-hexosamine (UDP-GlcNAc and UDP-GalNAc) and UDP-hexose (UDP-glucose and UDP-galactose). Based on folate’s critical role in *de novo* purine and pyrimidine synthesis and the top differential metabolites we observed in FA-deprived CD8^+^ T cells, we rationalized that folate deprivation may perturb nucleotide sugar biosynthesis.

We noted, that although the pyrimidines themselves were not significantly perturbed by folate deprivation (**Extended Data Fig. 4g–i**), the pyrimidine-sugars we analyzed showed a similar pattern to purine-sugars with a drop by day 1 followed by an increase in day 3 (**Fig. 3g– m**), except for CMP-NeuAc (**Fig. 3m**), that remained low in day 3. In summary, metabolite profiling by LC-MS revealed that FA-deprived CD8^+^ T cells manifest reduced nucleotide sugar levels at day 1 post-restimulation.

### Purine synthesis bottleneck leads to perturbed nucleotide sugar metabolism in folate-deprived CD8^+^ T cells

To distinguish between perturbed *de novo* synthesis of nucleotide sugars in FA-deprived CD8^+^ T cells and other potential causes of reduced nucleotide sugar levels, we performed stable-isotope tracing using ^15^N-labelled L-glutamine (amide-^15^N) in FA-replete and FA-deprived CD8^+^ T cells after restimulation. The labelled amide-^15^N from glutamine is incorporated into purine and pyrimidine *de novo* synthesis at multiple steps through the deamination of glutamine, allowing tracing of the ^15^N label into purine and pyrimidine nucleotides and their downstream purine and pyrimidine nucleotide sugars (**Fig. 4a**). ^15^N-labelled L-glutamine was added on days 1 or 3 for 4 hours to label *de novo* synthesized metabolites. Both purine nucleotide sugars, GDP-mannose and GDP-fucose, had significantly reduced labelling in FA-deprived CD8^+^ T cells on both days 1 and 3 post-restimulation (**Fig. 4b and Extended Data Fig. 5a**), indicating reduced *de novo* synthesis. As observed previously (**Fig. 3g,h**), total levels of GDP-mannose and GDP-fucose were lower on day 1 but not on day 3 post-restimulation (**Fig. 4c and Extended Data Fig. 5b**). Other purine metabolites like IMP and GTP followed a similar trend (**Extended Data Fig. 5c–f**). The observed recovery of total purine nucleotide and nucleotide sugar levels by day 3 post-restimulation, despite decreased *de novo* synthesis, may potentially be due to reduced utilization of these metabolites. *De novo* synthesis of the pyrimidine nucleotide sugars UDP-hexosamine, UDP-hexose, UDP-GlcA, and UDP-xylose was mildly reduced in FA-deprived CD8^+^ T cells on day 1 post-restimulation, and recovered by day 3 post-restimulation (**Fig. 4d,e and Extended Data Fig. 5g–l**). *De novo* synthesis of UTP, the precursor nucleotide for UDP-sugars, showed a similar trend, while total levels of UTP were unchanged in FA-deprived CD8^+^ T cells (**Extended Data Fig. 5m,n**). Overall, these results suggest that the purine synthesis pathway serves as the major metabolic bottleneck in FA-deprived CD8^+^ T cells, resulting in depleted purine nucleotide sugar levels.

**Fig. 4.**
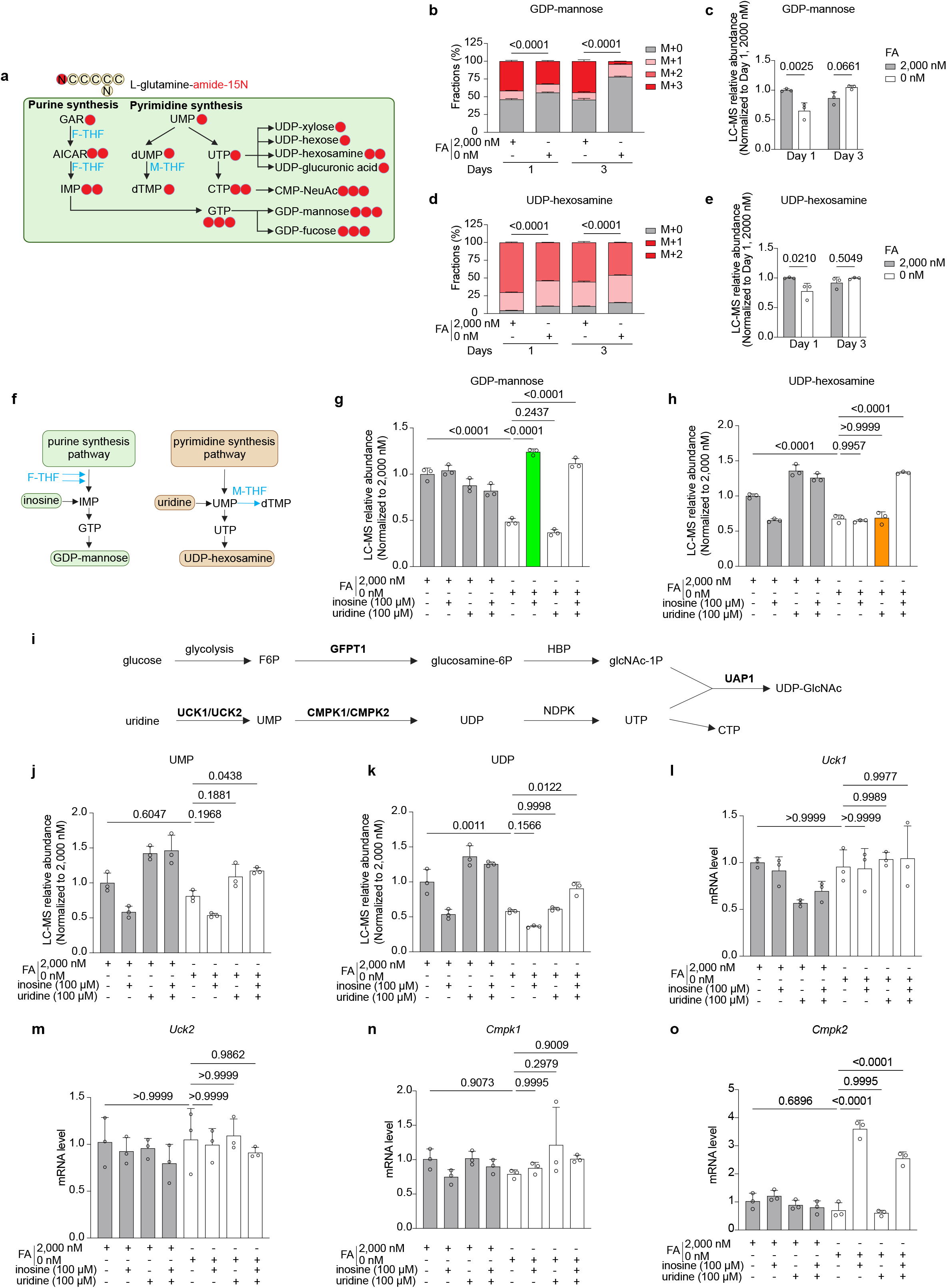
Purine synthesis is a bottleneck for impaired purine and pyrimidine nucleotide sugar metabolism in folate-deprived CD8^+^ T cells. **a,** Schematic of amide-^15^N L-glutamine stable isotope tracing for *de novo* synthesis of purine and pyrimidine nucleotides and downstream nucleotide sugars. Red dots indicate ^15^N atoms incorporated into downstream metabolites through glutamine deamination. Folate-dependent steps are marked by the required folate forms (in blue text): 10-formyl THF (F-THF) and 5,10-methylene THF (M-THF). **b–e**, ^15^N labelling in GDP-mannose (**b**, **c**) and UDP-hexosamine (**d**, **e**) in control (2,000 nM) vs. FA-deprived CD8^+^ T cells (0 nM) at days 1 and 3 after restimulation. For labelling, M+0 represents the endogenous unlabelled metabolite (grey bars). *De novo* synthesis is represented by incorporation of the stable isotope, indicated by M+1, M+2, and M+3 (red bars). Data is shown both as the fraction of labeled metabolites (**b**, **d**) and the relative abundance (**c, e**). **f,** Schematic illustration of purine and pyrimidine nucleotide/nucleotide sugar biosynthesis, with supplemented inosine and uridine metabolic integration indicated. Folate-dependent steps are marked by the required folate forms (in blue text): 10-formyl THF (F-THF) and 5, 10-methylene THF (M-THF). **g, h,** Relative abundance of GDP-mannose (**g**) and UDP-hexosamine (**h**) levels in FA-deprived CD8^+^ T cells supplemented with inosine (100 µM), uridine (100 µM), or both. Grey bars represent T cells in 2,000 nM FA while white bars correspond to 0 nM FA. The green bar represents inosine supplementation in 0 nM FA, while the orange bar represents uridine supplementation in 0 nM FA. **i,** Schematic illustration of UDP-GlcNAc biosynthesis including the hexosamine biosynthetic pathway (HBP), which synthesizes GlcNAc-1P (N-Acetyl-D-glucosamine 1-phosphate), and the uridine salvage pathway, which synthesizes UTP. **j, k,** Relative abundance of UMP (**j**) and UDP (**k**) in FA-deprived CD8^+^ T cells supplemented with inosine (100 µM), uridine (100 µM), or both. **l, o,** mRNA levels of pyrimidine biosynthesis enzymes in FA-deprived CD8^+^ T cells supplemented with inosine (100 µM), uridine (100 µM), or both. **l,** Uridine-cytidine kinase like-1 (UCK1). **m**, Uridine-cytidine kinase like-2 (UCK2). **n,** Cytosine/uridine monophosphate kinase 1 (CMPK1). **o,** Cytosine/uridine monophosphate kinase 2 (CMPK2). All data in **b–e**, **g**, **h**, **j–o** represent 3 biological replicates and are presented as mean and s.e.m. *P* values were determined by a repeated-measures two-way ANOVA. To evaluate the treatment effect at each time point, Šídák’s multiple comparisons test was used (**b–e**, with **b** and **d** comparing the M+0 fraction of each condition), or one-way ANOVA with Tukey’s post-hoc test for multiple comparisons (**g**, **h**, **j–o**).

The mild changes we observed in pyrimidine sugars are not expected, because folate is not required for the synthesis of UTP and CTP (only for dTMP) (**Fig. 4f**). To better understand the limiting steps in purine-sugar and pyrimidine-sugar biosynthesis in FA-deprived CD8^+^ T cells, we investigated whether boosting the purine and pyrimidine salvage pathways could rescue the perturbed nucleotide sugar levels. This can be done by supplementation of the purine and pyrimidine nucleosides, inosine and uridine, respectively (**Fig. 4f**). Inosine is converted to IMP, bypassing the folate requirement for *de novo* purine synthesis, while uridine is converted to UMP, boosting the folate-independent pyrimidine biosynthesis pathway (**Fig. 4f**).

We observed that inosine, but not uridine, was able to rescue most of the differentially abundant metabolites induced by folate deprivation (**Extended Data Fig. 6a–d**). Supplementation with inosine alone was sufficient to rescue GDP-mannose levels in FA-deprived CD8^+^ T cells (**Fig. 4g**). Similarly, other purine metabolites like GDP-fucose, GMP, and GDP were also rescued by inosine alone (**Extended Data Fig. 6e–g**). In contrast, uridine supplementation alone failed to restore UDP-hexosamine levels in FA-deprived CD8^+^ T cells; however, the combination of both inosine and uridine supplementation was able to rescue UDP-hexosamine levels (**Fig. 4h**). This was also generally observed for other pyrimidine nucleotide sugars, except for UDP-xylose (**Extended Data Fig. 6h–j**). These results indicate that sufficient inosine is required for uridine to rescue the pyrimidine-sugar synthesis in folate-deprived CD8^+^ T cells. This implies that a crosstalk between inosine and the pyrimidine-sugar synthesis might exist and regulate pyrimidine-sugar synthesis.

To further investigate this potential inosine-pyrimidine-sugar crosstalk, we asked which step might be regulated by inosine and focused on UDP-GlcNAc synthesis as an example. UDP-GlcNAc synthesis requires contributions from both the pyrimidine nucleotide synthesis and hexosamine biosynthesis pathways (HBP) (**Fig. 4i**). Our metabolite profiling data did not point to perturbed sugar synthesis (**Supplementary Table 2**), so we focused on the UTP synthesis arm. In our uridine supplementation experiments, we provided uridine to the cells, that can be detected intracellularly at the expected levels (**Extended Data Fig. 6k**). Similarly, the subsequent metabolite in the pathway, UMP, was also elevated following uridine supplementation - as expected (**Fig. 4j**). However, UDP, the next downstream metabolite, was not increased in FA-deprived and uridine-supplemented CD8^+^ T cells (**Fig. 4k**). Notably, supplementation of uridine and inosine together did result in elevated UMP and UDP levels, as well as elevated levels of most downstream pyrimidine metabolites (**Fig. 4j,k and Extended Data Fig. 6h–j,l**). This indicates that the potential purine-dependent bottleneck lies between the UMP to UDP conversion step, which is mediated by two cytidine/uridine monophosphate kinases (CMPKs): CMPK1 and CMPK2 (**Fig. 4i**). Subsequently, we asked whether inosine levels regulate one of these enzymes, allowing the supplemented uridine to be metabolized to UDP and further downstream pyrimidine metabolites. Indeed, CMPK2 transcript levels were increased three-fold when inosine was supplemented to FA-deprived CD8^+^ T cells, while other key enzymes in the UDP-GlcNAc synthesis pathway (e.g., UCK1, UCK2, CMPK1, UAP1, and GFPT1) did not display an inosine-dependent transcriptional upregulation (**Fig. 4l–o and Extended Data Fig. 6m,n**). Altogether, these data corroborate the key role folate deprivation has in perturbing purine nucleotide synthesis and subsequently both purine and pyrimidine nucleotide sugar depletion. This work revealed that sugar nucleotide synthesis is perturbed in FA-deprived CD8^+^ T cells through both direct inhibition of folate-dependent purine synthesis, and indirect, purine-dependent transcriptionally regulated inhibition of pyrimidine sugars.

### Folate deprivation compromises functionally essential cell surface glycosylation

Nucleotide sugar perturbations in FA-deprived CD8^+^ T cells likely disturb subsequent glycosylation processes, which may result in globally dysregulated glycosylation. Nucleotide sugars are the required activated sugar donors in glycosylation reactions, serving as the building blocks for glycans that can subsequently be added to modify glycoconjugates (e.g., glycoproteins and glycolipids). These cell surface glycoconjugates form and dictate the complexity of the glycocalyx, an elaborate structure that is critical for mediating immune functions^76^ (**Fig. 5a**). While we recognize that nucleotide sugars are essential for both N-glycosylation and O-glycosylation, we focused here on the N-glycosylation profile of FA-deprived CD8^+^ T cells using matrix-assisted laser desorption/ionization mass spectrometry (MALDI-MS)-based N-glycomics at 0, 1, and 3 days post-restimulation. A total of 27 distinct N-glycan structures were detected across all samples (**Fig. 5b and Supplementary Table 4**). Following folate deprivation, CD8^+^ T cells showed mild glycome remodeling on the day of restimulation, with dynamic remodeling of the N-glycome on days 1 and 3. We ranked the N-glycans based on their differential score (0 nM / 2,000 nM) on day 3 post-restimulation (**Fig. 5b**) and observed that enriched N-glycans in FA-deprived CD8^+^ T cells were predominantly composed of high GlcNAc (pyrimidine sugar) N-glycans, with several complex-type structures including Hex6dHex1HexNAc4 and Hex6HexNAc4 (**Supplementary Table 4**). Of particular interest, Hex6HexNAc5—a glycan recognized by the lectin Phaseolus vulgaris Leucoagglutanin (PHA-L)—was one of the observed glycans that exhibited a transient decrease at day 1 post-restimulation, followed by a marked increase by day 3. Given that PHA-L recognized glycans have previously been correlated with CD8^+^ T cell PD-1 and TIM-3 exhaustion marker expression in tumor-infiltrating lymphocytes (TILs)^77^, this temporal upregulation raises the possibility that exhaustion-associated glycan remodeling is initiated as early as day 3 post-restimulation in folate-deprived CD8^+^ T cells. Using flow cytometry, we validated and extended the timeline of our PHA-L N-glycan profiling, and confirmed the temporal pattern we observed in our glycomics data. We also noticed that PHA-L glycans remain high in FA-deprived CD8^+^ T cells at days 3, 5, and 7 post-restimulation, and correlated with high levels of the exhaustion markers, PD-1 and LAG-3 (**Extended Data Fig. 7a–c**).

**Fig. 5.**
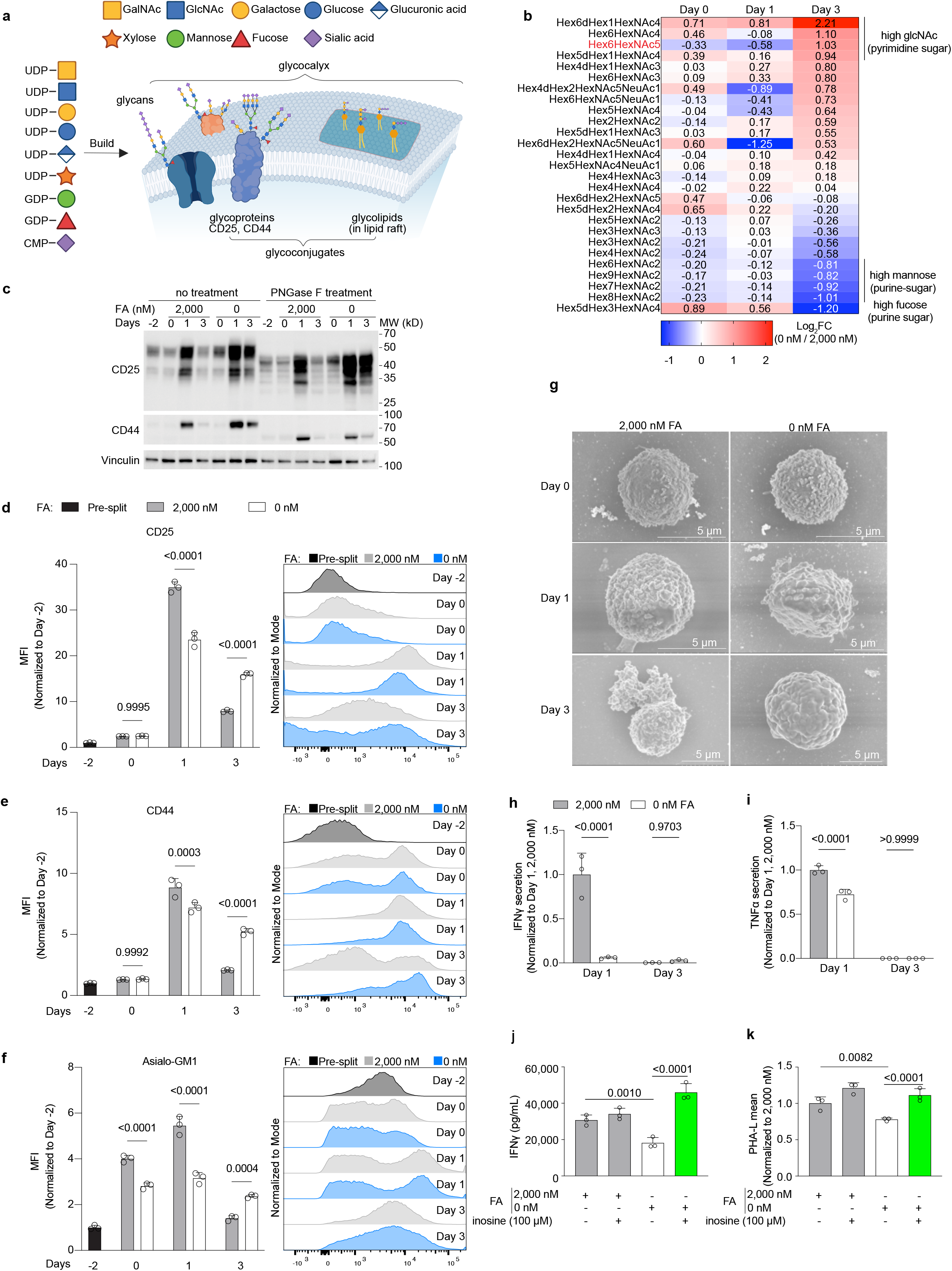
Folate deficiency induces global glycome remodeling, which can be rescued by inosine supplementation *ex vivo*. **a**, Schematic illustration of utilizing nucleotide sugars as building blocks for glycosylation in cells. Top, legend for sugars annotation. **b**, Heatmap of N-glycan glycomics of 0 nM FA / 2,000 nM FA across days 0, 1, and 3 post-restimulation characterized by MALDI-MS. N-glycans were ranked and ordered by their day 3 log_2_FC differential score. Red text indicates the N-glycan recognized by the lectin Phaseolus vulgaris Leucoagglutinin (PHA-L). **c**, Immunoblots for cell surface glycoproteins (CD25, CD44) in control (2,000 nM) and FA-deprived (0 nM) CD8^+^ T cells at various timepoints. Immunoblots for samples that were PNGase F treated (removes N-glycans on glycoproteins) are shown on the right. **d–f**, Flow cytometry cell surface quantification (left) and representative histograms (right) of CD25 (**d**), CD44 (**e**), and asialo-GM1 (**f**). **g**, Representative scanning electron microscopy images of CD8^+^ T cell glycocalyx morphology in 2,000 nM vs 0 nM FA at days 0, 1, and 3 post-restimulation. Scale bar, 5 µm. **h**, **i**, ELISA-based quantification of interferon gamma (IFNγ) (**h**) and tumor necrosis factor (TNFα) (**i**) secretion by control (2,000 nM) and FA-deprived (0 nM) CD8^+^ T cells at days 1 and 3 post-restimulation. **j**, **k**, Inosine supplementation (100 µM) in control and FA-deprived CD8^+^ T cells at day 1 post-restimulation. 2,000 nM FA conditions are colored grey, 0 nM FA is colored white, and 0 nM FA + inosine is colored green. **j**, ELISA-based quantification of IFNγ secretion. **k**, Flow cytometry of N-glycan cell surface abundance (PHA-L). Data are presented as mean and s.e.m. (*n* = 3 biological replicates for all panels). *P* values were determined by a repeated-measures two-way ANOVA. Statistical significance was evaluated either using a repeated-measures two-way ANOVA with Šídák’s multiple comparisons test to evaluate the treatment effect at each time point (**d–i**), or one-way ANOVA with Tukey’s post-hoc test for multiple comparisons (**j**, **k**).

T cells rely on several key glycoproteins for their activation and fitness^78^. Thus, we assessed the expression of CD25 (interleukin-2 receptor alpha chain [IL-2Rα]; essential for proliferation) and CD44 (essential for cell adhesion and migration) as two examples of key glycoproteins in CD8^+^ T cell function. We used Western blot for total cell protein level assessment (**Fig. 5c**) and flow cytometry for cell surface levels (**Fig. 5d,e**). Total CD25 and CD44 protein levels were elevated at day 1 post-restimulation in both FA-replete and FA-depleted conditions, but FA-deprived CD8^+^ T cells maintained high levels of total protein in both glycoproteins at day 3 (**Fig. 5c**). Using PNGase F enzymatic treatment, which removes N-glycans, we assessed the overall glycosylation profile of CD25 and CD44 (**Fig. 5c**), and found that CD44 is predominantly N-glycosylated, while CD25 is both N- and O-glycosylated (because it appeared as two bands at 50 kDa and 35 kDa, which represent intact CD25 (membrane-bound form) and cleaved CD25 (soluble form), with both hyper- and hypo-glycoforms remained).

At the cell-surface level, determined by FACS, we observed a pattern that is similar to PHA-L cell surface levels (**Fig. 5b and Extended Data Fig. 7a**): CD25 and CD44 cell surface levels were reduced at day 1 post-restimulation but were increased by day 3 (**Fig. 5d,e**). Additionally, cell surface levels of the glycolipid asialo-GM1 followed a similar trend (**Fig. 5f**). These data, combined with the whole-cell protein level assessment by Western blot, led us to propose that glycosylation processes in FA-deprived CD8^+^ T cells are faulty, leading to disrupted trafficking. Protein and lipid glycosylation begins at the endoplasmic reticulum (ER), and glycoconjugates are transported by coat protein complex II (COPII) vesicles to the Golgi apparatus for more terminal processing^79^. From there, these glycoconjugates can either be maintained intracellularly in various organelles, integrated into the plasma membrane and presented on the cell surface, or secreted extracellularly. However, if the glycosylation process is faulty (e.g., due to shortage of nucleotide sugars), it may cause a bottleneck in trafficking and accumulation of these proteins intracellularly at day 1 post-restimulation. This slowdown appears to be somewhat relieved by day 3 post-restimulation, allowing for glycoconjugates to finally reach the cell surface, as observed by flow analysis (**Fig. 5d–f**).

One noteworthy observation is that CD44 and CD25 glycoprotein cell surface levels were unaffected by low folate prior to restimulation (day 0) (**Fig. 5d,e**), while cell surface levels of the glycolipid asialo-GM1 were significantly reduced in low folate even before restimulation (**Fig. 5f**). This suggests that among the glycoconjugates we profiled, gangliosides are more sensitive to folate deprivation consequences than the glycoproteins CD25 and CD44, and this is in line with our CRISPR screen results (**Fig. 3a–c**).

In contrast to the pyrimidine glycans at the top of the glycomics data table (**Fig. 5b**), high mannose and high fucose glycans (purine sugars) were progressively downregulated at both days 1 and 3 post-restimulation (including Hex9HexNAc2, Hex7HexNAc2, Hex8HexNAc2, and Hex5dHex3HexNAc4), which is aligned with our findings that purine sugar biosynthesis is the main bottleneck in FA-deprived CD8^+^ T cells. These high mannose and high fucose glycans are important for the composition and construction of the glycocalyx. T cells extensively develop and elaborate their glycocalyx structure after activation, which is critical for their function^80,81^. Therefore, we investigated whether folate deprivation reduces glycocalyx complexity in restimulated CD8^+^ T cells. We performed scanning electron microscopy (SEM) to assess morphological changes (**Fig. 5g and Extended Data Fig. 7d,e**). Prior to restimulation on day 0, both FA-replete and FA-deprived activated CD8^+^ T cells had glycocalyx morphology as expected. On days 1 and 3 post-restimulation, FA-replete CD8^+^ T cells developed increasingly elaborate glycocalyx structures, while FA-deprived CD8^+^ T cells failed to develop an elaborate glycocalyx structure, in line with the glycomics data (lower part of the heatmap, **Fig. 5b**). Collectively, these data demonstrate that FA-deprived CD8^+^ T cells have impaired glycocalyx structure after restimulation.

Because activated CD8^+^ T cells secrete cytokines as part of their effector functions, we next investigated whether FA-deprived, restimulated CD8^+^ T cells can effectively secrete IFNγ and TNFα. Both human and mouse forms of IFNγ are glycosylated, and N-glycosylation of human IFNγ’s Asn-25 and Asn-97 sites are essential for its secretion^82,83^. Although human TNFα is not glycosylated and lacks predicted N-glycosylation sites, mouse TNFα is N-glycosylated^84^. We found that FA-deprived CD8^+^ T cells had significantly reduced IFNγ and TNFα secretion (**Fig. 5h,i**), which likely contributed to their impaired cytotoxic capacity (**Fig. 2a**).

Since we found that supplementation with the purine nucleoside, inosine, rescued nucleotide sugar levels in low folate (**Fig. 4g and Extended Data Fig. 6e–j**), we examined whether inosine supplementation could rescue glycoprotein IFNγ secretion in FA-deprived CD8^+^ T cells. Indeed, inosine supplementation alone was sufficient to rescue IFNγ secretion in FA-deprived CD8^+^ T cells (**Fig. 5j**). We also confirmed that inosine supplementation rescued PHA-L glycan cell surface expression, as a marker for a broader cell surface N-glycosylation state (**Fig. 5k**). Together, these data confirm the 1CM—purine nucleotide sugar metabolic axis and its crucial role in CD8^+^ T cell glycosylation and effector function.

### Folate deprivation triggers the pro-apoptotic unfolded protein response (UPR)

Cell surface glycoconjugates play essential roles in almost every aspect of T cell function, and glycosylation serves critical functions in the early secretory pathway. Most proteins are co-translationally translocated into the ER and simultaneously N-glycosylated, promoting proper protein folding^85^. Impaired N-glycosylation of nascent proteins leads to protein misfolding^86^, resulting in misfolded protein accumulation in the ER and triggering the unfolded protein response (UPR). The ER chaperone BiP (binding immunoglobulin protein, aka heat shock protein family A member 5 [HSPA5]) binds to misfolded proteins, which releases the ER stress sensors IRE1α and PERK to undergo homodimer activation (**Fig. 6a**). Transient or mild stress stimulates the adaptive UPR, which temporarily halts protein translation and mRNA splicing of the transcription factor, X-box binding protein 1 (XBP1). Spliced XBP1 (sXBP1) controls target gene expression that modulates protein folding, secretion, and endoplasmic reticulum-associated protein degradation (ERAD). While glycosylation defects can activate the adaptive UPR, activation and resolution of the adaptive UPR is also inherent to T cell development, activation, and expansion due to increased synthesis demands during these stages^87,88,89^. If protein-folding homeostasis fails to be restored, prolonged UPR stimulation shifts the cell towards a pro-apoptotic program. Key hallmarks of the pro-apoptotic UPR are chronic translational inhibition, PERK hyperactivation, downstream upregulation of the transcription factor CHOP, downregulation of sXBP1 levels, and activation of the intrinsic apoptotic pathway.

**Fig. 6.**
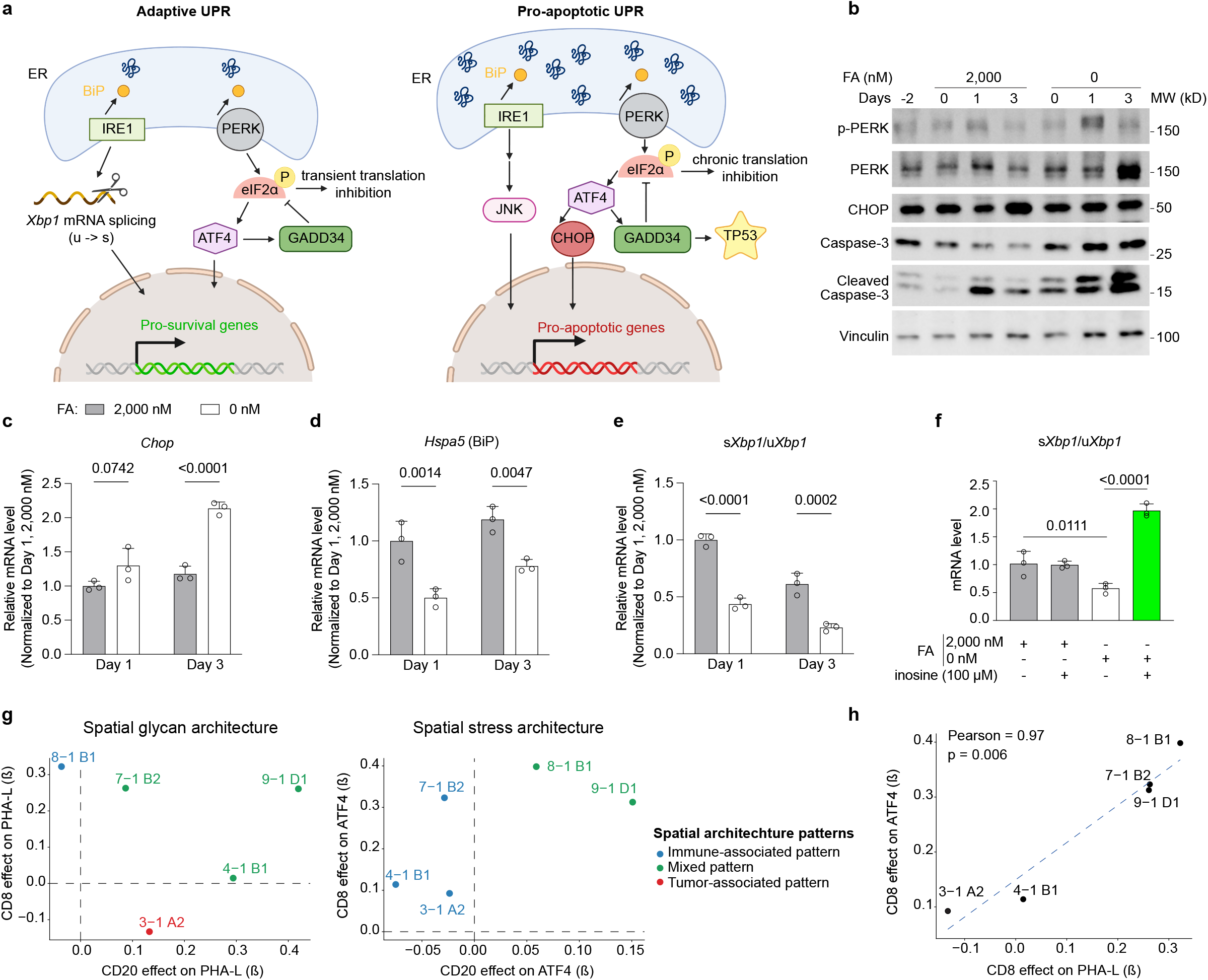
Folate deprivation triggers the pro-apoptotic unfolded protein response (UPR) in CD8^+^ T cells. **a**, Schematic of the adaptive and pro-apoptotic unfolded protein responses (UPR) triggered by glycosylation defects and subsequent protein unfolding in the endoplasmic reticulum (ER). **b**, Immunoblots for markers of adaptive and pro-apoptotic UPR in control (2,000 nM) and FA-deprived (0 nM) CD8^+^ T cells at days 0, 1 and 3 post-restimulation: p-PERK, phosphorylated (T982) PERK; p-IRE1, phosphorylated (S724) IRE1. **c–e**, qPCR quantification of *Chop* (pro-apoptotic UPR marker) (**c**), and adaptive UPR markers Heat shock protein 5 (*Hspa5*), which encodes for binding immunoglobulin protein (BiP) (**d**), and the ratio of spliced *Xbp1* (s*Xbp1*) over unspliced *Xbp1* (u*Xbp1*) (**e**) XBP1: X-box binding protein 1. All plotted transcript levels are in control vs. FA-deprived CD8^+^ T cells at days 1 and 3 post-restimulation. **f**, qPCR quantification of s*Xbp1/*u*Xbp1* transcript levels (adaptive UPR marker) in inosine-supplemented (100 µM) control and FA-deprived CD8^+^ T cells on day 1 post-restimulation. **g, h,** Multivariable spatial global regression modeling of immunofluorescence (IF) stained human PCNSL tissue sections (*n* = 5). Spatial IF phenotype maps of tumor cells (CD20), CD8 T cells (CD8), branched N-glycans (PHA-L), and the stress responsive transcription factor, ATF4 were constructed and used for regression coefficient (β) calculations. **g**, Spatial glycan (top, PHA-L) and stress (bottom, ATF4) architectures of PCNSL tumors. The relationship strength (β coefficient) of CD8 (β1, y-axis) and CD20 (β2, x-axis) signal intensities with PHA-L and ATF4 are shown. Each data point represents one tumor and is colored by spatial architecture category, which were classified by β coefficient (blue = immune-associated pattern [CD8 dominant, β1 > β2], red = tumor-associated pattern [CD20 dominant, β2 > β1], and green = mixed pattern [both coefficients contribute substantially to module signal]). **h**, Correlation between CD8-associated effects on lectin and ATF4 signals across PCNSL tumors. Data in **c–f** represent 3 biological replicates and are presented as mean and s.e.m. β coefficients; data in **g, h** were calculated from linear regression models across spatial bins (400 µm resolution) within each tumor. Statistical significance was determined by a repeated-measures two-way ANOVA with Šídák’s multiple comparisons test to evaluate the treatment effect at each time point used (**c–e**), one-way ANOVA with Tukey’s post-hoc test for multiple comparisons (**f**), or *t*-test on Pearson’s correlation coefficient (**h**).

Since we posited that FA-deprived CD8^+^ T cells suffer from aberrant protein trafficking due to perturbed glycosylation (**Fig. 5c–f**), we reasoned that FA-deprived CD8^+^ T cells may have elevated UPR activation. To investigate this, we characterized the activation status of the ER stress sensor PERK and its downstream effectors. Compared to FA-replete CD8^+^ T cells, FA-deprived CD8^+^ T cells had higher CHOP on day 1 and higher p-PERK and cleaved caspase-3 on days 1 and 3 post-restimulation (**Fig. 6b**). We confirmed the activation of an apoptotic program by flow cytometry for the apoptotic marker Annexin V on FA-deprived CD8^+^ T cells (200, 100, 50, 20, and 0 nM folate) and observed dose-dependent elevation in Annexin V^+^ populations across all FA-deprived groups by day 3 (**Extended Data Fig. 8a**). We also assessed changes in the adaptive and pro-apoptotic UPR transcriptional programs in FA-replete and FA-deprived CD8^+^ T cells, and observed increased pro-apoptotic CHOP transcript levels, as well as reduced adaptive *BiP* and spliced to unspliced ratio of XBP1 (s*Xbp1*/u*Xbp1*) transcripts (**Fig. 6c–e**). Downstream effectors of the adaptive UPR, including ER degradation-enhancing α-mannosidase-like protein 1 (EDEM1; an ERAD protein) and the ER chaperone protein, glucose-regulated protein 94 (aka grp94/gp96, the ER resident form of heat shock protein 90 kda beta member 1 [HSP90B1]; can bind to unfolded proteins and alleviate UPR stress), were also downregulated transcriptionally, but only on day 1 (**Extended Data Fig. 8b,c**). Inosine supplementation was sufficient to rescue the s*Xbp1*/u*Xbp1* ratio in FA-deprived CD8^+^ T cells on day 1 post-restimulation (**Fig. 6f**), restoring the adaptive UPR and highlighting 1CM’s critical role in the observed ER stress.

We next asked if the ER stress we observed in CD8^+^ T cells *ex vivo* might also be happening in human PCNSL tumors, and whether this is associated with cell surface glycosylation status. To address this, we performed immunofluorescence staining of tumor samples from five PCNSL patients (**Extended Data Fig. 9a**). The analysis was done on verified tumor regions (**Extended Data Fig. 9b**) and included markers for malignant B cells (CD20), CD8^+^ T cells (CD8), glycosylation (PHA-L), and ER stress (ATF4) (**Extended Data Fig. 9c**). Using the signal intensities from CD8/ATF4/PHA-L and CD20/ATF4/PHA-L co-staining (**Extended Data Fig. 9d**) from serial sections, we performed multivariable spatial regression modeling to investigate the independent contributions of CD8 vs. CD20 (ß coefficients) on spatial glycan (PHA-L) and ER stress (ATF4) architectures (**Fig. 6g and Extended Data Fig. 9e**). We observed some overlap between the glycan and stress modules, but the coupling patterns (i.e., immune-associated, tumor-associated, and mixed) across tumors varied. Next, we focused on the tumor-infiltrating CD8^+^ T cells by assessing the glycan and stress module strength in relation to the CD8^+^ Tumor Infiltration Index (TII; readout for the ratio of positive CD8^+^ signal in CD20 high regions) (**Extended Data Fig. 9f**). We observed that PCNSL samples with a higher CD8^+^ TII score tended to also have higher glycan and stress module signal intensity, suggesting that intratumoral CD8^+^ T cells have higher PHA-L recognized glycan(s) and ATF4 expression. Additionally, we found a positive correlation between the CD8^+^ effect on PHA-L signal and the CD8^+^ effect on the ATF4 stress marker signal (**Fig. 6h**). Altogether, these analyses reveal heterogenous tumor- and immune-associated spatial architectures across individual PCNSL patients, but a consistent correlation between the immune-associated pattern of both glycan and stress markers.

### Folate transporter overexpression enhances CD8^+^ T cell fitness *in vivo*

While inosine supplementation was sufficient to rescue glycosylation defects, cytokine secretion, and restore the adaptive UPR in FA-deprived CD8^+^ T cells *ex vivo,* we observed significantly elevated inosine levels in both B16-OVA and MC38-OVA TIF (**Extended Data Fig. 3d,h,l**). Thus, we rationalized that inosine supplementation may not be beneficial to CD8^+^ T cells that are already in an inosine-high environment *in vivo* and sought to enhance CD8^+^ T cells in a cell-specific manner utilizing a genetic approach.

To assess what 1CM and nucleotide sugar metabolism genes are lowly expressed in tumor-infiltrating CD8^+^ T cells, we analyzed scRNA-seq data from the publicly available GSE116390 dataset^90^ of tumor-infiltrating CD8^+^ T cells from B16 melanoma mouse tumors (**Fig. 7a**). We found that genes encoding proteins responsible for folate uptake (i.e., *Folr1, Folr2, Slc19a1,* and *Slc46a1*) are very lowly expressed in most CD8^+^ tumor infiltrating cell types in B16 tumors, including effector T cells (**Fig. 7b**). Although some of these proteins are not expected to be expressed on T cells, we were surprised by low expression levels of *Slc19a1* (solute carrier family 19 member 1; reduced folate carrier), a ubiquitously expressed folate transporter. We therefore set to overexpress folate transporters on CD8^+^ T cells with the goal of enhancing folate uptake. This approach is also aligned with the need to improve folate uptake in cases of tumor-induced folate depletion^26,27,28,29,30,31^ and in the folate-depleted TME (**Fig. 2e,f and Extended Data Fig. 3m**).

**Fig. 7.**
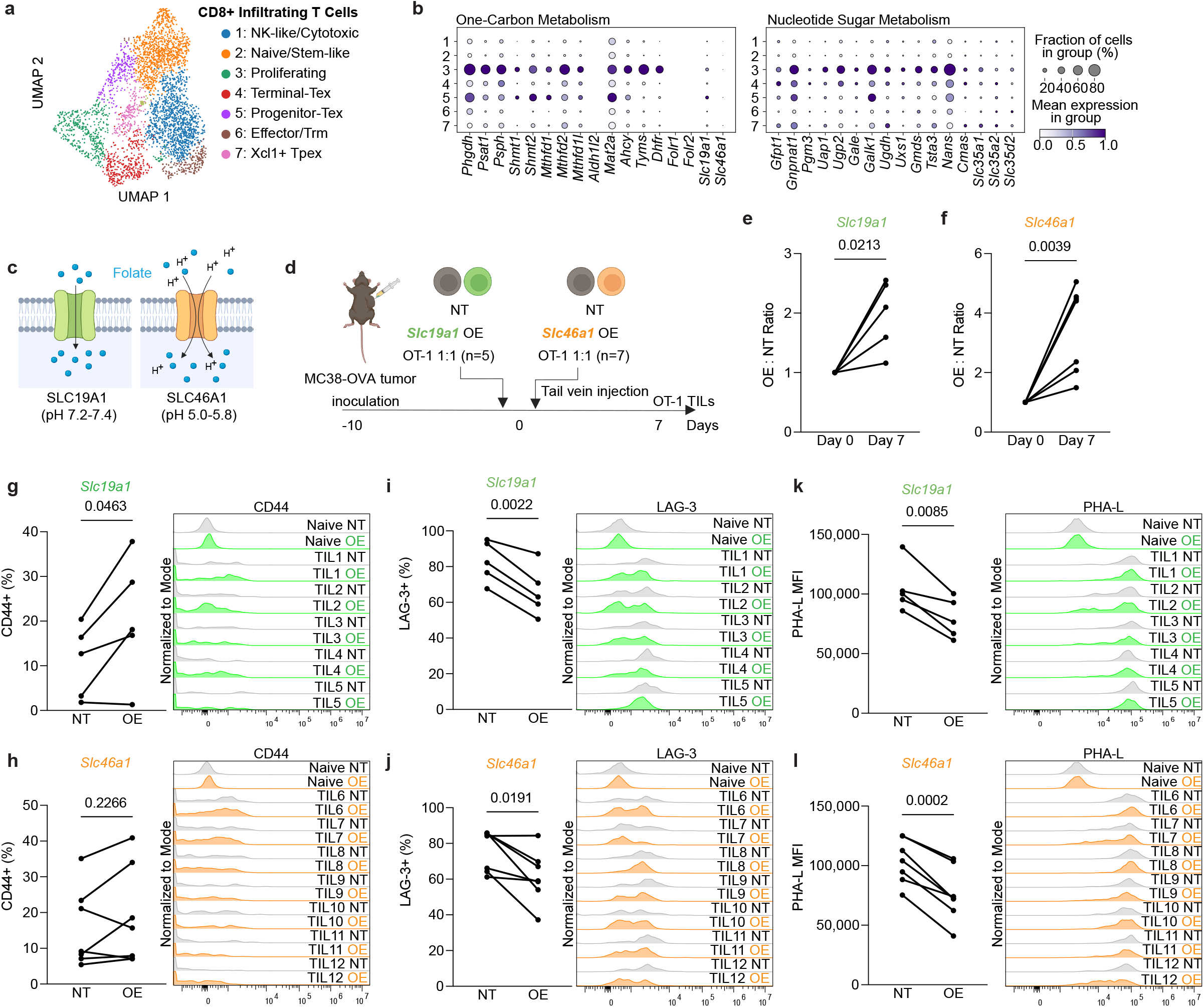
Overexpression of the folate transporters SLC19A1 and SLC46A1 enhances intratumoral CD8^+^ T cells *in vivo*. **a,** UMAP representation of integrated scRNA-seq of tumor infiltrating CD8^+^ T cells from B16 melanoma mouse tumors (GSE116390). **b,** Percentage of positive cells and average expression for one-carbon metabolism (left) and nucleotide sugar metabolism (right) genes. **c**, Schematic of the folate transporters encoded by *Slc19a1* (reduced folate carrier 1, RFC1) and *Slc46a1* (proton-coupled folate transporter, PFCT). **d**, Experimental schematic of non-targeting control (NT) vs. *Slc19a1* or *Slc46a1* overexpressing (OE) CD8^+^ T cells competitive adoptive transfer into MC38-OVA tumor-bearing mice (*Slc19a1 OE, n* = 5; *Slc46a1 OE, n =* 7). *Slc19a1* OE is represented in green text, while *Slc46a1* OE is represented by dark orange text. **e**, **f,** Flow cytometry of the ratio of *Slc19a1* OE (**e**) or *Slc46a1* OE (**f**) vs. NT control tumor infiltrating CD8^+^ T cells. **g**, **h**, Levels of the activation marker, CD44, on *Slc19a1* OE (**g**) or *Slc46a1* OE (**h**) vs. NT control tumor infiltrating CD8^+^ T cells. Quantification of the percentage of CD44^+^ cells (left) and representative histograms (right) are shown. **i**, **j**, Levels of the exhaustion marker, LAG-3, on *Slc19a1* OE (**i**) or *Slc46a1* OE (**j**) vs. NT control tumor infiltrating CD8^+^ T cells. Quantification of the percentage of LAG-3^+^ cells (left) and representative histograms (right) are shown. **k**, **l**, Levels of the N-glycan recognized by the lectin Phaseolus vulgaris Leucoagglutinin (PHA-L) on *Slc19a1* OE (**k**) or *Slc46a1* OE (**l**) vs. NT control tumor infiltrating CD8^+^ T cells. Quantification of the median fluorescent intensity of PHA-L^+^ cells (left) and representative histograms (right) are shown. Statistical significance (**e–l**) was calculated by a two-tailed paired Student’s *t*-test.

We set to overexpress the folate transporters SLC19A1 and SLC46A1 in adoptively transferred CD8^+^ T cells to test whether this would improve their fitness. *Slc19a1* encodes the reduced folate carrier (RFC), which is a ubiquitously expressed plasma membrane transporter that mediates the bidirectional transport of reduced folates at a neutral pH. *Slc46a1* (solute carrier family 46 member 1) encodes the proton-coupled folate transporter (PFCT), which is highly expressed on the apical membrane of the small intestine and functions optimally at an acidic pH (**Fig. 7c**). Although SLC46A1 is not as highly expressed in CD8^+^ T cells compared to SLC19A1, we decided to test this transporter because the TME is acidic.

We performed a competitive adoptive transfer experiment in tumor-bearing mice to evaluate whether folate transporter overexpression (OE) would improve adoptive CD8^+^ T cell fitness in tumors (**Fig. 7d**). CD45.1^+^/2^+^ heterozygous mice were inoculated with MC38-OVA tumors which grew for ten days to allow for folate deprivation in the TME. An equal ratio of non-targeting control (NT) CD45.2^+^ OT-I cells and folate transporter overexpressing (*Slc19a1 OE* or *Slc46a1 OE*) CD45.1^+^ OT-I cells were mixed and adoptively transferred into MC38-OVA tumor-bearing mice, and OT-I TILs were isolated for flow analysis seven days later (**Extended Data Fig. 10a**). *Slc19a1 OE* and *Slc46a1 OE* TILs were both significantly enriched intratumorally compared to NT controls (**Fig. 7e,f**). *Slc19a1 OE* TILs had significantly increased expression of the activation marker CD44 (**Fig. 7g**), while *Slc46a1 OE* TILs trended towards increased CD44 levels (**Fig. 7h**). Furthermore, both *Slc19a1 OE* and *Slc46a1 OE* TILs had reduced expression of the exhaustion marker LAG-3 (**Fig. 7i,j**). Notably, cell-surface complex N-glycan levels were reduced on both *Slc19a1 OE* and *Slc46a1 OE* TILs (**Fig. 7k,l**), demonstrating that increased folate availability modifies TIL surface glycosylation profiles. This observation is consistent with Kearney and Vervoort et al.’s observation that less exhausted B16-OVA and MC38-OVA TILs have lower PHA-L levels^77^, as well as our *ex vivo* data demonstrating that FA-replete, restimulated CD8^+^ T cells downregulate PHA-L levels from day 3 onwards (**Extended Data Fig. 7a**). Together, these data suggest that folate transporter overexpression increases TIL numbers and sustains their function and fitness in tumors *in vivo*.

## DISCUSSION

Significant advances in immunotherapy in the last couple of decades has revolutionized cancer treatment. However, while the estimated percentage of patients eligible for immune checkpoint therapies has increased, the overall response rate for most cancer subtypes only reaches 20%^91,92^. Specifically, CAR-T therapies face significant hurdles in solid tumors, with an overall 9% pooled response rate^93^. Thus, understanding the factors governing immunotherapy response and discovering combination therapies to bolster existing immunotherapy regimens are a few of the urgent questions currently driving oncology research. Enhancing 1CM in CD8^+^ T cells by 1C unit supplementation can improve αPD-1/αPD-L1 treatment in mouse tumor models^23,24,25^. However, the role of the required 1C carrier, folate, in CD8^+^ T cell function is underexplored. Literature reporting quantitative levels of 5-methyl THF in biological fluids is sparse. Among the published work, Schmitz and colleagues used a reduced folate ternary complex assay to estimate mouse plasma 5-methyl THF levels in C3H/HeJ (CH3) mice, which was approximately 76 nM^66^. Sullivan and colleagues applied LC-MS for folate detection in NOD.Cg-*Prkdcscid Il2rgtm1Wjl*/SzJ (NSG) mice and estimated 5-methyl THF levels to be slightly higher, at approximately 100 nM^67^. However, 5-methyl THF levels in mouse TIF is reported here for the first time.

Our focus on PCNSL as our clinical model was motivated by the fact that this tumor is responsive to the antifolate drug, methotrexate, suggesting reliance on 1CM^94^. PCNSL often exhibits an immune-poor microenvironment characterized by low T cell infiltration^38^, HLA I/II loss^95,96^, and lack of T cell activation with high exhaustion marker expression^39^, consistent with a functionally T cell excluded phenotype. Although there is extensive literature investigating the connection between a metabolically hostile TME and impaired CD8^+^ T cell function^97,98^, there are a lack of studies investigating PCNSL metabolic vulnerabilities.

While both proliferating cancer and immune cell populations in PCNSL patients’ samples highly expressed folate metabolism genes, CD8^+^ T cells were exhausted while the cancer cells were thriving. This led us to ask whether folate is a limiting nutrient in the TME and if folate is utilized differently in cancer and T cells. First, we indeed found that 5-methyl THF (circulating folate form) is one of the few depleted metabolites in MC38-OVA and B16-OVA TIF. We then sought to deepen our understanding of T cell folate utilization and need, and how this may differ from cancer cell folate utilization. Next, we found that folate-deprived activated CD8^+^ T cells have dysregulated nucleotide sugar metabolism. We suggest that this explains the differential essentiality of genes involved in ganglioside biosynthesis, which we observed in a genome-wide, loss of function CRISPR screen of folate replete and folate deplete conditions in primary mouse CD8^+^ T cells. We used glycomics to verify that folate deprivation induced globally aberrant glycosylation in activated CD8^+^ T cells. We also found that perturbed cell surface glycan levels informed the dynamics of cell surface glycoconjugates’ levels (e.g., CD25, CD44, asialo-GM1) in folate-deprived CD8^+^ T cells following restimulation. Additionally, glycomics revealed purine sugar- and pyrimidine sugar-distinct fates, which led us to study the differential contribution of purine sugar and pyrimidine sugar biosynthesis pathways in folate-deprived CD8^+^ T cells. Through this investigation, we discovered a regulatory crosstalk between purine and pyrimidine metabolism. Specifically, we found that transcriptional regulation of the pyrimidine synthesis enzyme, CMPK2, is inosine-dependent.

These findings led us to focus on inosine as a key metabolite at the junction of T cell decision making and fitness. Not only do inosine levels dictate purine and purine-sugar availability, but it also allows for enhanced pyrimidine and pyrimidine-sugar synthesis. Indeed, inosine supplementation was sufficient to rescue several key features in the pathology induced by folate deprivation, including cytokine secretion (**Fig. 5j**) and cell surface N-glycan levels (**Fig. 5k**). Additionally, inosine supplementation shifted folate-deprived CD8^+^ T cells towards the adaptive UPR (**Fig. 6f**), and thus constructive ER stress management. Interestingly, inosine is not depleted in the TME (**Extended Data Fig. 3d,h,l**), implying that locally high levels of inosine *in vivo* is not sufficient for the metabolic rescue of T cells. This can potentially be explained by the double-edged nature of the inosine transporter, equilibrative nucleoside transporter 1 (ENT1): while T cells uptake inosine with ENT1, its expression and activity also inhibits T cell function and tumor cell killing^99^.

At the cell biology level, we revealed that low folate-induced aberrant glycosylation triggered the pro-apoptotic UPR, which is likely part of the underlying cause of reduced fitness and functionality of folate-deprived T cells. Previous studies have demonstrated that the UPR, an ER stress pathway, can both promote and hinder T cell anti-tumor responses^100,101,102,103,104^. We found that low folate-induced glycosylation defects can push activated CD8^+^ T cells towards a pro-apoptotic UPR, which can be rescued and shifted towards an adaptive UPR through inosine supplementation. In PCNSL patient samples, we observed a correlation between N-glycan levels on CD8^+^ T cells (which may reflect their need for proper nucleotide-sugar synthesis) and UPR stress, which is induced when glycosylation is perturbed. This implies that high glycosylation flux induces the UPR in humans as well, although our patient sample data did not distinguish between the adaptive and apoptotic UPR. Together, our findings link disturbed 1CM to the pro-apoptotic UPR, highlighting the role of 1CM in supporting basic cell functions (i.e., translation and trafficking) and further establishing the UPR’s role as a critical rheostat for T cell fate.

As a forward-facing approach, we aimed to capitalize on our findings for strategic planning of metabolic enhancement of T cells. We observed that, of the 1CM enzymes, the proteins responsible for folate uptake are the most lowly expressed on CD8^+^ TILs. This is a possible confounder for T cells in the competitive, folate-depleted TME. We therefore tested the impact of folate transporter overexpression on adoptively transferred CD8^+^ T cells, and found that this approach can improve TIL numbers, activation, and reduce exhaustion markers—including the glycosylation exhaustion marker PHA-L.

Our study uncovered a previously unappreciated connection between 1CM and glycosylation in activated CD8^+^ T cells. While glycosylation has been widely studied for its essential role in CD8^+^ T cell anti-tumor functions^73,105,106,107^, the role of 1CM in sustaining this important cell biology process has not been addressed in T cells. Further, and especially given the limiting availability of folate in the TME, it is important to understand the metabolic and biochemical consequences of folate deprivation in activated CD8^+^ T cells because of its immediate implications on T cell function. For example, our results demonstrate that folate-deprived CD8^+^ T cell glycosylation is globally impaired at multiple levels (e.g., glycan, glycoconjugate, and glycocalyx), and this included key functional cell-surface expressed and secreted proteins. As nearly all T cell surface receptors (e.g., T cell receptor (TCR), CD44, IL-2R, PD-1), lipid rafts, and secreted factors are glycosylated, understanding the glycosylation changes that occur in low folate may elucidate future therapeutic avenues to enhance CD8^+^ T cell activation and functionality in folate-deprived tumors.

While this study focused on the functional outcome of folate deprivation in CD8^+^ T cells, low folate levels in the TME likely impact all cancer cells and tumor-associated cells (e.g., stromal cells like cancer-associated fibroblasts (CAFs) and other immune cells like dendritic cells, natural killer cells (NKs), and macrophages). Altered glycosylation is known to be a hallmark of cancer^108,109^, and tumor stromal cells may have increased UPR signaling to secrete pro-tumor factors, aiding tumorigenesis^110^. Also, a recent study showed that tumor-infiltrating NK cells have disrupted glycocalyx morphology, facilitating cancer cell immune evasion^111^. Moreover, the UPR has been shown to be essential for dendritic cell homeostasis^112^ and B cell differentiation^113^. Interestingly, tissue-resident FOLR2^+^ macrophages are characterized by their high overexpression of folate receptor β (FRβ, encoded by *FOLR2*), suggesting that FOLR2^+^ macrophages may be more heavily impacted by low folate levels in the TME. FOLR2^+^ macrophages have been found to be enriched in human breast cancer and lung adenocarcinoma tumors^114,115^; however, whether they promote or inhibit tumor progression is currently unclear^116^. Future work investigating the impact of tumor-derived folate deprivation on other cells in the TME will be critical to uncovering cell-type specific and cell-cell interaction contributions to the immunosuppressive TME.

We found that folate transporter overexpression confers a competitive advantage to adoptively transferred CD8^+^ T cells. Folate-enhanced CD8^+^ T cells were increased intratumorally—whether this is due to enhanced migration, infiltration, or retention is unknown and should be investigated in future work. Also, folate transporter overexpressing CD8^+^ TILs had higher activation and reduced exhaustion markers, suggesting that improving folate availability is a viable method to sustain effector function and persistence, which are crucial goals of immunotherapy treatment. Since folate is systemically deprived in various human cancers, these patients may best benefit from adoptive cell therapies that overexpress folate transporters. Importantly, this approach can also be incorporated into systems of *in vivo* CAR-T induction, making this an important consideration for next-generation immunotherapy. Future studies evaluating this approach in human CAR-T cells will be important in assessing its potential therapeutic benefits for both conventional and *in vivo* CAR-T therapies.

### Limitations of the study

The presented research examines perturbed 1CM-mediated effector T cell impairments and T cell genetic enhancement through folate transporter overexpression. Further work is required to understand the impact of nutritional folate status, dietary intake, and folate levels in different organs. This may help stratify which patients and cancer types are at higher risk of folate deprivation induced-T cell defects, and subsequently best benefit from this therapeutic approach. There were a limited number of PCNSL patients (*n* = 5) profiled for spatial phenotyping of glycan and stress architectures. Further studies with a larger number of patients, and possibly additional cancer types, will be required to understand the applicability of the glycosylation-UPR pathway to human patients.

A limitation of our CRISPR screen is that although sgRNA abundance can reveal proliferation and survival differences, it does not allow us to survey whether the surviving CD8^+^ T cells have modified effector functionality; it is possible that surviving *Uxs1, B4galnt1,* and *B3galt4* KO CD8^+^ T cells with potential glycosylation and ganglioside defects may have diminished effector capacity.

Additional studies are warranted to better understand where glycoproteins may accumulate intracellularly along the secretory pathway and the molecular mechanisms underlying this potential traffic jam.

## RESOURCE AVAILABILITY

### Lead Contact

Additional information and requests for raw data or reagents should be directed to, and will be made available by, the corresponding author, Dr. Naama Kanarek

### Materials availability

All unique, stable reagents generated in this study will be made available by the lead contact with a completed material transfer agreement. Plasmids generated in this study will be deposited to Addgene.

### Data and code availability

Data are available in the main text, figures, supplemental files, and/or indicated data repositories. RNA-seq data were accessed from the European Genome-Phenome Archive (EGAD; accession number EGAC50000000250) and NCBI’s Gene Expression Omnibus (GEO; accession number GSE116390). Metabolomics data will be available upon deposition to Metabolomics Workbench. RNA-seq, glycomics, and metabolomics analysis details are provided in the STAR Methods section.

## ACKNOWLEDGEMENTS

We thank members of Roberto Chiarle’s lab for scientific discussions contributing to this manuscript. P.W. is supported by a Croucher Foundation Fellowship (FP01030497). A.Y. is supported by a National Science Foundation Graduate Fellowship (NSF GRF; DGE-2146755) and the Landry Cancer Biology Honorary Fellowship. A.S.E. is supported by the Swiss National Science Foundation (SNSF) under the SPARK funding scheme (CRSK-3_229139). L.H. is supported by the Deutscher Akademischer Austauschdienst (DAAD: German Academic Exchange Service) under the PROMOS scholarship. R.C.S. is supported by the National Institute of Aging (NIA; R01AG066653, R01AG078702), National Cancer Institute (NCI; R01CA266004, R01CA288696), and National Institute of Neurological Disorders and Stroke (NINDS; RM1NS133593). J.T. is supported by a Geneva Translation Oncology Award (GTO), which is part of the Fondation privée des HUG (Private Foundation of the Geneva University Hospitals). M.W. is supported by the Max Cloëtta Foundation and a GTO award. N.H. is supported by a Mark Foundation Endeavor Award and the Adelson Medical Research Foundation. N.K. is supported by the Pew Charitable Trusts Biomedical Science Innovation Fund (Biomed Scholars-2022-A-20060), The Smith Family Awards Program for Excellence in Biomedical Research, NCI R01CA279550-01, The Binational Science Foundation (2021127), and the Boston Children’s Hospital Research Executive Council Pilot Award. Graphical abstract and experimental schematic images were created with BioRender.com.

## AUTHOR CONTRIBUTIONS

N.K. supervised the study and acquired funding. P.W., A.Y., and N.K. designed and conceptualized the study. P.W. cloned the constructs, designed and performed experiments, and analyzed data. A.Y. designed and performed animal experiments, bench experiments, and analyzed data. M.M., K.H., M.M., and L.H. performed animal experiments and collected data. S.M., R.C.L., M.G., M.Z., J.R.B., E.O., A.Y.L.W., B.P., and M.W. performed and/or assisted with experiments and collected data. A.S.E., P.W., and A.G. performed biostatistical analysis. R.C.S., J.T., M.W., M.S., N.H., and N.K. advised on the experimental design. A.Y., P.W., and N.K. visualized and wrote the manuscript. All authors commented on and revised the manuscript.

## SUPPLEMENTARY INFORMATION: TABLES

### Supplementary Data Tables

| Table | Title | Description | Sheet |
| --- | --- | --- | --- |
| Supplementary Table 1 | gRNA used for single gene KO screen validation | Related to Extended Data Fig. 4a–c | S1_gRNA_List |
| Supplementary Table 2 | Differentially expressed metabolites between 0 nM and 2,000 nM folic acid | Related to Fig. 3b | S2_T cell_Metabolomics_Volcano |
| Supplementary Table 3 | Primers used for quantitative PCR (qPCR) analysis | Related to Fig. 4i–o, 6c–f and Extended Data Fig. 6m,n and 8b,c | S3_qPCR_Primers |
| Supplementary Table 4 | N-glycan reference library used for glycomic analysis | Related to Fig. 5b | S4_NGlycan_Reference |

## KEY RESOURCES TABLE

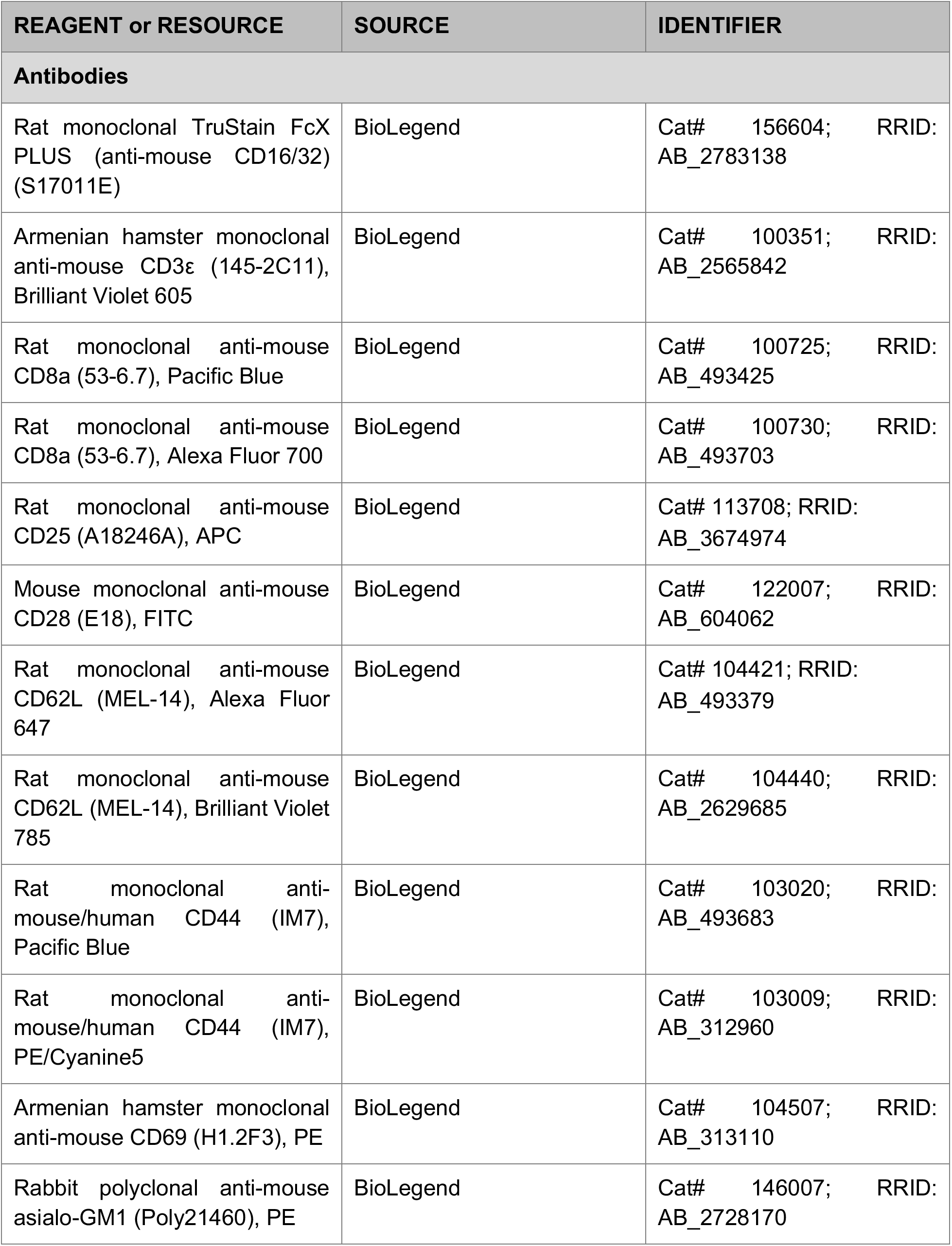
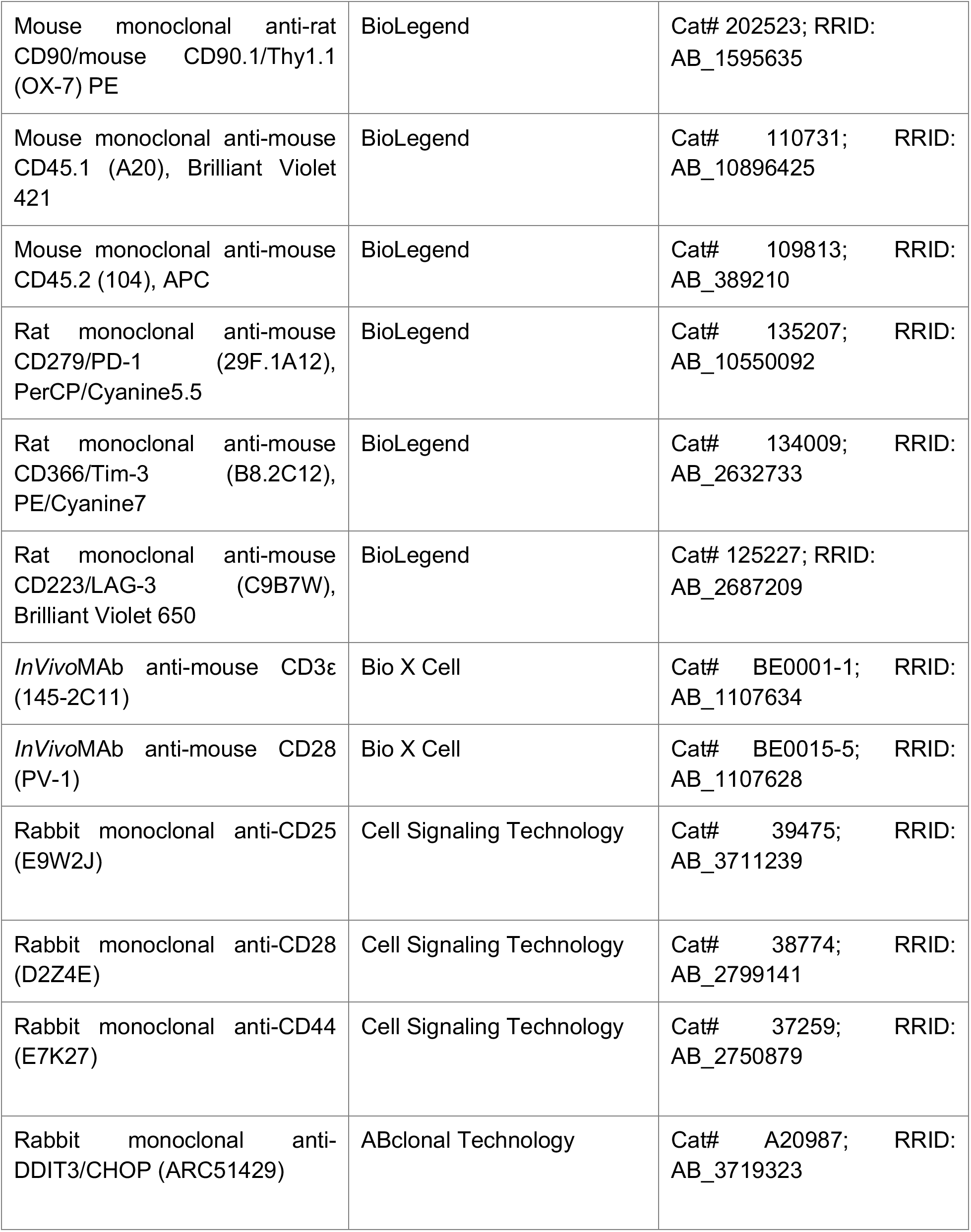
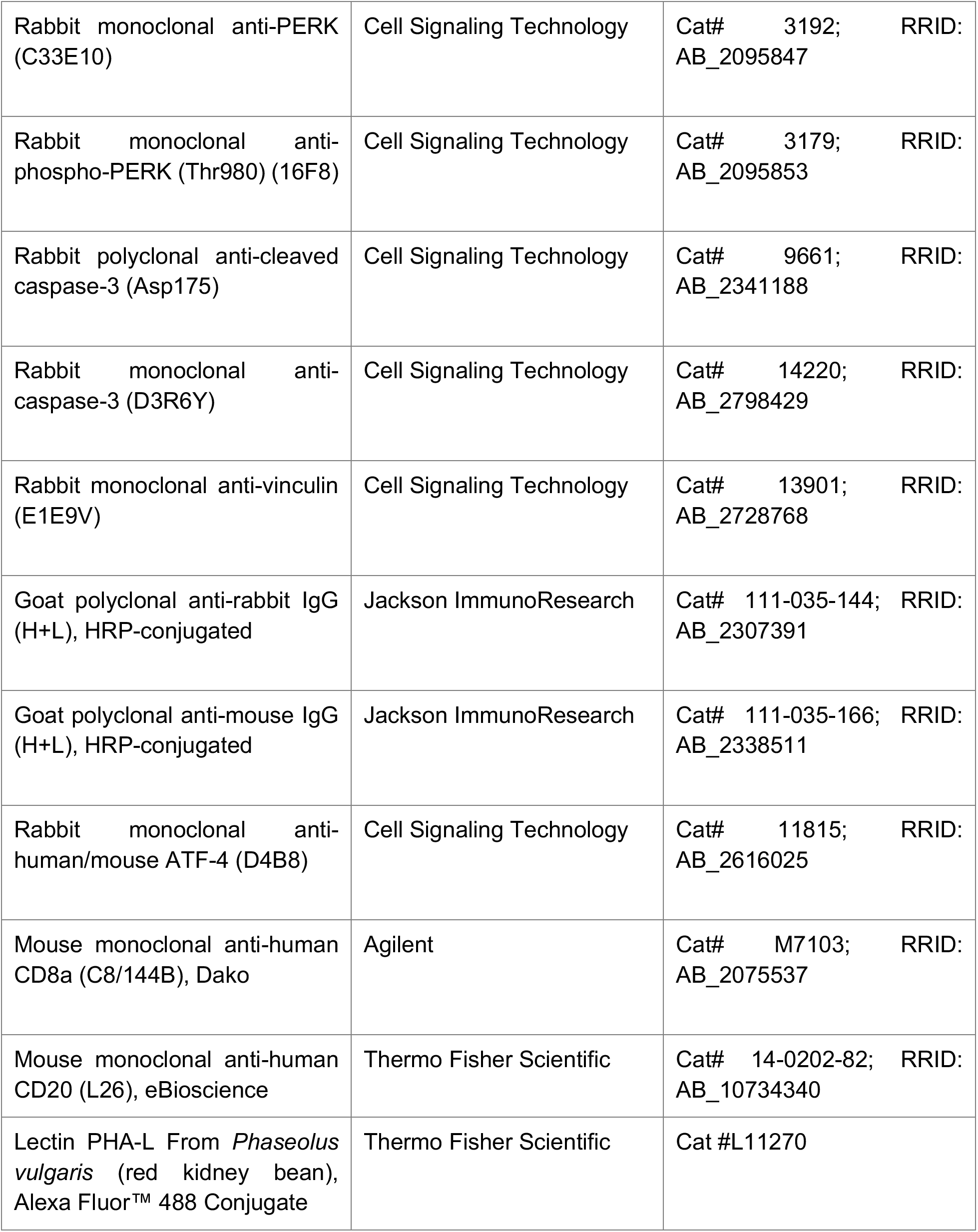
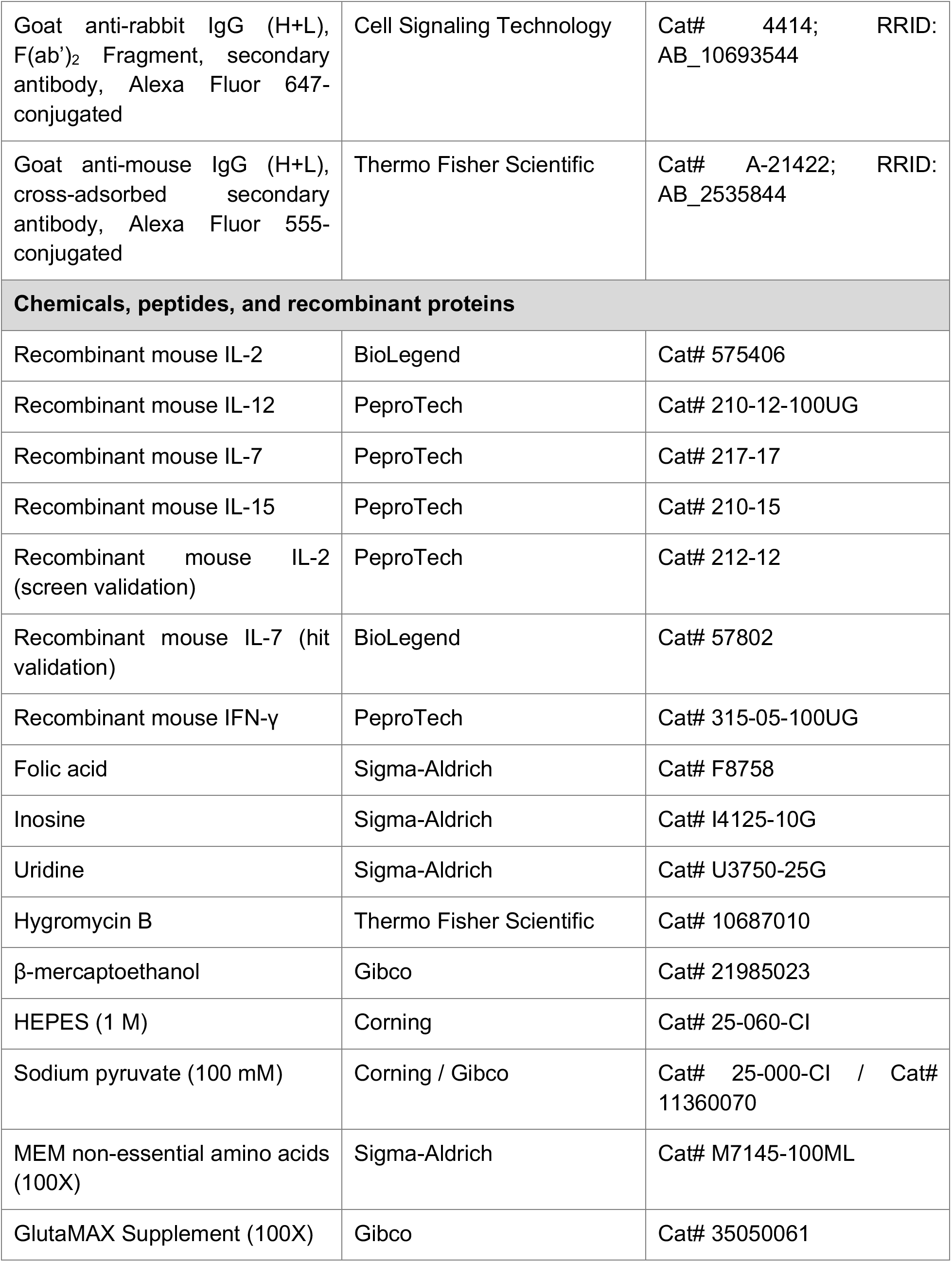
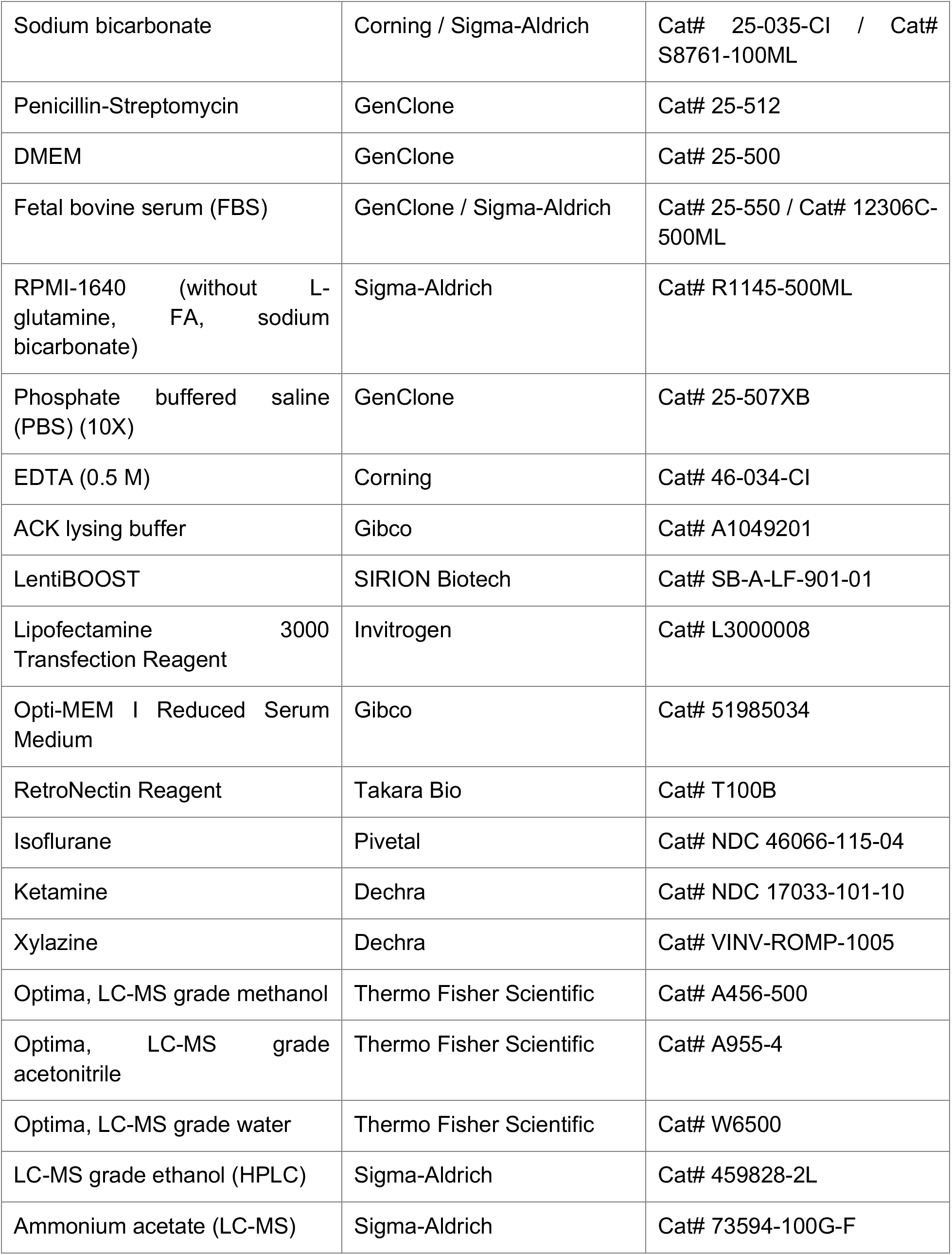
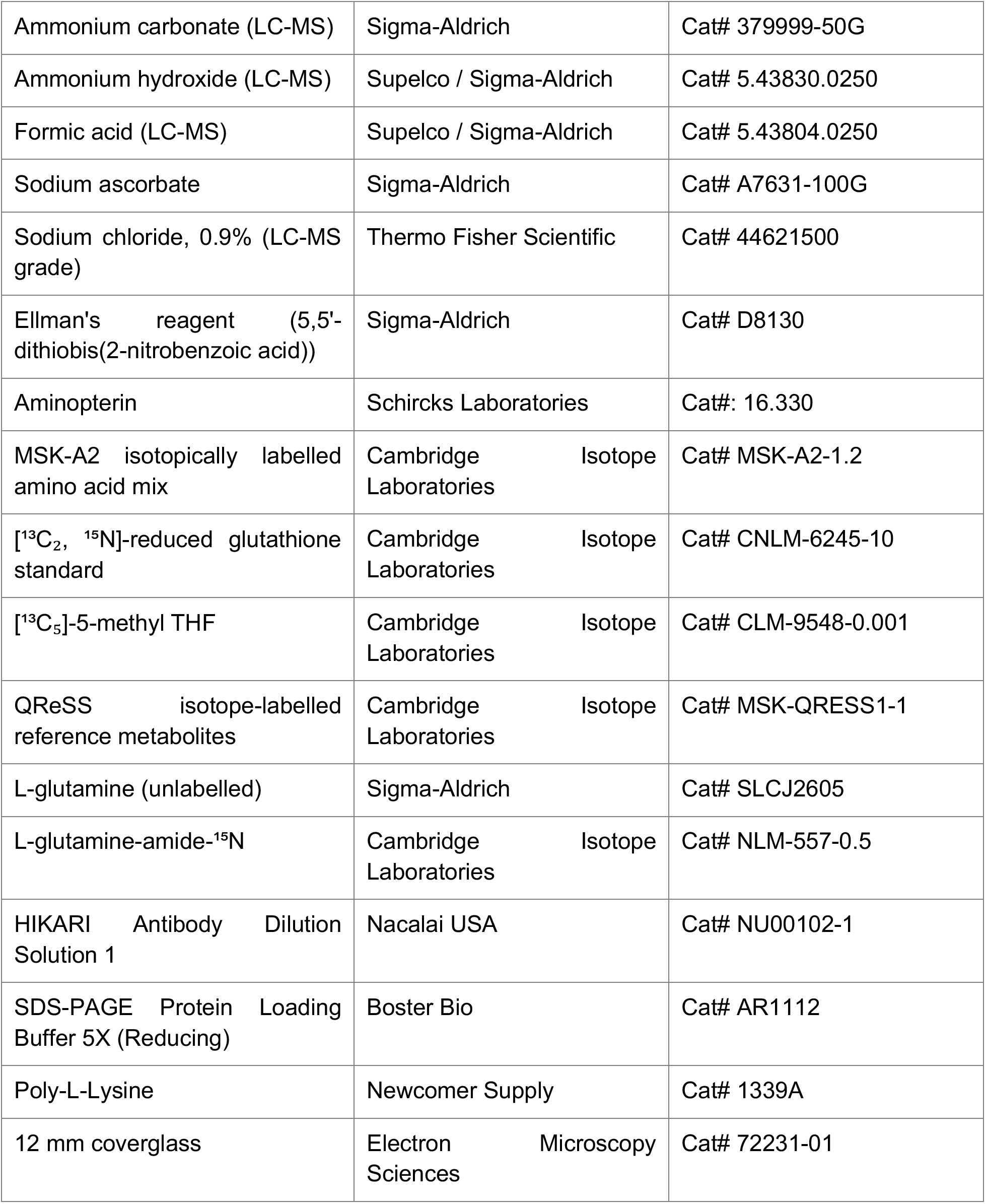
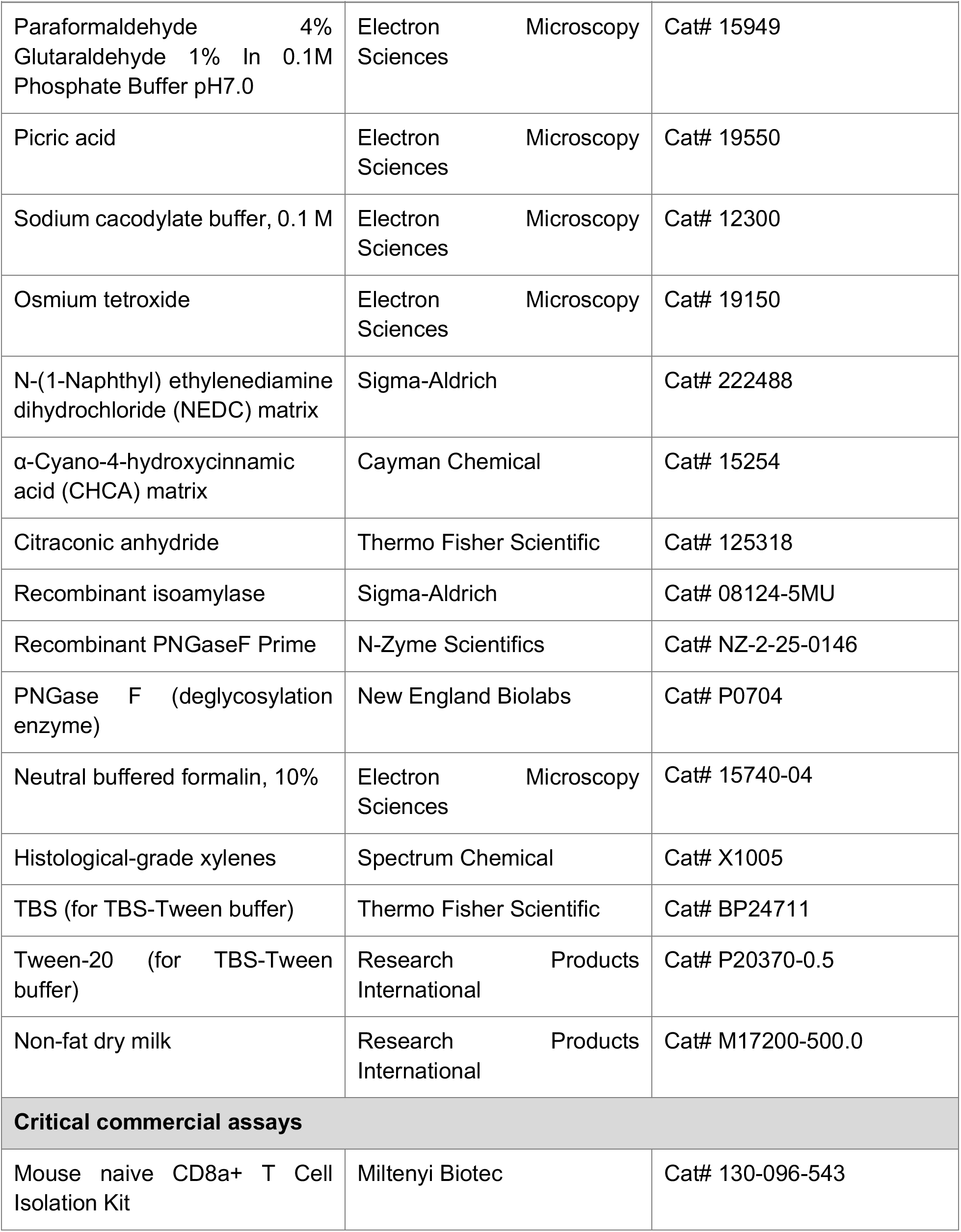
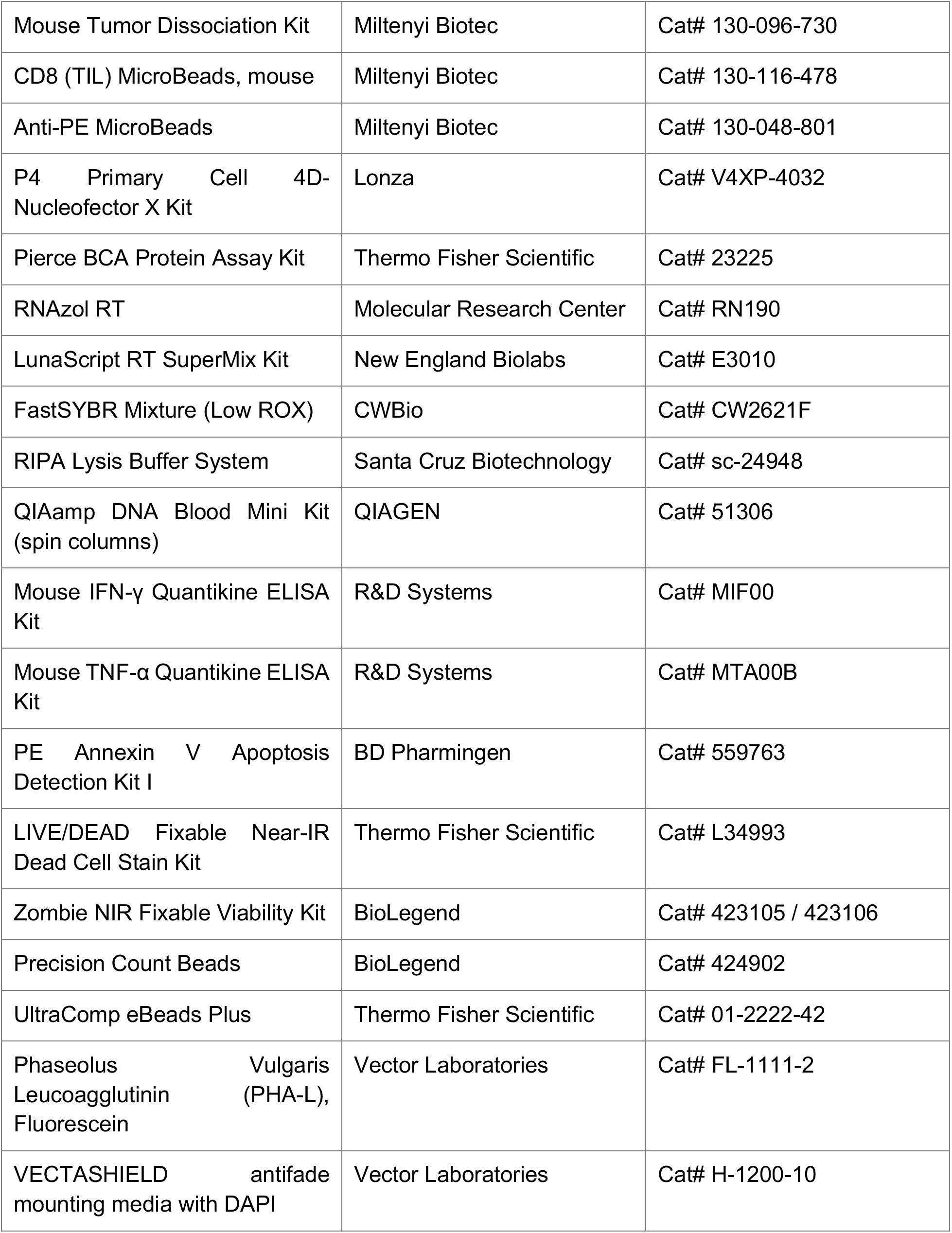
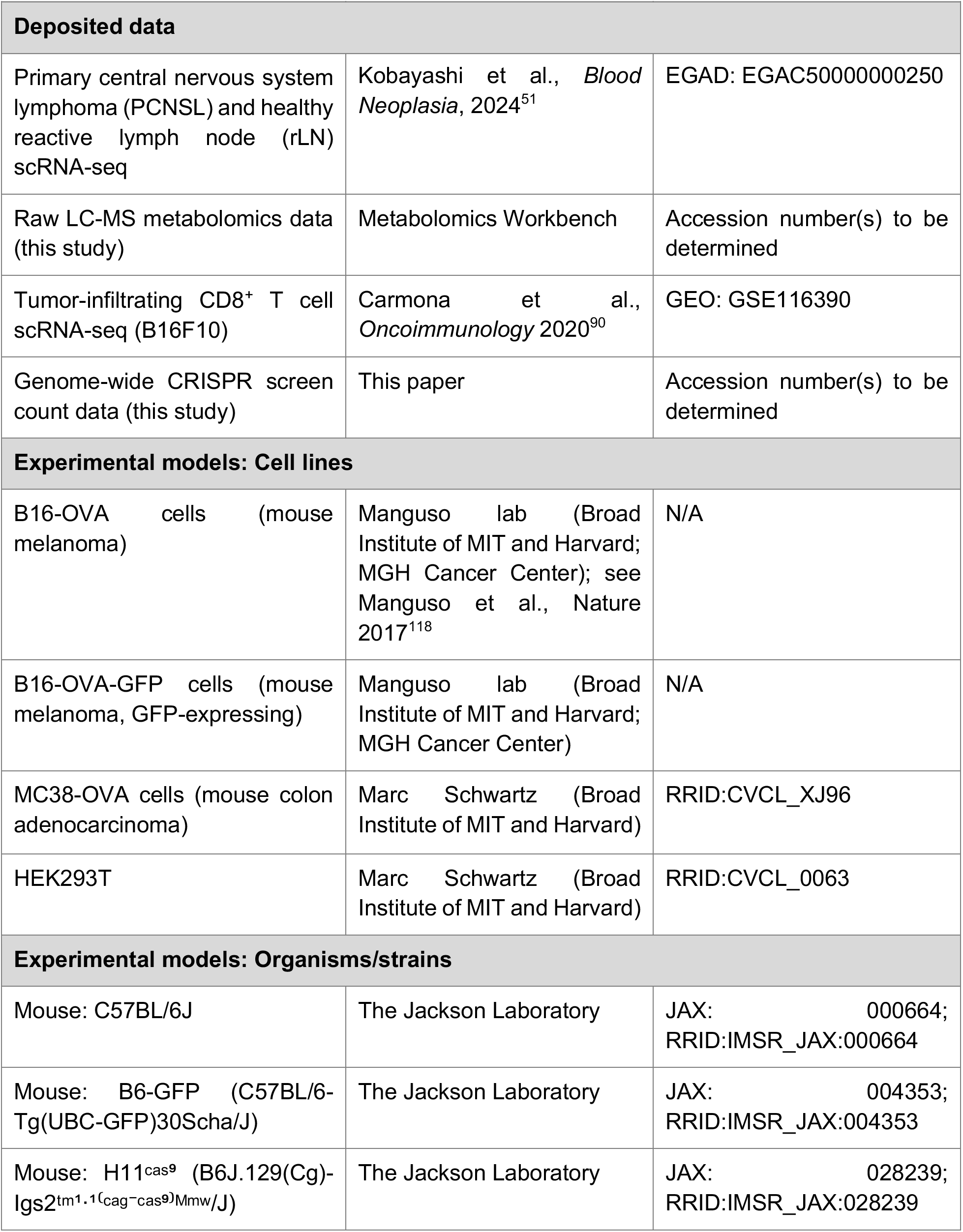
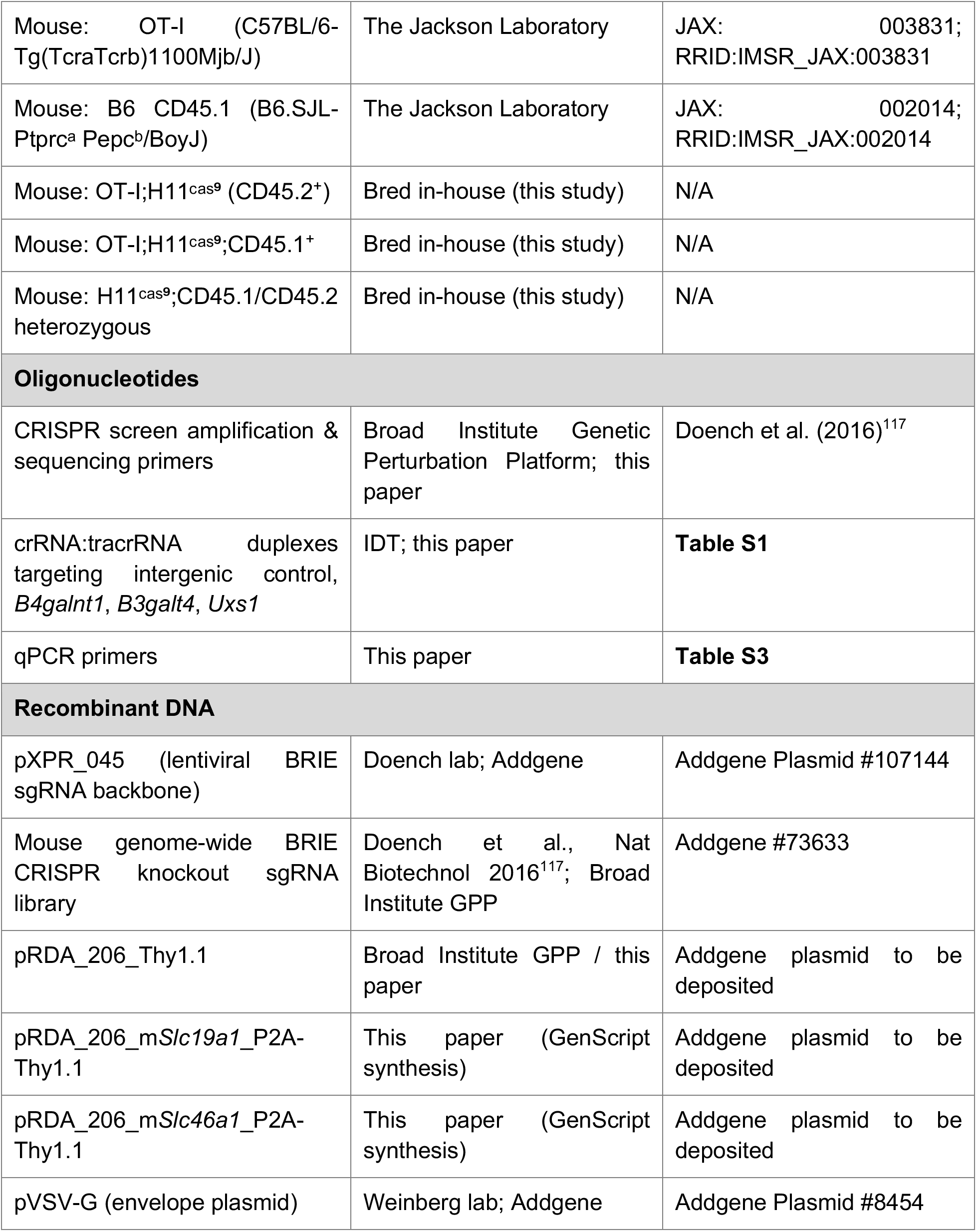
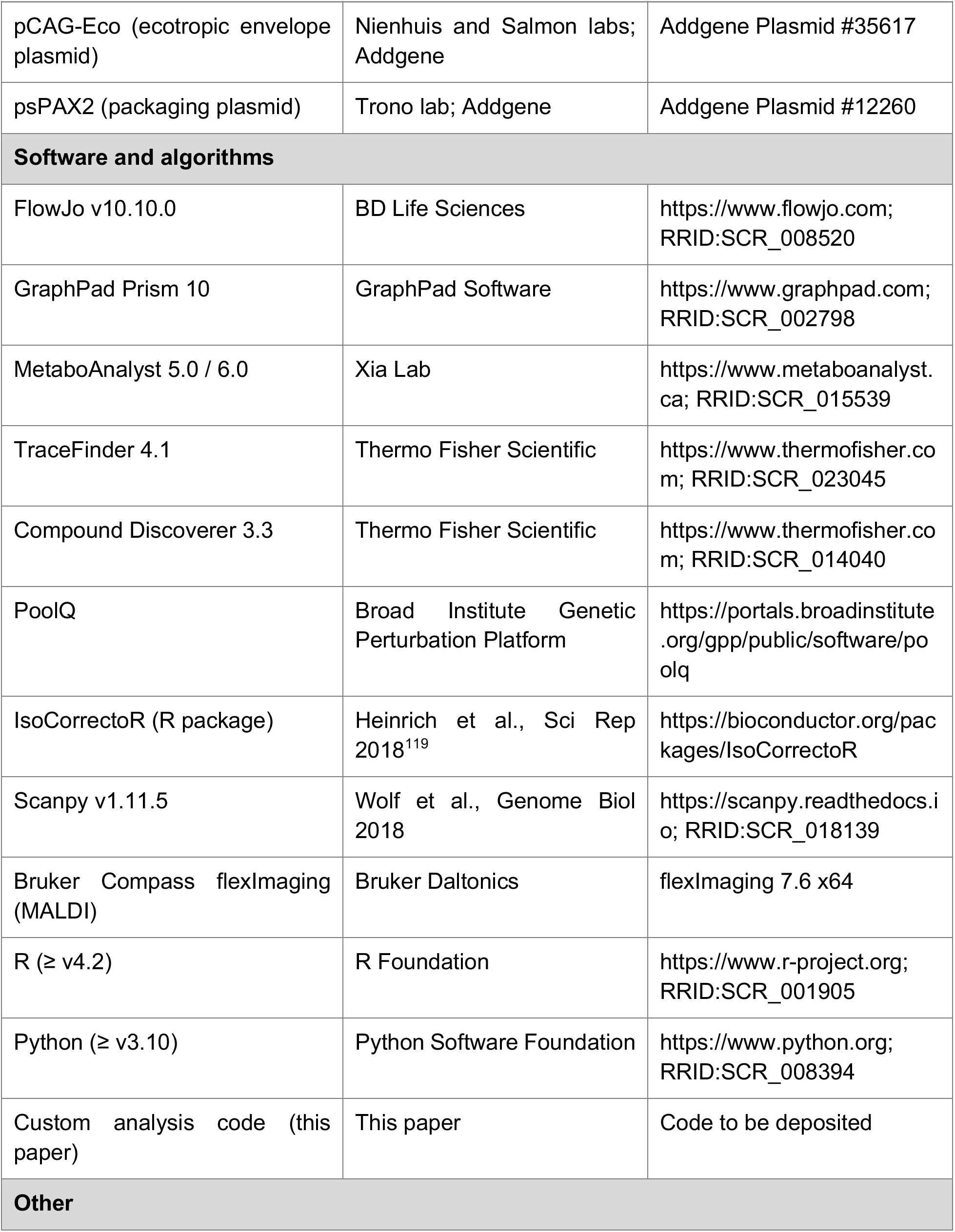
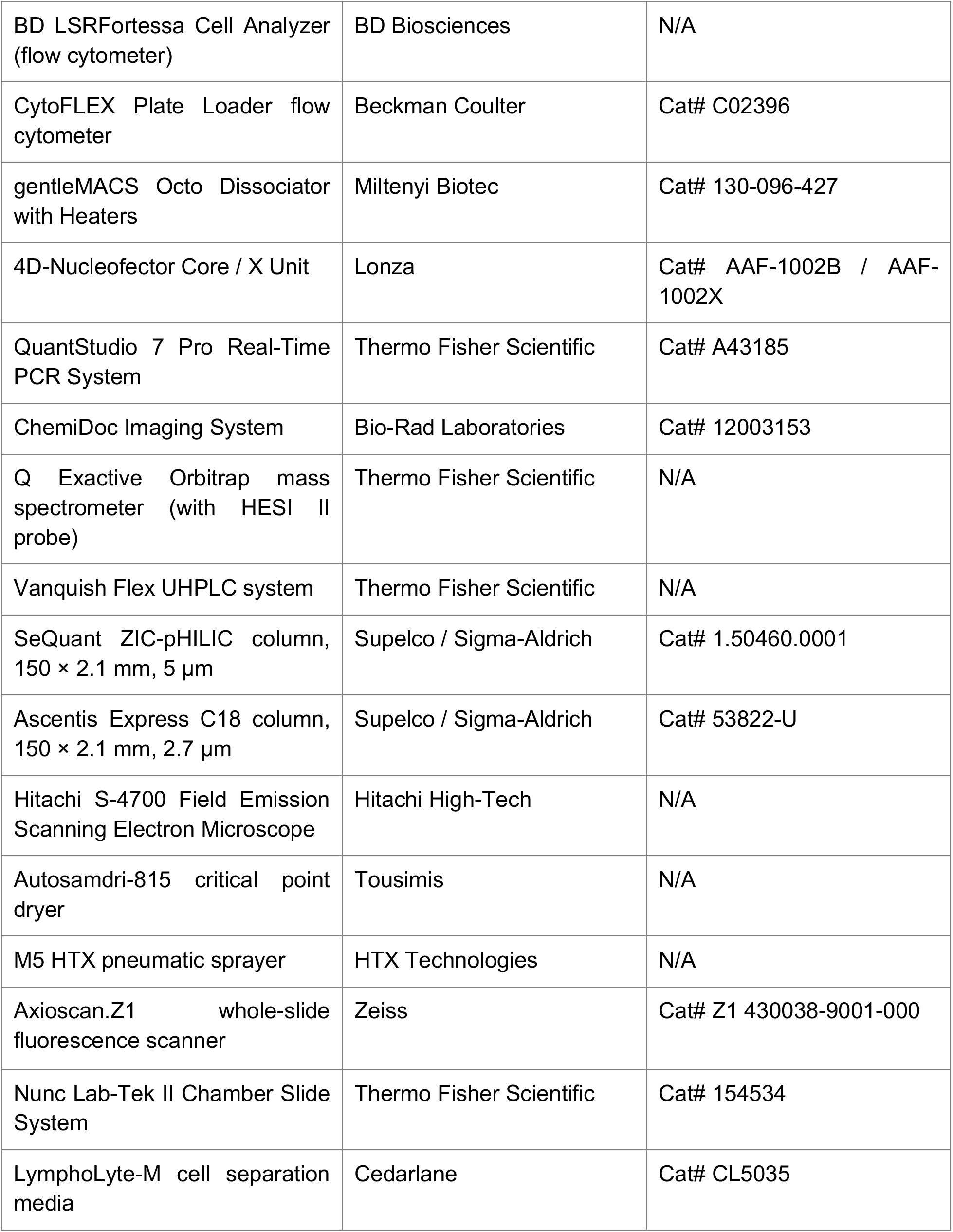
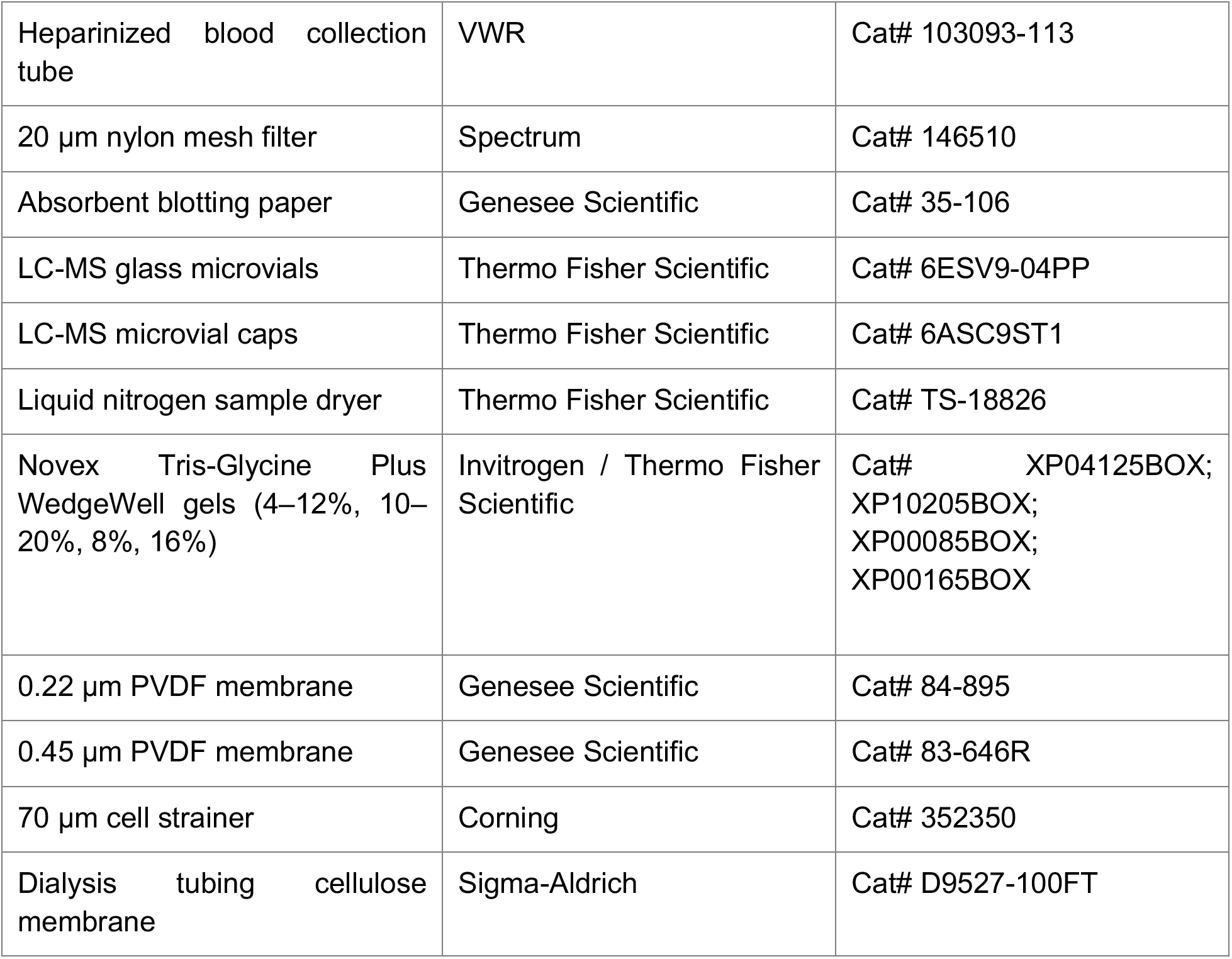

## METHODS

### Mice

All animal studies were reviewed and approved by the Institutional Animal Care and Use Committees (IACUCs) of Boston Children’s Hospital and the Broad Institute of MIT and Harvard. Mice were housed under specific pathogen-free (SPF) conditions on a 12 h light / 12 h dark cycle with ad libitum access to food and water.

Wild-type C57BL/6J (JAX 000664), B6-GFP (C57BL/6-Tg(UBC-GFP)30Scha/J; JAX 004353), H11^Cas^^9^ (B6J.129(Cg)-*Igs2*^tm1.1(CAG-Cas9*)Mmw^/J; JAX 028239), OT-I (C57BL/6-Tg(*Tcra/Tcrb*)1100Mjb/J; JAX 003831), and B6 CD45.1 (B6.SJL-*Ptprc^a^ Pepc^b^*/BoyJ; JAX 002014) mice were purchased from The Jackson Laboratory and bred in-house. Both male and female mice aged 6–16 weeks were used for all *ex vivo* T cell isolation and tumor experiments, unless otherwise specified.

For adoptive transfer experiments, congenically marked OT-I;H11^Cas9^ mice were generated in-house by first crossing H11^Cas9^ knock-in mice (JAX 028239) with OT-I mice. OT-I;H11^Cas9^;CD45.1⁺ animals were subsequently generated by crossing this line with B6 CD45.1 mice. Female donors (OT-I;H11^Cas9^;CD45.2⁺/CD45.2⁺ or CD45.1⁺/CD45.1⁺, aged 13–38 weeks) were used as T cell donors for naïve CD8 T cell adoptive transfer. Congenically marked heterozygous H11^Cas9^;CD45.1⁺/CD45.2⁺ male and female mice (age-matched) were used as transfer recipients.

### Cell lines

B16-OVA and B16-OVA-GFP (C57BL/6-derived murine melanoma overexpressing chicken ovalbumin) cells were kindly provided by Dr. Robert Manguso (Broad Institute of MIT and Harvard; Massachusetts General Hospital Cancer Center; Manguso et al., *Nature* 2017). HEK293T and MC38-OVA (C57BL/6-derived murine colon adenocarcinoma stably expressing OVA) cells were kindly provided by Dr. Marc Schwartz (Broad Institute of MIT and Harvard). All cell lines were maintained at 37 °C in 5% CO₂ and were routinely tested for *Mycoplasma* contamination (all negative throughout the study).

B16-OVA and B16-OVA-GFP cells were cultured in DMEM (GenClone 25-500) supplemented with 10% fetal bovine serum (FBS; GenClone, 25-550 or Sigma-Aldrich, 12306C-500ML) and 1% penicillin–streptomycin (GenClone, 25-512). HEK293T and MC38-OVA cells were cultured in DMEM with 10% FBS, 10% GlutaMAX (Gibco, 35050061), and 1% penicillin– streptomycin. MC38-OVA cells carry an OVA-expressing vector conferring hygromycin resistance (the hygromycin-resistance cassette is encoded on the OVA vector); cells were selected with 125 µg mL⁻¹ hygromycin B for two consecutive days beginning at least three days prior to *in vivo* injection. For lentivirus production, HEK293T cells were cultured in DMEM supplemented with 10% FBS, 10% GlutaMAX, 1% MEM non-essential amino acids, 1% penicillin–streptomycin, and 1 g per 100 mL lyophilized BSA.

Primary murine CD8 T cells were maintained in complete T cell medium: RPMI-1640 (Sigma-Aldrich, R1145-500ML; diluted to 1×, without L-glutamine, folic acid (FA), or sodium bicarbonate) supplemented with 10% FBS, 55 µM β-mercaptoethanol (Gibco, 21985023), 25 mM HEPES (Corning, 25-060-CI), 1 mM sodium pyruvate (Corning, 25-000-CI or Gibco, 11360070), 1% MEM non-essential amino acids (Sigma-Aldrich, M7145-100ML), 1% GlutaMAX (Gibco, 35050061), 2 g L⁻¹ sodium bicarbonate (Corning, 25-035-CI or Sigma-Aldrich, S8761-100ML), 1% penicillin–streptomycin (GenClone, 25-512), and 2,000 nM folic acid (Sigma-Aldrich, F8758). For *in vitro* folate deprivation experiments, FBS in the T cell medium was replaced with in-house dialyzed FBS (dialysis against 1× PBS using cellulose membrane tubing with a 14 kDa molecular weight cutoff (Sigma-Aldrich, D9527), with four buffer exchanges of 4 L each at 12-hour intervals over 48 hours at 4 °C, followed by filter sterilization), and folic acid was added back to reach final concentrations of 2,000, 200, 100, 50, 20, or 0 nM. For nucleoside supplementation experiments, 100 µM inosine (Sigma-Aldrich, I4125-10G) and/or 100 µM uridine (Sigma-Aldrich, U3750-25G) were added to the medium.

## METHOD DETAILS

### Isolation and culture of primary murine CD8 T cells

Naïve murine CD8 T cells were isolated from the spleens of 6–16 week-old male and female mice. Spleens were gently filtered through a 70 µm cell strainer using the flat end of a 10 mL syringe plunger into a 6 cm cell culture dish on ice. Splenocytes were eluted from the strainer with 13 mL of ice-cold, 0.22 µm sterile-filtered PBE buffer (1× PBS with 2% FBS and 2 mM EDTA; GenClone, 25-507XB; Corning, 46-034-CI). The strainer and dish were each washed with an additional 6 mL PBE. Splenocytes were pelleted (350 × g, 5 min, 4 °C), resuspended in 1 mL per spleen of ACK lysing buffer (Gibco, A1049201), vortexed, and incubated for 2–3 min at room temperature. The lysed cell suspension was diluted with 10 mL PBE per spleen, filtered through a second 70 µm strainer (washed with another 10 mL PBE), and centrifuged again (350 × g, 5 min, 4 °C). Naïve CD8 T cells were purified using the Mouse Naive CD8a⁺ T Cell Isolation Kit (Miltenyi Biotec, 130-096-543) per the manufacturer’s instructions.

For *in vitro* assays, naïve CD8 T cells were activated on tissue culture plates precoated with anti-CD3ε (InVivoMAb clone 145-2C11, Bio X Cell, BE0001-1) and anti-CD28 (InVivoMAb clone 37.51, Bio X Cell, BE0015-5). Coating was performed by adding 10 µg/mL anti-CD3ε and 5 µg/mL anti-CD28 in sterile 1XPBS to either non-TC treated 48-well (100 µL/well) or 6-well plates (1 mL/well); incubation was either minimum 2 hours at 37°C, or overnight at 4°C wrapped with parafilm. Cells were activated on the coated plates for 48 h (day −7 to day −5) and then expanded in T cell medium supplemented with 200 IU mL⁻¹ recombinant mouse IL-2 (rmIL-2; BioLegend, 575406), with medium refreshed every 2 d. For cytotoxicity assays, 10 ng mL⁻¹ rmIL-12 (PeproTech, 210-12) was additionally included during expansion. Five days after purification (day −2), T cells were split into media containing the indicated folic acid concentrations. Seven days after purification (day 0), T cells were restimulated on freshly coated anti-CD3ε/anti-CD28 plates; after 24 h (day 1), cells were lifted and expanded in T cell medium with rmIL-2.

### In vitro killing assay

Naïve OT-I CD8 T cells were isolated from the spleens of 6–16 week-old male and female OT-I mice using the Mouse Naive CD8a⁺ T Cell Isolation Kit (Miltenyi Biotec, 130-096-543) as described above, and activated on anti-CD3ε/anti-CD28-coated plates for 48 h (day −7 to day −5). Cells were then expanded in T cell medium supplemented with 200 IU mL⁻¹ rmIL-2 (BioLegend, 575406) and 10 ng mL⁻¹ rmIL-12 (PeproTech, 210-12). On day −3 (four days after purification), OT-I cells were split into media containing the indicated folic acid concentrations. Seven days after purification, OT-I cells were co-cultured with B16-OVA-GFP target cells that had been pre-stimulated with 50 U mL⁻¹ IFN-γ (PeproTech, 315-05) for 48 h, at an effector:target ratio of 1:6. After 48 h of co-culture, cells were harvested and analyzed by flow cytometry to quantify antigen-specific target cell killing. Precision Count Beads (BioLegend, 424902) were used to normalize absolute cell counts.

### Flow cytometry

Flow cytometry was performed on a BD LSRFortessa (BD Biosciences) or a CytoFLEX with Plate Loader (Beckman Coulter, C02396). Data were analyzed with FlowJo v10.10.0. Briefly, cells were washed in FACS buffer (PBS with 2% FBS and 2 mM EDTA), then incubated with the indicated fluorophore-conjugated antibodies and live/dead stain for 30 min at room temperature in the dark. After staining, cells were washed twice in FACS buffer and acquired. Unless otherwise indicated, all antibodies were used at a 1:200 dilution. For tumor-infiltrating lymphocyte (TIL) staining, cells were first incubated with TruStain FcX PLUS (anti-CD16/32, BioLegend, 156604) at a 1:100 dilution, then stained with cell-surface antibodies at a 1:400 dilution. A complete antibody list, with fluorophores and catalogue numbers, is provided in the Key Resources Table. Compensation was performed with UltraComp beads (Thermo Fisher Scientific, 01-2222-42). Apoptosis was assessed with the PE Annexin V Apoptosis Detection Kit I (BD Pharmingen, 559763). Cell viability was determined using the LIVE/DEAD Fixable Near-IR Dead Cell Stain Kit (Thermo Fisher Scientific, L34993) or Zombie NIR Fixable Viability Kit (BioLegend, 423105/423106).

### Syngeneic tumor models

Mice were anaesthetized with 3% isoflurane and received subcutaneous inoculations of 1 × 10⁶ MC38-OVA or 2.5 × 10⁵ B16-OVA cells into the right flank, using a 1 mL insulin syringe (Becton Dickinson, 309659) with a 23G needle (Becton Dickinson, 305194). Sterile PBS was administered as a vehicle control. Tumors were measured with digital calipers every 2–3 d once palpable, and tumor volume was calculated using the formula V = (W² × L)/2, where W is width and L is length. Median tumor volume was 1,189–1,555 mm^3^ for both models. Mice were euthanized 21 d after injection or earlier, when tumor volume reached ∼1.5–2 cm³ or at pre-defined analysis time points as described in the figure legends.

### Genome-wide CRISPR screen

Naïve CD8 T cells isolated from 28 OT-I;H11^Cas9^ donor mice (approximately 200 × 10⁶ cells total) were transduced with the mouse genome-wide BRIE CRISPR knockout sgRNA library^117^ cloned into the lentiviral vector pXPR_045 (Addgene, 107144). Following lentivirus titration directly on primary T cells in 6-well plates, 2 mL of concentrated lentivirus per well was selected as the optimal volume to maximize transduction efficiency while maintaining library representation. Five days post-transduction, Thy1.1⁺ (transduced) cells were enriched by magnet-activated cell sorting (MACS) using anti-PE microbeads (Miltenyi Biotec, 130-048-801).

Fifty million purified cells were resuspended in T cell medium supplemented with 10 ng mL⁻¹ IL-7 (PeproTech, 217-17), 10 ng mL⁻¹ IL-15 (PeproTech, 210-15), and 2 ng mL⁻¹ IL-2 (PeproTech, 212-12), and were stimulated overnight on plates precoated overnight at 4 °C with 5 µg mL⁻¹ each of anti-CD3ε (clone 145-2C11) and anti-CD28 (clone 37.51). Cells were then lifted and resuspended in T cell medium containing rmIL-7/IL-15/IL-2 adjusted to either high folate (2,000 nM) or low folate (50 nM) conditions. Fifty million cells per condition were collected on day 3 for genomic DNA (gDNA) isolation. An additional 50 × 10⁶ cells per condition were restimulated overnight on anti-CD3ε/anti-CD28-coated plates, lifted, and returned to their respective folate conditions. Cells were maintained until day 7 post-initial activation, at which time 117 × 10⁶ cells were recovered from the high-folate condition and 31.6 × 10⁶ cells from the low-folate condition. gDNA was isolated using QIAamp Mini spin columns (QIAGEN, 51306) per the manufacturer’s instructions.

sgRNA cassettes were PCR-amplified with indexed primers as described in Doench et al. (2016)^117^, with cycle numbers empirically determined by qPCR to ensure uniform amplification across all samples. PCR products were purified by double-sided SPRI size selection (0.6×/1.2× AMPure XP), quantified by Qubit and Bioanalyzer, pooled in equimolar ratios, and sequenced on an Illumina NextSeq using custom sequencing primers. Log-normalized sgRNA abundances were quantified using PoolQ (https://portals.broadinstitute.org/gpp/public/software/poolq); downstream hit calling was performed using the STARS algorithm at FDR < 10%, as described in Doench et al. (2016)^117^.

### CRISPR screen hit validation

Validation of top CRISPR screen hits was performed by nucleofection of crRNA:tracrRNA ribonucleoprotein (RNP) duplexes into H11^Cas9^ CD8 T cells essentially as described previously^120^. Freshly isolated naïve H11^Cas9^ CD8 T cells were cultured overnight in T cell medium with 5 ng mL⁻¹ mouse IL-7 (BioLegend, 57802). The following day, 2 × 10⁶ CD8 T cells per reaction were washed with PBS, resuspended in P4 Primary Cell nucleofection solution (Lonza, V4XP-4032), and mixed with a pool of four pre-assembled crRNA:tracrRNA duplexes (Basic Protocol 3 from Seki and Rutz, 2018) targeting intergenic control, *B4galnt1*, *B3galt4*, or *Uxs1* (guide sequences in **Supplementary Table 1**). Nucleofection was performed on a Lonza 4D Nucleofector (AAF-1002B / AAF-1002X) using the Primary Cell P4 preset with pulse code DS137. Cells were allowed to recover for at least 2 h in T cell medium, activated on anti-CD3ε/anti-CD28-coated plates for 2 d, and expanded with rmIL-2 for a further 3 d.

To competitively assess gene-specific phenotypes, nucleofected H11^Cas9^ T cells were mixed 1:1 with GFP-expressing CD8 T cells (isolated and cultured in parallel under identical conditions but without nucleofection), split into 2,000 nM or 50 nM folate media, and analyzed by flow cytometry to quantify the fraction of knockout (GFP⁻) cells and cell-surface ganglioside levels.

### Plasmids and molecular cloning

Mouse *Slc19a1* and *Slc46a1* overexpression cassettes were synthesized by GenScript and cloned into the lentiviral vector pRDA_206_Thy1.1, using a P2A self-cleaving peptide to link the transporter coding sequence to a Thy1.1 (CD90.1) reporter. Plasmids used in this study are listed in the Key Resources Table and include: pXPR_045 (Addgene, 107144); pRDA_206_Thy1.1 (Addgene plasmid ID upon deposition); pRDA_206_m*Slc19a1*_P2A-Thy1.1 (Addgene plasmid ID upon deposition); and pRDA_206_m*Slc46a1*_P2A-Thy1.1 (Addgene plasmid ID upon deposition). Full plasmid maps and sequences will be made available on Addgene upon publication.

### Lentivirus production and transduction of naïve CD8 T cells

HEK293T cells were co-transfected with pVSV-G, pCAG-Eco, psPAX2, and either the *Slc19a1* or *Slc46a1* overexpression construct using Lipofectamine 3000 (Invitrogen, L3000008) in Opti-MEM I Reduced Serum Medium (Gibco, 51985034) per the manufacturer’s instructions. Viral supernatants were harvested, clarified, and titered on HEK293T cells to determine the optimal transduction volume for primary CD8 T cells.

For CD8 T cell transduction, lentiviral supernatants were added to non-tissue culture plates precoated with 20 µg mL⁻¹ RetroNectin (Takara Bio, T100B) and blocked with 2% BSA in PBS. Plates were centrifuged at 2,000 × g for 2 h at 32 °C to immobilize virus onto the plate surface. Freshly isolated naïve CD8 T cells were then layered onto the RetroNectin/lentivirus-coated plates (following centrifugation, 4 mL of lentiviral supernatant was removed from the 5 mL total volume, and 4 mL of T cell suspension was added directly onto the virus-coated surface) in T cell medium supplemented with 10 ng mL⁻¹ rmIL-7 (PeproTech, 217-17), 10 ng mL⁻¹ rmIL-15 (PeproTech, 210-15), and 1:100 LentiBOOST (SIRION Biotech, SB-A-LF-901-01). After 48 h, cells were lifted, counted, and transferred to flasks in T cell medium with rmIL-7/rmIL-15 for an additional 5 d prior to enrichment of transduced cells and adoptive transfer.

### Adoptive T cell transfer

Prior to transfer, transduced (Thy1.1⁺) naïve CD8 T cells were stained with PE-conjugated anti-Thy1.1 antibody and further enriched using anti-PE MicroBeads (Miltenyi Biotec, 130-048-801) on MACS columns. Transduced Thy1.1⁺ CD45.1⁺ naïve CD8 T cells (expressing either m*Slc19a1* or m*Slc46a1*) were mixed 50:50 with CD45.2⁺ naïve CD8 T cells transduced with a non-targeting control construct, and mixing was confirmed by flow cytometry. Each recipient mouse received 3 × 10⁵ mixed T cells, resuspended in sterile PBS, by tail-vein injection.

### Tumor-infiltrating lymphocyte (TIL) isolation

Tumors were excised from euthanized mice and dissociated using the Mouse Tumor Dissociation Kit (Miltenyi Biotec, 130-096-730) with the gentleMACS Octo Dissociator (Miltenyi Biotec, 130-096-427), following the manufacturer’s tough tumor program. Dissociated cell suspensions were passed through a 70 µm cell strainer. Lymphocytes were then enriched by density-gradient centrifugation over LymphoLyte-M (Cedarlane, CL5035) per the manufacturer’s protocol. CD8⁺ TILs were further enriched using CD8 (TIL) MicroBeads (Miltenyi Biotec, 130-116-478) and MACS. TIL counts and viability were assessed, and cells were stained for flow cytometry as described above.

### Plasma, tumor interstitial fluid, and lymph-node extracellular fluid isolation

Whole blood was collected from awake mice by submandibular bleed into heparinized collection tubes (VWR, 103093-113) and centrifuged at 2,000 × g for 10 min at 4 °C. Plasma supernatants were collected and used immediately for LC-MS metabolite profiling.

For tumor interstitial fluid (TIF), tumors were excised from mice anaesthetized with ketamine/xylazine (120/10 mg kg⁻¹, i.p.), briefly rinsed in ice-cold PBS, blotted on absorbent paper (Genesee Scientific, 35-106), and placed on 20 µm nylon mesh filters (Spectrum, 146510) affixed to 50 mL conical tubes. Tubes were centrifuged at 106 × g for 10 min at 4 °C; the eluate (TIF) was collected and used immediately for LC-MS.

For lymph-node extracellular fluid (EF), tumor-draining inguinal and axillary lymph nodes (ipsilateral to the tumor) were excised from ketamine/xylazine-anaesthetized mice, briefly rinsed in ice-cold PBS, blotted, and bisected with sharp scissors. Due to sample volume limitations, tdLN EF were pooled within groups for LC-MS analysis (*n* = 3 pooled samples/group; *n* = 4 – 5 mice/pooled sample). 8–10 lymph nodes from 4–5 mice were pooled per biological replicate, placed on 20 µm nylon mesh filters atop 1.5 mL tubes, and centrifuged at 700 × g for 10 min at 4 °C. Tumor-draining lymph node EF was collected and used immediately for LC-MS metabolite profiling.

### LC-MS metabolite profiling

#### Sample preparation

For cultured CD8 T cells, 2 × 10⁶ cells were collected by centrifugation (21,300 × g, 20 s, 4 °C), washed with 1 mL of 0.9% NaCl (LC-MS grade), and resuspended in 300 µL of ice-cold folate extraction buffer [80% LC-MS methanol / 20% LC-MS water containing 25 mM ammonium acetate, 2.5 mM sodium ascorbate, supplemented with isotopically labelled amino acid standards (Cambridge Isotope Laboratories, MSK-A2-1.2), aminopterin, and the isotopically labelled reduced glutathione internal standard (Cambridge Isotope Laboratories, CNLM-6245-10)]. Samples were vortexed for 15 s to lyse cells and extract metabolites.

Fresh plasma, TIF, and tumor-draining lymph node (tdLN) EF were used for immediate LC-MS analysis after isolation. For relative quantification of 5-methyl-THF, 10 µL of plasma and 6–10 µL of TIF per mouse were extracted in 300 µL of extraction buffer [80% LC-MS methanol / 20% LC-MS water with 25 mM ammonium acetate, 2.5 mM sodium ascorbate, and 20 mM Ellman’s reagent (5,5′-dithiobis(2-nitrobenzoic acid), Sigma-Aldrich, D8130), supplemented with MSK-A2 amino acid standards, aminopterin, and glutathione standard]. For absolute quantification of 5-methyl-THF, 4 µL plasma, 4 µL TIF, and 1 µL of pooled tdLN EF per mouse were extracted in 150 µL of the same extraction buffer, and a 200 nM [¹³C₅]-5-methyl-THF internal standard (Cambridge Isotope Laboratories, CLM-9548-0.001) was spiked into each sample (final concentration 20 nM after drying and reconstitution). [¹³C₅]-5-methyl-THF calibration curves (6.25, 12.5, 25, 50, 100, 200, 400, and 800 nM) were prepared in non-spiked plasma and TIF matrices for absolute quantification.

After addition of extraction buffer, samples were vortexed for 10 s and centrifuged at 21,300 × g for 10 min at 4 °C to pellet debris. Supernatants were transferred to fresh 1.5 mL tubes and dried on ice using a liquid nitrogen sample dryer (Thermo Scientific, TS-18826). Dried T cell culture extracts were reconstituted in 15 µL LC-MS water supplemented with QReSS isotope-labelled reference metabolites (Cambridge Isotope Laboratories, MSK-QRESS1-1); dried mouse biological fluid extracts were reconstituted in 20 µL LC-MS water with QReSS, except tdLN EF extracts, which were reconstituted in 25 µL. Reconstituted samples were spun again at 21,300 × g for 10 min at 4 °C, and the cleared supernatants were transferred to LC-MS glass microvials (Thermo Fisher Scientific, 6ESV9-04PP, caps 6ASC9ST1). For each sample matrix, aliquots from every sample were combined into a matrix-matched pooled sample, which was then serially diluted 3- and 10-fold and injected throughout the randomized sequence for quality control and as ion-suppression/technical-drift controls.

#### Polar metabolite detection

1 µL of each sample (3 µL for tdLN EF) was injected onto a ZIC-pHILIC column (150 × 2.1 mm, 5 µm; EMD Millipore) on a Vanquish Flex UHPLC system (Thermo Fisher Scientific). One of two chromatographic gradients, which produced equivalent retention times, was used:

<u>Gradient 1:</u> Buffer A = 95% acetonitrile + 5% aqueous 20 mM ammonium carbonate, 0.1% ammonium hydroxide; Buffer B = 95% aqueous 20 mM ammonium carbonate, 0.1% ammonium hydroxide + 5% acetonitrile. Flow rate 0.150 mL min⁻¹; column oven 25 °C; autosampler 4 °C. 0–20 min: linear gradient from 16.6% to 83.4% B; 20–24 min: hold at 83.4% B; 24–24.1 min: 83.4% to 16.6% B; 24.1–32 min: hold at 16.6% B.

<u>Gradient 2:</u> Buffer A and Buffer B as in Gradient 1. Flow rate 0.150 mL min⁻¹; column oven 25 °C; autosampler 4 °C. 0–17.5 min: linear gradient from 16.6% to 75% B; 17.5–18.0 min: 75% to 85% B; 18–20 min: hold at 85% B; 20–20.5 min: 85% to 16.6% B; 20.5–24 min: hold at 16.6% B.

Mass spectrometry data were acquired on a Q Exactive benchtop Orbitrap mass spectrometer equipped with an Ion Max source and a heated electrospray ionization (HESI) II probe (Thermo Fisher Scientific), operated with polarity switching. Tune file parameters: spray voltage = 3.5 kV; capillary temperature = 320 °C; S-lens RF = 50; auxiliary gas temperature = 350°C.

For cell culture and ¹⁵N-glutamine tracing polar metabolomics, Gradient 1 and five scan events were used: (i) full scan in positive and negative ionization, m/z 70–1,000, resolution 70,000, AGC target 1 × 10⁶, max IT 20 ms from 0–20 min; (ii) full scan in positive and negative ionization, m/z 480–650, resolution 70,000, AGC target 1 × 10⁶, max IT 20 ms from 4–14 min; (iii) full scan in positive ionization, m/z 1,000–1,100, resolution 70,000, AGC target 5 × 10⁴, max IT 10 ms from 22.5–32 min.

For 5-methyl-THF relative quantification polar metabolomics, Gradient 1 and five scan events were used: (i) full scan in positive and negative ionization, m/z 70–1,000, resolution 70,000, AGC target 1 × 10⁶, max IT 40 ms from 0–20 min; (ii) full scan in negative ionization, m/z 220–700, resolution 35,000, AGC target 1 × 10⁶, max IT 20 ms from 0–20 min; (iii) full scan in positive ionization, m/z 1,000–1,100, resolution 70,000, AGC target 5 × 10⁴, max IT 10 ms from 22.5–32 min; (iv) targeted-SIM in negative ionization, isolation window 1.0 m/z, resolution 35,000, AGC target 1 × 10⁵, max IT 20 ms from 0–20 min.

For 5-methyl-THF absolute quantification polar metabolomics, Gradient 2 and two scan events were used: full scans in positive and negative ionization, m/z 70–1,000, resolution 70,000, AGC target 1 × 10⁶, max IT 40 ms from 0–17.5 min.

#### Folate metabolite detection (LC-MS and LC-MS/MS)

5 µL of each sample (18 µL for tdLN EF) was injected onto a Supelco Ascentis Express C18 column (150 × 2.1 mm, 2.7 µm; EMD Millipore) on a Vanquish Flex UHPLC system (Thermo Fisher Scientific). One of two gradients was used:

<u>Gradient 1:</u> Buffer A = LC-MS water with 0.1% formic acid; Buffer B = acetonitrile with 0.1% formic acid. Flow rate 0.250 mL min⁻¹; column oven 30 °C; autosampler 4 °C. 0–2 min: hold 5% B; 2–10 min: linear gradient 5% to 36% B; 10–10.1 min: 36% to 95% B; 10.1–14 min: hold 95% B; 14–14.1 min: 95% to 5% B; 14.1–18 min: hold 5% B.

<u>Gradient 2:</u> Buffer A and Buffer B as in Gradient 1. Flow rate 0.450 mL min⁻¹; column oven 30 °C; autosampler 4 °C. 0–1 min: hold 3% B; 1–5 min: linear gradient 3% to 55% B; 5–5.3 min: 55% to 95% B; 5.3–6 min: hold 95% B; 6–6.3 min: 95% to 3% B; 6.3–7.5 min: hold 3% B.

Mass spectrometry was performed on a Q Exactive benchtop Orbitrap equipped with an Ion Max source and HESI II probe (Thermo Fisher Scientific). HESI parameters: sheath gas flow rate = 40; auxiliary gas flow rate = 10; sweep gas = 0; spray voltage = 2.8 kV (negative) / 3.5 kV (positive); capillary temperature = 300 °C; S-lens RF = 50; auxiliary gas heater = 350 °C.

For 5-methyl-THF relative quantification, Gradient 1 was used and MS was operated in full-scan positive-ionization mode with three narrow-range scans: m/z 438–450; m/z 452–462; and m/z 470–478 (resolution 70,000; AGC target 1 × 10⁶; max IT 150 ms; 1–12 min). Folate signals were normalized to aminopterin as an internal standard and to measured polar metabolites.

For 5-methyl-THF absolute quantification, Gradient 2 was used. MS was operated in positive-ionization mode using two scans for MS¹ and MS² spectra: (i) full scan m/z 100–1,000, resolution 70,000, AGC target 3 × 10⁶, max IT 50 ms from 0–5.5 min; (ii) a parallel reaction monitoring (PRM) scan with resolution 35,000, AGC target 1 × 10⁵, max IT 100 ms, in-source CID = 0.0 eV, microscans = 1, loop count = 6, MSX count = 1, MSX isochronous ITs = on, fixed first mass = 100.0 m/z, isolation window = 1.0 m/z, and isolation offset = 0.0 m/z from 0–5.5 min. Folate signals were normalized to aminopterin and [¹³C₅]-5-methyl-THF internal standards and to measured polar metabolites.

### ¹⁵N-glutamine isotope tracing

¹⁵N-glutamine tracing was performed in folate- and glutamine-free T cell medium prepared with dialyzed FBS (prepared as described above), supplemented with 2,000 nM or 0 nM folic acid and either unlabelled L-glutamine (Sigma-Aldrich, SLCJ2605) or L-glutamine-amide-¹⁵N (Cambridge Isotope Laboratories, NLM-557-0.5). A 25× stock solution of labelled and unlabelled glutamine (7.5 g L⁻¹) was prepared in sterile Milli-Q water prior to addition. Prior to tracing, cells were pre-conditioned for 4 h in folate- and glutamine-free T cell medium supplemented with unlabelled glutamine and 2,000 nM or 0 nM folic acid. Tracing was performed on primary murine CD8 T cells that had been cultured in 2,000 nM or 0 nM folic acid since day −2. On days 0, 1, and 3, cells were seeded into folate- and glutamine-free T cell medium supplemented with 2,000 nM or 0 nM folic acid and either unlabelled or ¹⁵N-amide-labelled glutamine. After 4 h of tracing, cells were washed in 0.9% NaCl and extracted in 300 µL of 80% methanol / 20% water (LC-MS grade) containing 25 mM ammonium acetate, 2.5 mM sodium ascorbate, and the isotopic standards described above. Chromatography and mass spectrometry data acquisition were performed as described for polar metabolites.

Relative quantification of polar metabolites was performed in TraceFinder 4.1 as described below. Expected masses for ¹⁵N-labelled species were calculated by adding the mass of a single extra neutron (0.997 Da) per labelled nitrogen to the m/z of each unlabelled target, and relative quantification of these masses yielded percent labelling for each nitrogen position (M+1, M+2, M+3,…). Natural ¹⁵N abundance was corrected using the R package IsoCorrectoR^119^. Corrected labelling is presented either as the change in percent labelling between experimental conditions or as a fraction of the total abundance measured in parallel unlabelled samples.

### Metabolomics data analysis

Polar metabolites were relatively quantified against an in-house library of authentic chemical standards using TraceFinder 4.1 or Compound Discoverer 3.3 (Thermo Fisher Scientific), with a 5 ppm mass tolerance. Matrix-matched pooled samples and serial dilutions of pooled samples were prepared as quality controls and were injected at the start, end, and at regular intervals throughout each run to assess technical drift and to calculate the coefficient of variation (CV) for each metabolite. Normalization was performed in R using scripts found at www.github.com/FrozenGas/KanarekLabTraceFinderRScripts.

Data normalization was performed in two steps. (i) *Technical normalization:* integrated peak areas for internal standards added to the extraction buffer were mean-centered (for each standard, peak area divided by the mean across samples) and averaged per sample; sample peak areas were divided by the resulting per-sample factor, normalizing for MS signal fluctuation and pipetting/injection error (typically within 10%). (ii) *Biological normalization:* CV values (from pooled sample reinjections) and coefficients of determination (R²) (from linear dilutions of pooled samples) were calculated per metabolite. Metabolites with CV < 30% and R² > 0.95 were mean-centered and averaged across samples, and peak areas were divided by the resulting biological normalizer to account for any global shift in metabolite amounts driven by differences in biological material. Median bridge normalization (Lippa et al., as described previously^121^) was used to correct for batch effects.

Visualization and additional statistical analysis of metabolomics data were performed with MetaboAnalyst 5.0 and 6.0.

### Immunoblot

Cells were lysed with the RIPA Lysis Buffer System (Santa Cruz Biotechnology sc-24948) per the manufacturer’s instructions. Whole-cell lysates were centrifuged at 10,000 × g for 10 min at 4 °C, and cleared supernatants were quantified using the Pierce BCA Protein Assay Kit (Thermo Fisher Scientific, 23225) for protein loading normalization. Lysates were denatured in 1× SDS-PAGE Protein Loading Buffer (5× Reducing; Boster Bio, AR1112) for 5 min at 85 °C. 20 µg of protein per lane was resolved on Invitrogen NuPAGE Bis-Tris gels (4–12%, 10–20%, 8%, or 16%, as indicated) and transferred onto a 0.22 µm (Genesee Scientific, 84-895) or 0.45 µm PVDF membrane (Genesee Scientific, 83-646R). Membranes were blocked for 1 h at room temperature in Tris-buffered saline with 1% Tween-20 (TBS-T) containing 5% non-fat milk.

Membranes were incubated overnight at 4 °C in HIKARI Solution 1 (Nacalai USA, NU00102-1) containing the following primary antibodies at 1:1,000: CD25 (CST, 39475), CD28 (CST, 38774), CD44 (CST, 37259), CHOP (ABclonal, A20987), PERK (CST, 3192), phospho-PERK (CST, 3179), cleaved caspase-3 (CST, 9661), caspase-3 (CST, 14220), or vinculin (CST, 13901). Membranes were washed three times in TBS-T, incubated for 1 h at room temperature in TBS-T / 5% milk containing HRP-linked anti-rabbit IgG (Jackson ImmunoResearch, 111-035-144) or HRP-linked anti-mouse IgG (Jackson ImmunoResearch, 111-035-166) at 1:10,000, and imaged with a ChemiDoc Imaging System (Bio-Rad, 12003153) or autoradiography.

For reprobing, membranes were stripped in a mild stripping buffer (200 mM glycine, 0.1% SDS, 1% Tween-20, pH 2.2; Abcam protocol) for 15 min at room temperature, then reblocked and reprobed as above. For enzymatic deglycosylation, 20 µg of protein in 1× loading buffer was treated with PNGase F (New England Biolabs, P0704) per the manufacturer’s instructions prior to SDS-PAGE. If a glycoprotein is N-glycosylated, then PNGase F treatment should remove all other glycoform bands, and a single downward-shifted band representing the core deglycosylated protein should remain on the blot. Other protein bands remaining after PNGase F treatment (i.e. hyper-glycosylated proteins appearing as higher band smears above the expected molecular weight band, or hypo-glycosylated proteins appearing as lower band smears below the expected molecular weight band) represent other modifications (e.g., O-glycosylation).

## ELISA

2 × 10⁶ CD8⁺ T cells were cultured in 1.5 mL of fresh medium per well in a 6-well plate for 3 h. Supernatants were collected, and secreted IFN-γ and TNF-α were quantified using the Mouse IFN-γ Quantikine ELISA Kit (R&D Systems, MIF00) and Mouse TNF-α Quantikine ELISA Kit (R&D Systems, MTA00B), respectively, following the manufacturer’s instructions.

### Scanning electron microscopy (SEM)

1 × 10⁶ CD8⁺ T cells in 1 mL medium were collected onto 12 mm poly-L-Lysine-coated coverslips (Newcomer Supply, 1339A; Electron Microscopy Sciences, 72231-01) by centrifugation (350 × g, 1 min, no brake). Samples were fixed overnight in 1.25% formaldehyde, 2.5% glutaraldehyde, and 0.03% picric acid in 0.1 M sodium cacodylate buffer (pH 7.4) (Electron Microscopy Sciences, 15949, 19550, and 12300), washed with 0.1 M sodium cacodylate buffer, and post-fixed in 1% osmium tetroxide ((Electron Microscopy Sciences, 19150)) in 0.1 M sodium cacodylate buffer for 2 h. Samples were rinsed in ddH₂O and dehydrated through a graded ethanol series (50%, 70%, 95%, and 100% × 2; 15 min each). Dehydrated samples were dried in an Autosamdri-815 critical point dryer, mounted on aluminum stubs with colloidal graphite, and sputter-coated with platinum (6 nm). Samples were imaged on a Hitachi S-4700 Field Emission Scanning Electron Microscope (FE-SEM) at an accelerating voltage of 5 kV.

### Gene expression analysis (RNA isolation, cDNA synthesis, and quantitative PCR)

2 × 10⁶ CD8⁺ T cells were collected from cell culture. Total RNA was extracted using RNAzol RT (Molecular Research Center, RN190) per the manufacturer’s instructions. RNA integrity and concentration were verified by A260/A280 NanoDrop measurement. 1 µg of RNA was reverse-transcribed using the LunaScript RT SuperMix Kit (New England Biolabs, E3010) per the manufacturer’s instructions. Quantitative PCR was performed on a QuantStudio 7 Pro Real-Time PCR System (Thermo Fisher Scientific, A43185) using FastSYBR Mixture (Low ROX; CWBio, CW2621F) per the manufacturer’s instructions. Primer sequences are listed in **Supplementary Table 3**. All samples were run in technical triplicate across 3 independent biological replicates. Relative expression was calculated by the 2^−ΔΔCq^ method with *Ubc* or *Hprt* as internal controls and the 2,000 nM folate condition as the reference.

### MALDI mass spectrometry imaging (MALDI-MSI) and analysis

HPLC-grade acetonitrile, ethanol, methanol, water, recombinant isoamylase, and α-cyano-4-hydroxycinnamic acid (CHCA) matrix were purchased from Sigma-Aldrich. Histological-grade xylenes were purchased from Spectrum Chemical. Citraconic anhydride for antigen retrieval was obtained from Thermo Fisher Scientific. Recombinant PNGaseF Prime was obtained from N-Zyme Scientifics (Doylestown, PA).

Cells were grown in Nunc Lab-Tek II Chamber Slides (Thermo Fisher Scientific, 154534) and pelleted by centrifugation. Slides were fixed in 10% neutral buffered formalin (37% formaldehyde in 1× PBS) for 1 h, dehydrated in 70% ethanol for 2 h, and then washed sequentially in 70%, 80%, 90%, and 100% ethanol (5 min each). Slides were desiccated overnight prior to antigen retrieval. Antigen retrieval was performed in 0.05% (v/v) citraconic anhydride (pH 3.0 ± 0.5) for 30 min at 95 °C, followed by vacuum desiccation.

PNGaseF was applied with an M5 HTX pneumatic sprayer. Three units of each enzyme was reconstituted in 1 mL HPLC-grade water and applied over 15 passes (0.025 mL min⁻¹, 3 mm offset, 900 mm min⁻¹ velocity, 45 °C nozzle temperature, ambient tray temperature). Slides were incubated at 37 °C for 2 hours at 100% relative humidity, vacuum desiccated, and then matrix-coated. 7 mg mL⁻¹ CHCA in 50% HPLC-grade acetonitrile was applied over 10 passes on the HTX M5 sprayer (0.1 mL min⁻¹, 3 mm offset, 1,200 mm min⁻¹ velocity, 79 °C nozzle, 50 °C heated tray).

Slides were imaged on a Bruker timsTOF fleX mass spectrometer. Acquisition and setup were performed in FlexImaging. High-resolution .jpg scans of the wells were uploaded to FlexImaging and manually aligned to the slide using three teachpoints, generating a .mis file with consistent masked ROIs.

Glycomics imaging was acquired in positive-ion MS¹ mode over m/z 600–4,000 using summation rate mode with a rate value of 544. Laser configuration: 50 × 50 µm pixels from a 46 × 46 µm raster range, 44.97% laser power, 1 burst of 396 shots, 10,000 Hz frequency. Transfer: 50 V MALDI plate offset, 70 V deflection 1 delta, 500 V_pp_ Funnel 1 RF, 500 V_pp_ Funnel 2 RF, 500 V_pp_ Multipole RF. Quadrupole: ion energy 5 eV, low-mass filter 500 m/z. Collision cell: collision energy 10 eV, collision RF 4,000 V_pp_. Focus Pre TOF: transfer time 180 µs; pre-pulse storage 25 µs.

### Human tissue samples

Formalin-fixed paraffin-embedded (FFPE) biopsy specimens from eight patients diagnosed with primary central nervous system lymphoma (PCNSL) were included in this study. Samples were obtained as part of routine clinical care and retrieved from institutional pathology archives following appropriate ethical approval. Paraffin blocks were sectioned using a microtome to generate consecutive tissue sections mounted on glass slides. Serial sections from each block were processed as follows:

● Slide 1: Hematoxylin and eosin (H&E) staining for histopathological evaluation and annotation of tumor regions.
● Slide 2: Negative control staining (secondary antibodies only, without primary antibodies).
● Slide 3: Multiplex immunofluorescence staining for CD8, ATF4, PHA-L, and DAPI.
● Slide 4: Multiplex immunofluorescence staining for CD20, ATF4, PHA-L, and DAPI.

### Multiplex immunofluorescence staining, slide scanning, and image acquisition

FFPE sections designated for immunofluorescence were deparaffinized in xylene and rehydrated through graded alcohols to distilled water. Antigen retrieval was performed using heat-induced epitope retrieval in citrate buffer. Following blocking of nonspecific binding sites, sections were incubated with primary antibodies against ATF4 (Cell Signaling Technology, 11815), CD8 (Dako, M7103), and CD20 (eBioscience, 14-0202-82). Glycan architecture was visualized using PHA-L lectin from *Phaseolus vulgaris* conjugated to GFP (Thermo Fisher Scientific, L11270). Primary antibodies were detected using fluorophore-conjugated secondary antibodies as follows: ATF4 was detected with an Alexa Fluor 647–conjugated secondary antibody (Cell Signaling Technology, 4414). CD8 and CD20 were detected using the same Alexa Fluor 555–conjugated secondary antibody (Thermo Fisher Scientific, A21422). Slides were mounted using antifade mounting medium containing DAPI (Vectashield, H1200) and coverslipped prior to imaging.

All slides were scanned at 20× magnification using a Zeiss Axioscan.Z1 whole-slide fluorescence scanner. Identical acquisition parameters were used across all samples to ensure comparability. Fluorescence channels corresponding to GFP (PHA-L), Cy3/Alexa 555 (CD8 or CD20), Cy5/Alexa 647 (ATF4), and DAPI were captured separately and exported as multichannel whole-slide images. Negative control slides were processed in parallel using identical staining and imaging procedures but with omission of primary antibodies to evaluate background fluorescence and nonspecific signal.

### Spatial image quantification

Quantitative image analysis was performed using QuPath (version 6.0). Regions of interest (ROIs) were manually annotated on the basis of H&E morphology and corresponding immunofluorescence signals to identify tumor-containing areas. Regions exhibiting strong autofluorescence were excluded using the negative control slides as reference.

Each ROI was subdivided into square spatial tiles of 300 µm × 300 µm to standardize spatial quantification. For each tile, mean fluorescence intensity was measured independently for each channel (GFP, Cy3, Cy5, and DAPI). To correct for background fluorescence, at least two adjacent regions outside the ROI were selected as background controls. Mean fluorescence intensity measured in these regions was subtracted from the corresponding ROI tile measurements. Identical thresholding and measurement parameters were applied across all samples to ensure consistent quantification.

### Multivariable spatial regression modeling and spatial gradient analysis

Whole-section spatial data were discretized into 300 or 400 µm square bins using spatial coordinates of segmented objects. For each bin, signal intensities for CD20, CD8, PHA-L, and ATF4 were quantified as the sum of normalized signal-to-noise ratios. Background regions were excluded from analysis. Within each tumor section, signals were z-normalized prior to spatial modeling to enable cross-sample comparison.

To assess independent spatial coupling between compartments, we fitted linear regression models across spatial bins within each tumor: Module_bin = β1 × CD8_bin + β2 × CD20_bin. Module_bin represents the PHA-L (glycan module) or ATF4 (stress module) signal intensity measured in each 400 µm spatial bin, and CD8_bin and CD20_bin represent the corresponding immune and tumor densities within the same bin.

The regression coefficients (β1 and β2) quantify the independent contribution of CD8 and CD20 densities to module signal intensity after mutual adjustment. Thus, β1 reflects the CD8-associated spatial effect independent of tumor density, while β2 reflects the CD20-associated spatial effect independent of immune density. Models were fitted separately and coefficients estimated independent for each tumor to capture patient-specific spatial architecture. Statistical significance of coefficients was assessed using standard Wald tests from linear regression models.

Spatial architectures were categorized according to the relative magnitude and direction of the regression coefficients in order to discriminate between immune-driven, tumor-driven, and hybrid spatial organization of glycan and stress modules:

● Immune-associated pattern: β_CD8 dominant (β1 > β2)
● Tumor-associated pattern: β_CD20 dominant (β2 > β1)
● Mixed pattern: both coefficients contribute substantially to module signal

To assess global directional alignment of spatial signals, principal spatial gradients were extracted for each marker by regressing signal intensity against spatial coordinates. Gradient vectors were compared across compartments using Spearman correlation. Correlation analyses were considered exploratory given the limited sample size (*n* = 5 tumors).

The CD8 Tumor Infiltration Index (TII) was defined as the fraction of CD20-high bins (upper quartile of CD20 density) that contained detectable CD8 signal. CD8 enrichment was calculated as the difference between mean CD8 intensity in CD20-high versus CD20-low bins. Module strength was represented by the CD8-associated regression coefficient (β1) from the multivariable model.

### Mouse scRNA-seq analysis

Raw count data for tumor-infiltrating CD8⁺ T cells from B16F10 subcutaneous melanoma tumors were obtained from the Gene Expression Omnibus (GEO accession GSE116390; Carmona et al., *Oncoimmunology* 2020^90^ in 10x Genomics Market Exchange Format (MEX). Data were processed with Scanpy (v1.11.5) in Python. Cells with fewer than 200 or more than 5,000 detected genes, or with mitochondrial transcript content exceeding 20%, were removed. Genes detected in fewer than 3 cells were excluded. A total of 3,574 cells and 16,527 genes passed quality control.

Library size was normalized by scaling each cell to a total count of 10,000, followed by log(1 + x) transformation. The top 2,000 highly variable genes were identified using the Seurat v3 method. Principal component analysis (PCA) was performed on scaled expression values (clipped at a maximum of 10), retaining the top 30 principal components. A shared nearest-neighbor graph was constructed using 15 neighbors and 30 principal components, followed by UMAP dimensionality reduction. Leiden graph-based clustering was applied at resolution 0.4, yielding 8 clusters.

Cluster marker genes were identified by Wilcoxon rank-sum test. Cell type identity was assigned manually based on canonical CD8⁺ T cell subset markers: *Tcf7*, *Sell*, and *Il7r* for Naïve/Stem-like cells; *Tcf7*, *Slamf6*, *Pdcd1*, and *Xcl1* for Progenitor-exhausted (Tpex) cells; *Xcl1* and *Xcl2* for an Xcl1⁺ Tpex subset; *Pdcd1*, *Havcr2*, *Lag3*, *Tox*, and *Tigit* for Terminally exhausted cells; *Gzmb*, *Prf1*, and *Nkg7* for NK-like/Cytotoxic cells; *Gzmk* and *Cx3cr1* for Effector/Tissue-resident memory cells; and *Mki67* and *Top2a* for Proliferating cells. One cluster characterized by expression of *Lyz2*, *C1qb*, and *Apoe* was identified as macrophage contamination and excluded from downstream visualization and analysis. Gene expression was visualized using dot plots with per-gene standard scaling and a sequential purple colormap. Figures were exported as PDF vector files with TrueType font embedding (pdf.fonttype = 42) for compatibility with Adobe Illustrator.

### Human PCNSL and TIL scRNA-seq analysis

scRNA-seq data were analyzed in R v4.3 using Seurat v5.3.0. Data from the European Genome-Phenome Archive (EGAD accession EGAC50000000250^51^) was derived from brain biopsies of 7 primary central nervous system lymphoma (PCNSL) patients and 2 reactive lymph nodes (rLN) biopsies of patients without cancer; these data were downloaded from Zenodo (https://zenodo.org/records/10656046) as a pre-integrated dataset. Precomputed normalization and integration values were removed, and raw counts were further processed. Data was split by patient and filtered for cells with a mitochondrial read fraction below 20% and more than 200 expressed genes. Normalization was performed using the NormalizeData() function with default parameters. Integration was performed using Harmony, and clustering and UMAP were computed with a resolution of 0.2 and 25 dimensions; layers were subsequently joined. Cell-cycle scores were assigned using the Seurat CellCycleScoring function with default parameters.

An initial cell type annotation was performed based on canonical marker expression: B cells (MS4A1, CD19, CD79A, CD79B, BLNK), T cells (CD8A, GZMA, CD3E, CD4, IL7R), myeloid cells (CD68, LYZ, C1QA, CD14, CD1C, CLEC10A), oligodendrocytes (Oligo; S100B, TF, GPM6B, PLP1, PTGDS, MOG, MAG, MBP), plasmacytoid dendritic cells (plasmacytoid DC; CLEC4C, IL3RA, LILR4A), and conventional dendritic cells (cDC; CLEC9A, CADM1, BATF3, BTLA, THBD, IRF8, XCR1). The T and B cell clusters were subsetted for more detailed annotation. T cell data were split by patient, and a new Harmony integration and clustering were performed (15 dimensions, resolution 2), with the layers subsequently joined. Clusters were annotated based on canonical markers for CD4^+^ T cells (CD3E, CD4, IL7R), CD8^+^ T cells (CD8A, GZMA), and NK cells (NCAM1, KLRK1, FCGR3A). One cluster expressing both T and B cell markers (MS4A1, CD19, CD79A, CD79B, BLNK) was removed.

The ProjecTIL package v3.6.0^52^ was used to further annotate CD4^+^ and CD8^+^ T cell clusters using the ProjecTILs.classifier() function with filter.cells = TRUE and min.confidence = 0.8. Reference datasets sketched_CD4T_human_ref_v2 and sketched_CD8T_human_ref_v1 were used. Clusters were annotated based on the majority of cells assigned to a ProjecTILs reference subset and canonical marker expression for exhausted CD8^+^ T cells (HAVCR2, TIGIT, CTLA4, LAG3, PDCD1), CD8^+^ central memory (CM) T cells, CD8^+^ effector memory (EM) T cells, CD8^+^ terminally differentiated effector memory (TEMRA) cells, CD4^+^ regulatory T cells (FOXP3, IL2RA), CD4^+^ T follicular helper (Tfh) cells, CD4^+^ naïve T cells, and CD4^+^ memory T cells. T cells were further marked as cycling based on the Seurat CellCycleScoring function assignment in S or G2/M phase.

B cell data were similarly split by patient, and a new Harmony integration and clustering were performed (15 dimensions, resolution 0.4), with the layers subsequently joined. One cluster expressing both B and T cell markers (CD8A, CD4, IL7R, CD3E) was removed. Clusters were annotated as healthy or malignant based on several criteria: (1) high number of unique genes expressed in malignant B cells, (2) high proportion of cells in S or G2/M phase in malignant B cells, (3) high expression of CD27 and CD38 in malignant B cells, with high expression of CD52, SCIMP, and BANK1 in healthy cells, (4) uniform ratio of IGKC to IGLC2 expression in healthy cells, calculated by assigning each cell as IGKC-expressing if IGKC/(IGKC+IGLC2) > 0.5, or IGLC2-expressing otherwise, then calculating cluster-level ratios, and (5) representation of healthy rLN cells in each cluster. All B cells from rLN samples were annotated as healthy. B cells were additionally marked as cycling based on Seurat CellCycleScoring assignment to S or G2/M phase. Finally, the detailed T and B cell annotations were reassigned to the full Seurat object containing all cell types. Gene signature scores from c5.go.bp.v2025.1.Hs.symbols.gmt and c2.cp.wikipathways.v2025.1.Hs.symbols.gmt were downloaded from https://www.gsea-msigdb.org/gsea/msigdb/human/genesets.jsp and added to the Seurat object using ClusterProfiler v4.14.6 and Escape v2.2.3/Ucell v2.10.1 with default parameters.

## QUANTIFICATION AND STATISTICAL ANALYSIS

### Statistical analysis

Unless otherwise specified in the figure legends, data are presented as mean ± s.e.m. Statistical analyses were performed in GraphPad Prism software (GraphPad, version 10). For two independent groups, two-tailed unpaired Student’s *t* tests or a Mann-Whitney U test (Wilcoxon rank-sum test) were used. All other statistical tests are specified in the corresponding figure legends. Exact *n* per experiment (biological replicates and independent experiments) and specific statistical tests are provided in the corresponding figure legends. Statistical significance is indicated in all figures by their *p*-values.

## SUPPLEMENTARY INFORMATION: FIGURE TITLES AND LEGENDS

**Extended Data Fig. 1.**
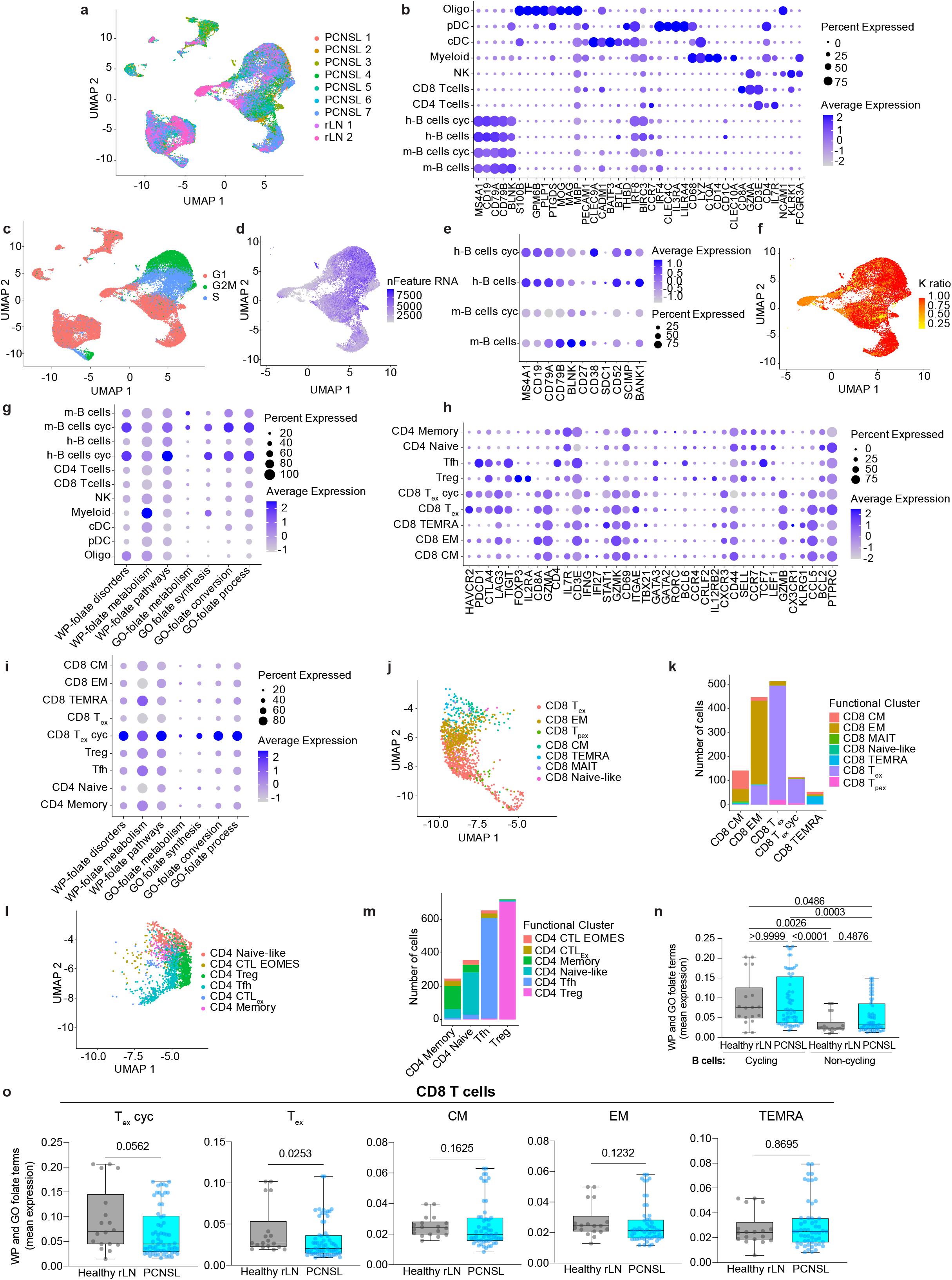
Cell type annotation and expression of folate metabolism gene signatures in human PCNSL scRNA-seq data. **a**, **b**, UMAP of integrated scRNA-seq data from PCNSL patient brain biopsies (*n* = 7) and reactive lymph node biopsies from patients without cancer (rLN; *n* = 2), representing all cell types. **a**, UMAP colored by patient samples. **b**, Percentage of positive cells and average expression for a selection of canonical cell type markers across cell type clusters from **Fig. 1a. c**, UMAP colored by cell cycle phase. **d**, UMAP of B cells only, colored by the number of unique genes expressed. **e**, Percentage of positive cells and average expression of a selection of genes with high and low expression among malignant and healthy B cell subsets. **f**, UMAP of B cells only, colored by the IGKC fraction (IGKC/(IGKC + IGLC2)) (immunoglobulin kappa constant, IGKC; immunoglobulin lambda constant 2, IGLC2), as a malignant B cell marker. **g,** Percentage of positive cells and average expression for WikiPathways signatures (WP) and Biological Processes Gene Ontology terms (GO) related to folate metabolism across cell types. Full signature names from left to right: WP – Disorders of folate metabolism and transport, WP – Folate metabolism, WP – Pathways into methionine and folate-cycles, GO – tetrahydrofolate metabolic process, GO – tetrahydrofolate biosynthetic process, GO – tetrahydrofolate interconversion, GO – 10-formyltetrahydrofolate metabolic process). **h**, Percentage of positive cells and average expression for a selection of canonical T cell subset markers across T cell clusters from Fig. 1e. **i**, Percentage of positive cells and average expression for WP signatures and GO terms related to folate metabolism across T cell subsets from Fig. 1e. **j**, Overlay of UMAP for CD8^+^ T cells only from **Fig. 1e** by ProjecTILs assigned CD8^+^ T cell clusters. **k**, Number of cells assigned with high confidence to a ProjecTILs reference CD8^+^ T cell subset. CM, central memory; EM, effector memory; MAIT, mucosal-associated invariant T cell; TEMRA, terminally differentiated effector memory; TEX, terminally exhausted; TPEX, progenitor exhausted. **l**, Overlay of UMAP for CD4^+^ T cells only from Fig. 1e by ProjecTILs assigned CD4^+^ T cell clusters. **m**, Number of cells assigned with high confidence to a ProjecTILs reference CD4^+^ T cell subset. Tfh, T follicular helper; Treg, regulatory T cells; CTL EOMES, cytotoxic CD4^+^ T cells expressing eomesodermin (Eomes) and Granzyme K (GZMK); CTL Exh, exhausted cytotoxic CD4^+^ T cells. **n, o,** Mean expression of each folate-related GO term and WP signature across PCNSL patients (*n* = 7) and rLN samples (*n* = 2). **n**, Proliferating (cycling) and non-proliferating (non-cycling) B cells. **o**, CD8^+^ T cell subsets. T_ex_ cyc; exhausted and proliferating/cycling CD8^+^ T cells; T_ex_, exhausted CD8^+^ T cells; CM, central memory; EM, effector memory; TEMRA, terminally differentiated effector memory. Box and whisker plots represent the median and interquartile range (IQR), whiskers represent the data min and max. Dots represent individual signatures in each patient (**n, o**). Statistical significance was assessed using either the Kruskal-Wallis test with Dunn’s multiple comparisons post-hoc test (**n**) or the Mann–Whitney U test (Wilcoxon rank-sum test) (**o**).

**Extended Data Fig. 2.**
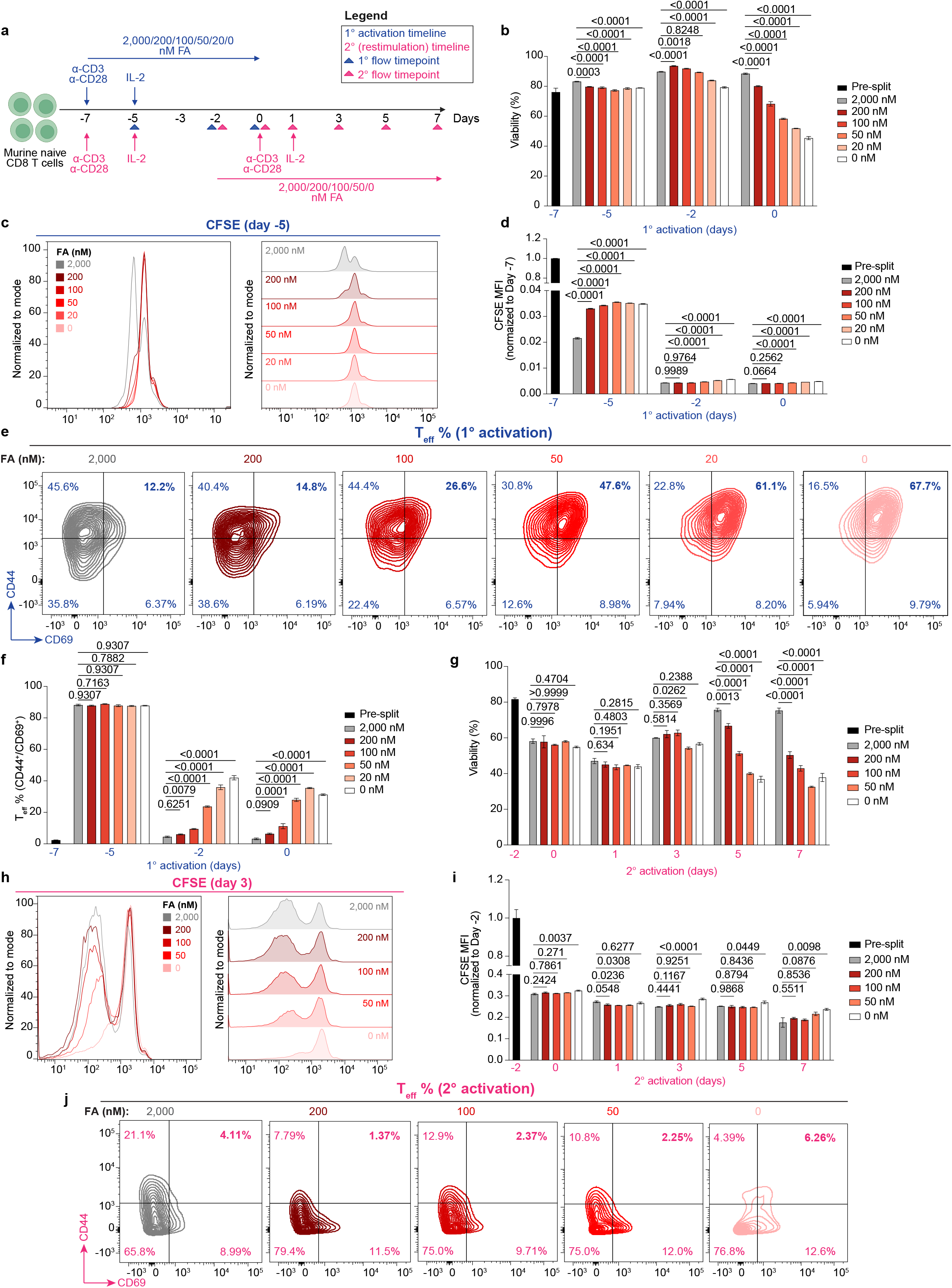
Proliferation, viability, and activation phenotypes of primary activated vs. restimulated folate-deprived CD8^+^ T cells. **a,** Experimental schematic of *ex vivo* primary activation and restimulation assays with various levels of folate (FA) deprivation. 2,000 nM FA represents the level of FA in RPMI media. Primary activation timeline is in blue, while the restimulation timeline is in pink. **b–f,** Viability (**b**), proliferation by CFSE dilution assay at day −5, **c, d**), and activation (**e**) for primary activated, FA-deprived CD8^+^ T cells. Activation in **e** is shown as representative flow cytometry plots for CD44 vs. CD69 expression on control (2,000 nM) and FA-deprived (200, 100, 50, 20, and 0 nM) CD8^+^ T cells on day −2. **f,** T_eff_ percentage (CD44^+^/CD69^+^) for all primary activation timepoints. **g–j,** Viability (**g**), proliferation by CFSE dilution assay on day 3 (**h, i**), and activation (**j**) for restimulated, FA-deprived CD8^+^ T cells. Activation data in **j** is shown as representative flow cytometry plots for CD44 vs. CD69 expression on control (2,000 nM) and FA-deprived (200, 100, 50, and 0 nM) CD8^+^ T cells on day 3. Data represent 3 independent experiments, and are presented as mean with s.e.m. Statistical significance was determined by one-way analysis of variance (ANOVA) with Dunnett’s multiple comparisons test.

**Extended Data Fig. 3.**
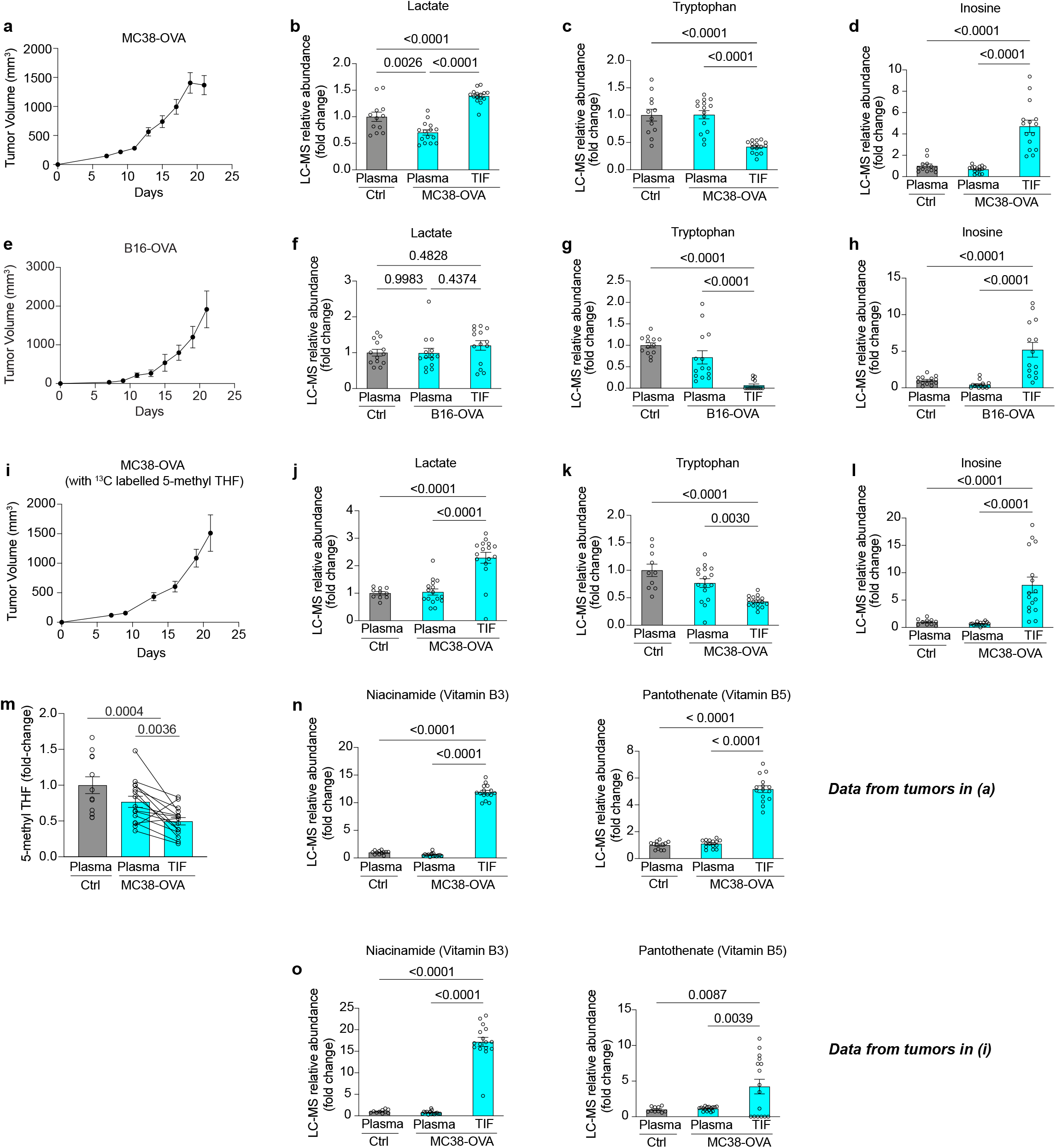
Metabolite profiling of MC38-OVA and B16-OVA tumor interstitial fluid. **a**, MC38-OVA tumor progression in mice (*n* = 16). **b–d**, Lactate (**b**), tryptophan (**c**), and inosine (**d**) levels in control plasma (*n* = 10), MC38-OVA plasma, and MC38-OVA tumor interstitial fluid (TIF) (*n* = 16). Fold change calculated relative to control plasma average. **e**, B16-OVA tumor progression in mice (*n* = 14). **f–h**, Lactate (**f**), tryptophan (**g**), and inosine (**h**) levels in control plasma (*n* = 13), B16-OVA plasma, and B16-OVA TIF (*n* = 14). Fold change calculated relative to control plasma average. **i**, MC38-OVA tumor progression in mice (*n* = 16). ^13^C-labelled 5-methyl THF standard curves in plasma and TIF were created for absolute quantification. **j–l**, Lactate (**j**), tryptophan (**k**), and inosine (**l**) levels in control plasma (*n* = 11), MC38-OVA plasma, and MC38-OVA TIF (*n* = 16). Fold change calculated relative to control plasma average. **m–o**, B-vitamin levels in plasma and TIF. **m**, 5-methyl THF levels (fold change calculated relative to control plasma) in plasma and TIF of control (*n* = 10) and MC38-OVA mice (*n* = 16) from (**a**). For **n, o**, niacinamide (vitamin B3) is on the left, pantothenate (vitamin B5) is on the right. Data is from MC38-OVA mice from both experiments presented here in **Extended Data Fig. 3a** (**n**), and **Extended Data Fig. 3i** (**o**). All data are representative of three independent experiments and are presented as mean and s.e.m. *P* values were determined by one-way analysis of variance (ANOVA) with Tukey’s post-hoc test for multiple comparisons (**b–d**, **f–h**, **j–o**) or paired, two-tailed parametric *t*-test (**m**).

**Extended Data Fig. 4.**
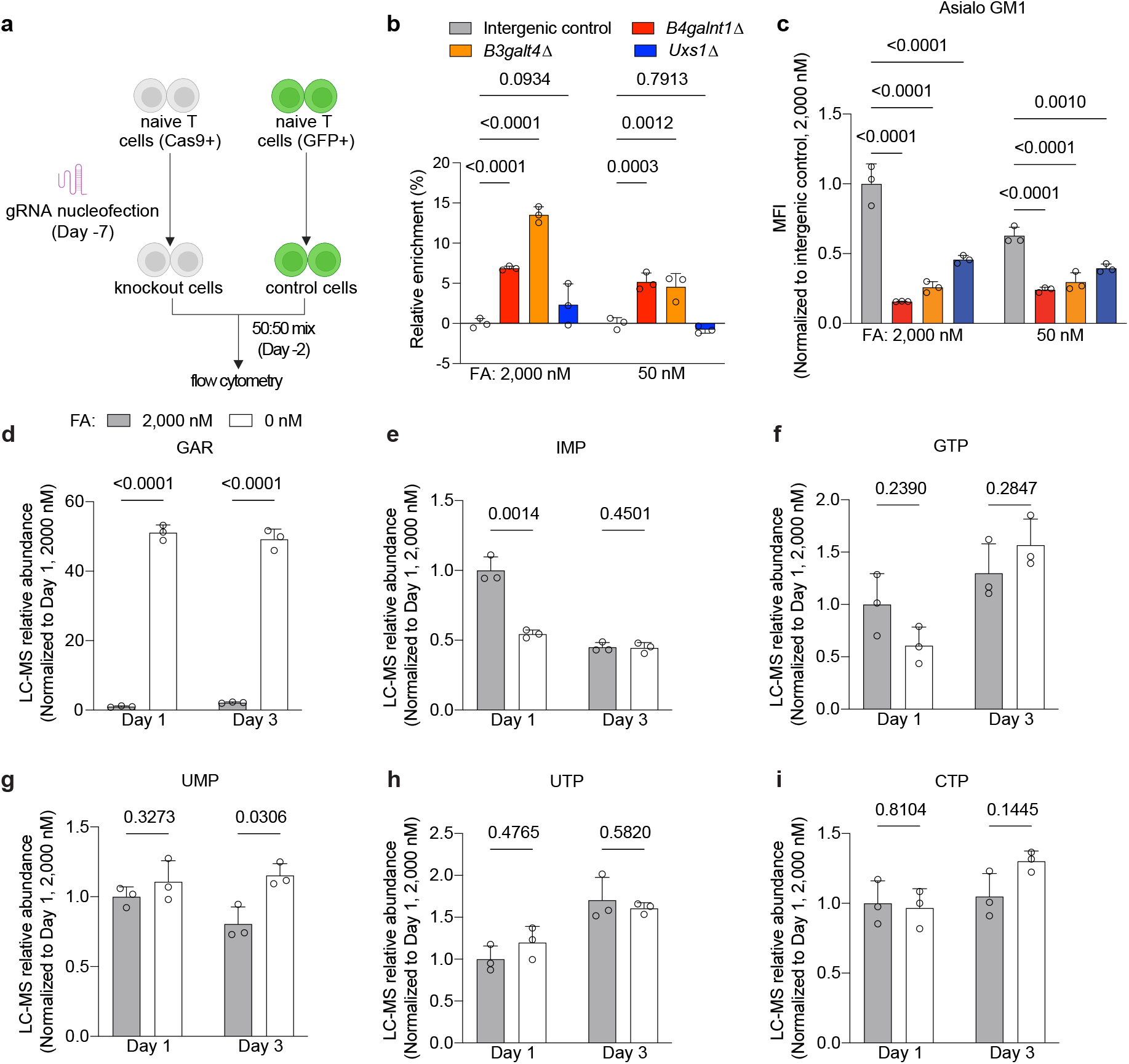
Folate-deprived CD8^+^ T cells’ metabolite profiling and gene essentiality. **a,** Experimental schematic of low FA CD8^+^ T cell CRISPR screen hits validation. Single gene knockouts (KO) of each hit were generated by nucleofecting Cas9^+^ CD8^+^ T cells at day −7. Single gene KO CD8^+^ T cells were mixed 50:50 with GFP^+^ CD8^+^ T cells at day −2 and split into high FA (2,000 nM) and low FA (50 nM). Competitive growth was measured by flow cytometry. **b**, Percentage of relative enrichment of CD8^+^ T cells with single gene KO in high FA (2,000 nM) and low FA (50 nM) at day 5, normalized to 2,000 nM FA intergenic control condition. **c**, Asialo-GM1 cell surface median fluorescent intensity (MFI) levels on CD8^+^ T cells with single gene KO in high FA (2,000 nM) and low FA (50 nM) at day 5, normalized to the 2,000 nM FA intergenic control condition. **d–i**, Relative abundance of purine intermediates GAR (**d**), IMP (**e**), and the nucleotides GTP (**f**), UMP (**g**), UTP (**h**), and CTP (**i**) in control (2,000 nM) vs. FA-deprived CD8^+^ T cells (0 nM) at days 1 and 3 after restimulation. GAR - Glycinamide ribonucleotide; IMP - inosine monophosphate; GTP - guanosine triphosphate; UMP - uridine monophosphate; UTP - uridine triphosphate; CTP - cytidine triphosphate. All data represent 3 biological replicates and are presented as mean and s.e.m. Statistical significance was determined by a repeated-measures two-way ANOVA with Šídák’s multiple comparisons test.

**Extended Data Fig. 5.**
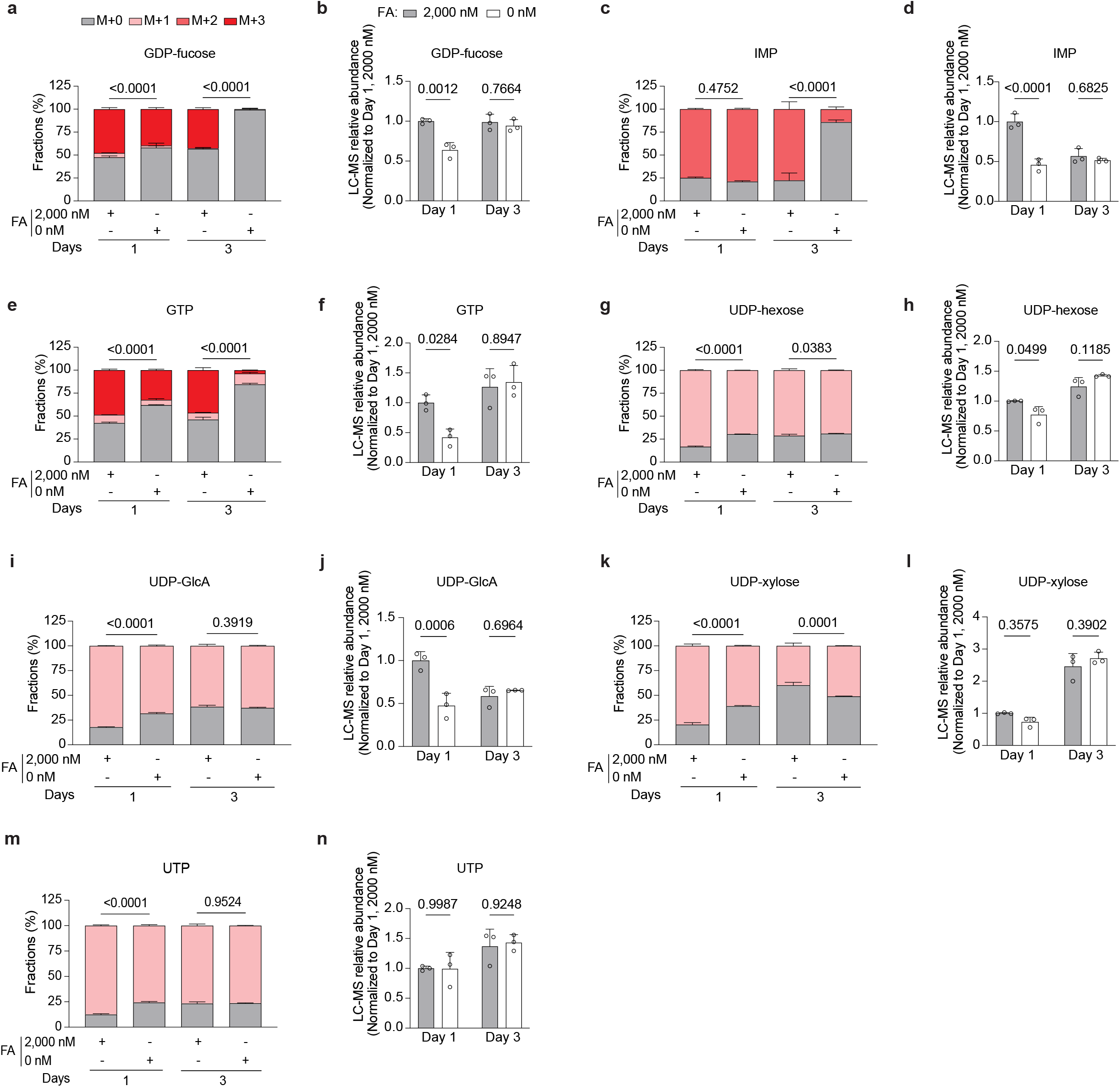
Purine and pyrimidine metabolism in folate-deprived CD8^+^ T cells. **a–n**, ^15^N glutamine labelling in control (2,000 nM) vs. FA-deprived CD8^+^ T cells (0 nM) at days 1 and 3 post-restimulation. Data includes fractional labeling and relative abundance of GDP-fucose (**a, b**), IMP (**c, d**), GTP (**e, f**), UDP-hexose (**g, h**), UDP-glucuronic acid (UDP-GlcA) (**i, j**), UDP-xylose (**k**, **l**), and UTP (**m, n**). For fractional labelling (**a, c, e, g, i, k, m**), M+0 represents the endogenous unlabelled metabolite (grey bars). *De novo* synthesis is represented by incorporation of the stable isotope, indicated by M+1, M+2, and M+3 (red bars); these mass isotopomer distributions are shown as fraction percentages. Relative abundance (**b, d, f, h, j, l, n**) is normalized to day 1 control condition (2,000 nM FA). All data represent 3 biological replicates and are presented as mean and s.e.m. *P* values were determined by repeated-measures two-way ANOVA. To evaluate the treatment effect at each time point, Šídák’s multiple comparisons test was used. (**a, c, e, g, i, k, m**; comparing the M+0 fraction of each condition) and (**b, d, f, h, j, l, n**).

**Extended Data Fig. 6.**
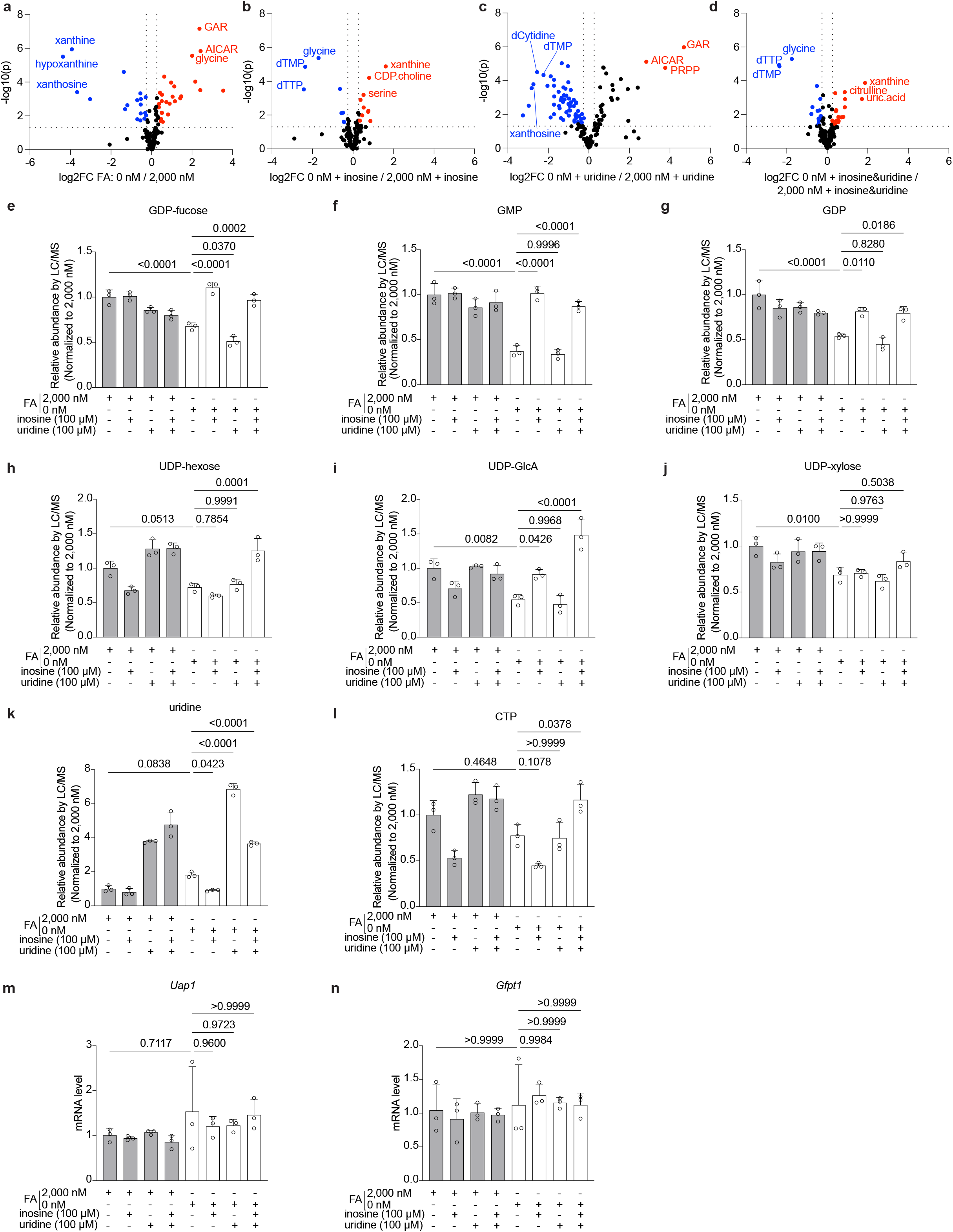
Inosine supplementation rescues nucleotide sugar metabolism defects in folate-deprived CD8^+^ T cells. **a–d**, Volcano plots of differentially depleted (blue dots) and enriched (red dots) metabolites in response to FA deprivation (0 nM FA vs 2,000 nM FA) (**a**) and with nucleoside supplementation with inosine (100 µM) (**b**), uridine (100 µM) (**c**), or both (**d**). **e–l**, Relative abundance of GDP-fucose (**e**), GMP (**f**), GDP (**g**), UDP-hexose (**h**), UDP-glucuronic acid (UDP-GlcA) (**i**), UDP-xylose (**j**), uridine (**k**), and CTP (**l**) in FA-deprived T cells supplemented with inosine (100 µM), uridine (100 µM), or both. Relative abundance is normalized to 2,000 nM FA only. **m, n,** mRNA transcript levels by qPCR of enzymes involved in nucleotide sugar metabolism and UDP-GlcNAc synthesis. **m**, UDP-N-Acetylglucosamine pyrophosphorylase 1 (*Uap1*). **n**, Glutamine fructose-6-phosphate transaminase 1 (*Gfpt1*). All data represent 3 biological replicates and are presented as mean and s.e.m. Statistical significance was determined by one-way ANOVA with Dunnett’s multiple comparisons test (**e–n**).

**Extended Data Fig. 7.**
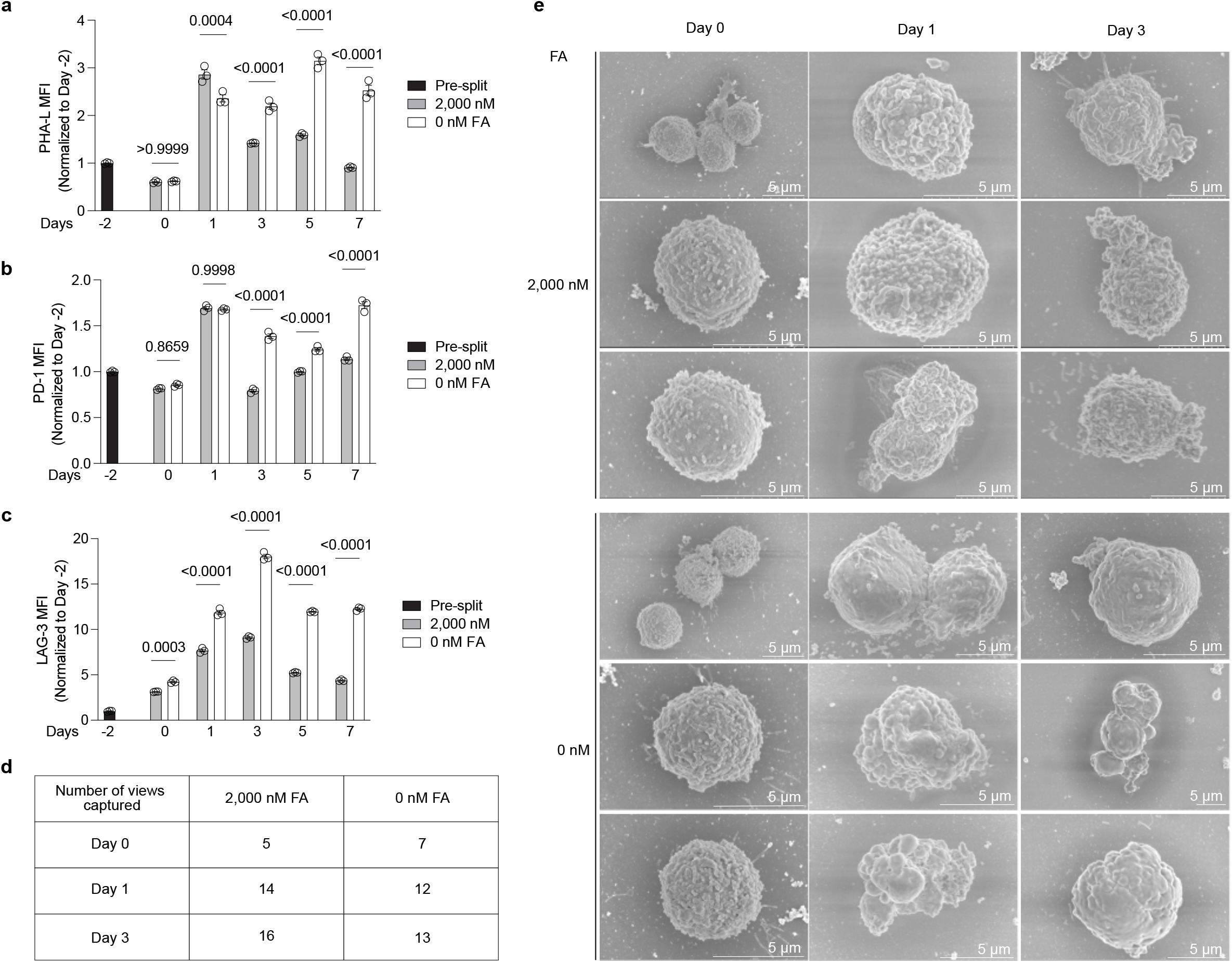
Reduced glycocalyx complexity and dysregulated cell surface exhaustion marker expression in folate-deprived CD8^+^ T cells. **a–c**, Median fluorescent intensity measured by flow cytometry of control (2,000 nM) vs. FA-deprived (0 nM) CD8^+^ T cells over time (days −2 pre restimulation to 7 days post-restimulation) of Phaseolus vulgaris Leucoagglutinin (PHA-L), an N-glycosylation cell surface marker (**a**), Programmed cell death protein 1 (PD-1) (**b**), and Lymphocyte-activation gene 3 (LAG-3) (**c**). **d**, Number of fields captured during scanning electron microscopy (SEM) for each condition. **e**, Additional representative SEM images of CD8^+^ T cell glycocalyx morphology in 2,000 nM vs. 0 nM FA at days 0, 1, and 3 post-restimulation. Scale bar, 5 µm. All data represent 3 biological replicates and are presented as mean and s.e.m. Statistical significance was determined by two-way ANOVA with Tukey’s post-hoc test for multiple comparisons.

**Extended Data Fig. 8.**
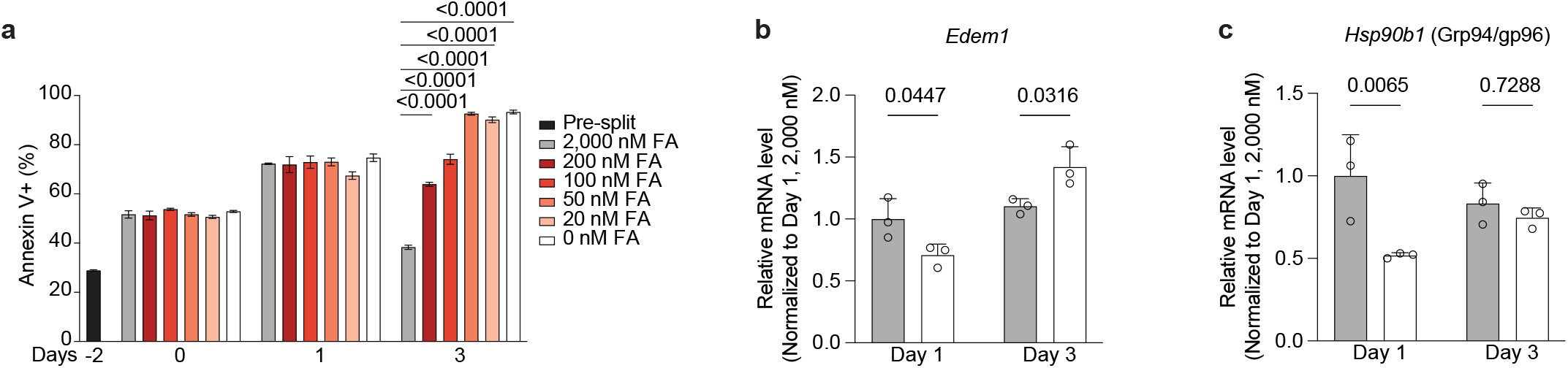
Folate deprivation triggers the unfolded protein response (UPR) and apoptosis in CD8^+^ T cells. **a**, Flow cytometry of the apoptotic marker Annexin V^+^ in CD8^+^ T cells in control (2,000 nM) vs. FA-deprived (200, 100, 50, 20, and 0 nM) conditions at day −2 (pre-split) and days 0, 1, and 3 after restimulation. **b**, **c**, qPCR quantification of the adaptive UPR markers Endoplasmic reticulum degradation enhancing alpha-mannosidase-like protein 1 (*Edem1*) (**b**), and Heat shock protein 90kDa beta member 1 (*Hsp90b1,* aka Grp94/Gp96) (**c**) in control and FA-deprived CD8^+^ T cells at days 1 and 3 post-restimulation. All data represent 3 biological replicates and are presented as mean and s.e.m. Statistical significance was determined by one-way ANOVA with Dunnett’s multiple comparisons test (**a**), or by a repeated-measures two-way ANOVA with Šídák’s multiple comparisons test to evaluate the treatment effect at each time point used (**b**, **c**).

**Extended Data Fig. 9.**
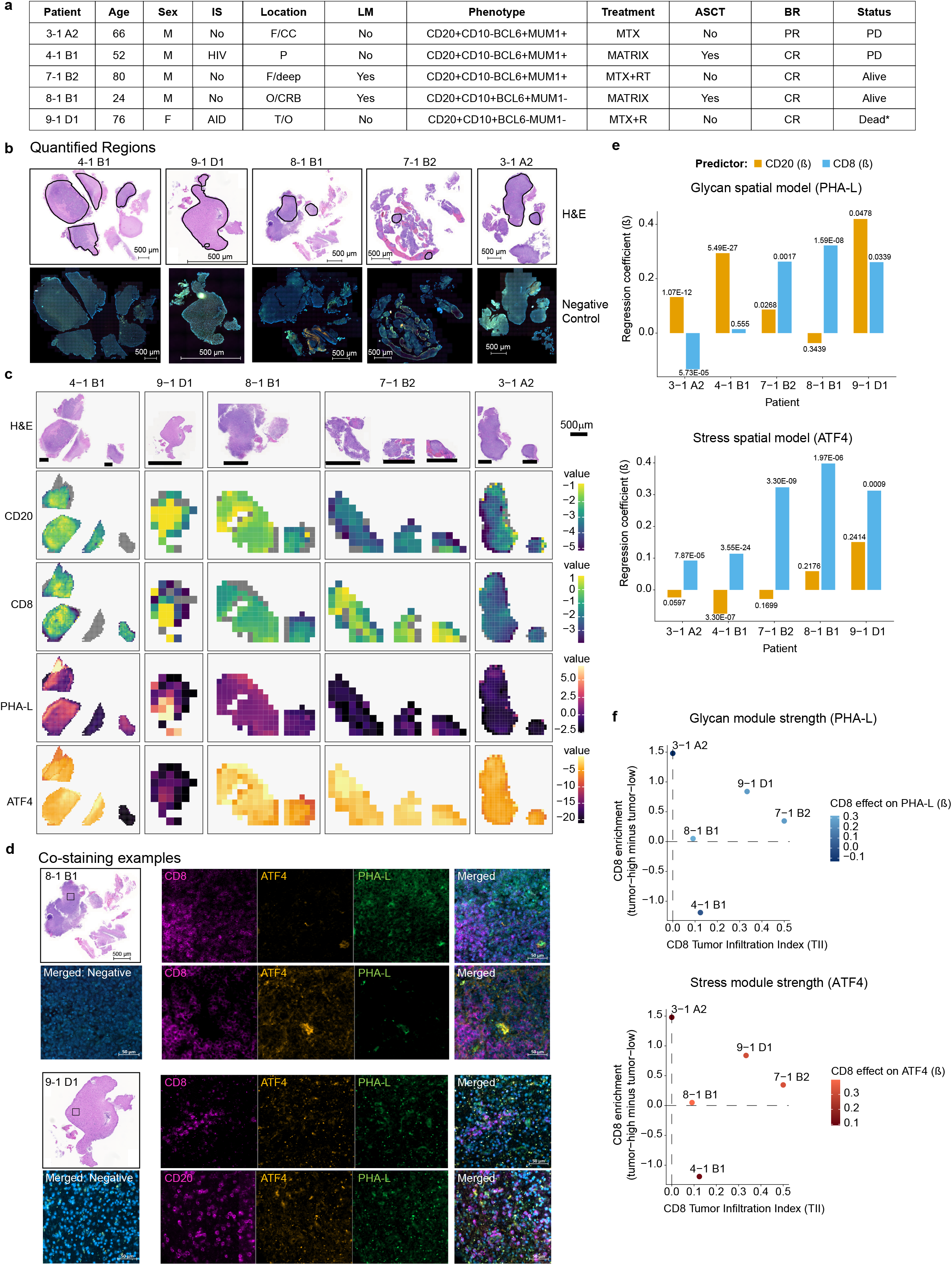
Independent spatial contributions of tumor and immune compartments to glycan and stress modules. **a,** Clinical and phenotypic characteristics of PCNSL patients (*n* = 5). IS, immunological status. F, frontal lobe; CC, corpus callosum; P, parietal lobe; deep, deep brain structures; O, occipital lobe; CRB, cerebellum; T, temporal lobe. LM, leptomeningeal involvement. MTX: high-dose methotrexate (3 g/m^2^); RT, radiotherapy (whole brain); R, rituximab (375 mg/m^2^); MATRIX, methotrexate, cytarabine, thiotepa, and rituximab regimen; ASCT, autologous stem cell transplantation. BR, best response; CR, complete response; PR, partial response; PD, progressive disease. *Patient 9-1 D1’s death unrelated to lymphoma progression (septic shock). **b**, Representative whole-section images of primary human PCNSL biopsies used for spatial analysis (*n* = 5). Top, H&E staining is shown for each patient with quantified regions for spatial analysis outlined in black (scale bar = 500 µm). Bottom, negative control channels for corresponding multiplexed immunofluorescence (IF) acquisition to illustrate the regions retained or excluded from spatial analysis based on background fluorescence levels across sections. **c**, IF staining of human PCNSL tissue sections (*n* = 5). Spatial IF phenotype maps of tumor cells (CD20), CD8 T cells (CD8), branched N-glycans (PHA-L), and levels of the stress responsive transcription factor, ATF4, from human PCNSL sections (square tiles, 300 µm x 300 µm). Color scales indicate relative marker intensity per bin (see color bars). Representative hematoxylin and eosin (H&E) sections are shown for each tumor (top row) to illustrate tissue morphology and overall architecture. Scale bar = 500 µm. **d**, Representative IF co-staining examples from 2 PCNSL patients for CD8 (purple), CD20 (purple), ATF4 (yellow), PHA-L (green), and DAPI (blue); scale bar = 50 µm. The displayed IF images correspond to the boxed areas indicated on the H&E images (left; scale bar = 500 µm). Merged negative control images (left) and merged fluorescent channel images (right) are provided. **e**, Multivariable spatial regression coefficients (β) derived from 400 µm bin-level models fitted separately for each tumor. For each section, PHA-L (glycan module) and ATF4 (stress module) signals were modeled as a function of CD20 (tumor) and CD8 densities. Bars represent β coefficients corresponding to the independent contribution of CD20 (gold) and CD8 (blue). Statistical significance of β coefficients was determined by standard Wald tests from linear regression models. **f**, Projection of glycan (top) and stress (bottom) module-associated CD8 effects onto immune-tumor architectural space. The CD8 Tumor Infiltration Index (TII) is plotted against CD8 enrichment (tumor-high minus tumor-low bins), with point color representing the magnitude of the CD8-associated module effect (β).

**Extended Data Fig. 10.**
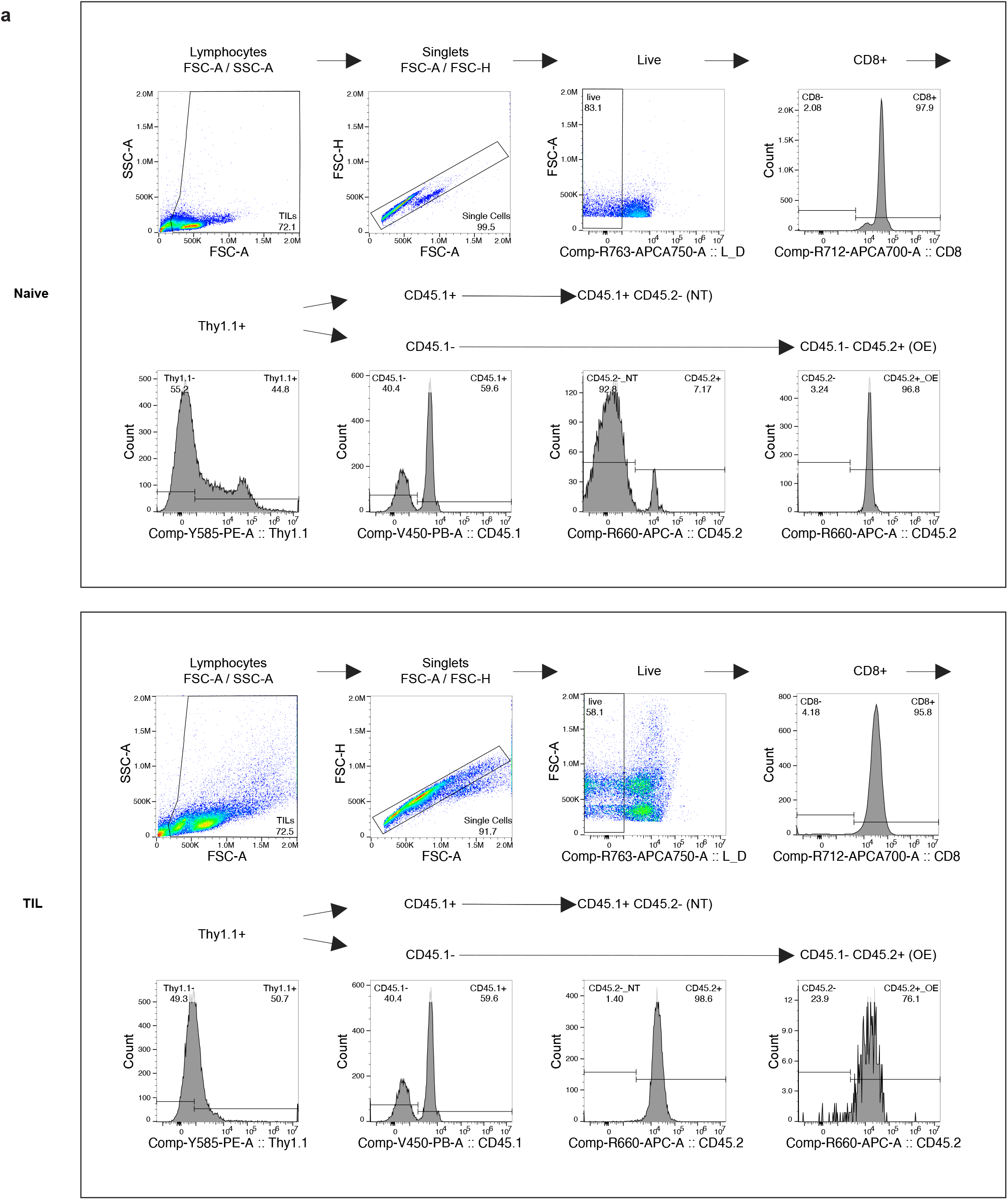
Flow cytometry gating strategy for *Slc19a1* OE/*Slc46a1* OE vs. NT CD8 TIL phenotyping. **a,** Representative flow cytometry plots demonstrating the gating strategy used to quantify CD45.1^+^/CD45.2^-^ (non-targeting control, NT) and CD45.1^-^/CD45.2^+^ (*Slc19a1* overexpressing (OE) or *Slc46a1* OE) OT-I cells from **Fig. 7**. Examples for naive cells (top) and tumor infiltrating lymphocytes (TIL, bottom) are shown.

